# Structural and mechanistic insights into bacterial hydrazine biosynthesis

**DOI:** 10.1101/2025.08.22.671884

**Authors:** Guiyun Zhao, Huisi Huang, Yifan Li, Jian Yang, Yuan-Yang Guo, Liqiao Xu, Lu Wang, Jingkun Shi, Miaolian Wu, Yu Feng, Binju Wang, Zhi-Min Zhang, Yi-Ling Du

**Affiliations:** Department of Microbiology and The Fourth Affiliated Hospital, Zhejiang University School of Medicine, Hangzhou, China; State Key Laboratory of Bioactive Molecules and Drugability Assessment, School of Pharmacy, Jinan University, Guangzhou 511436 P. R. China; State Key Laboratory of Physical Chemistry of Solid Surfaces and Fujian Provincial Key Laboratory of Theoretical and Computational Chemistry, College of Chemistry and Chemical Engineering, Xiamen University, 361005, Xiamen, China; State Key Laboratory of Antiviral Drugs, Henan Normal University, Xinxiang, Henan 453007, China; Department of Biophysics, Zhejiang University School of Medicine, Hangzhou 310058, China; Jinan Microecological Biomedicine Shandong Laboratory, Jinan, China

## Abstract

Nitrogen-nitrogen (N-N) bond-containing motifs are prevalent in both clinical drugs and natural products. Bacterial hydrazine synthetases catalyze N-N bond formation by coupling an amino acid and a hydroxylamine via a distinctive *O*-aminoacyl-hydroxylamine intermediate. Despite its wide occurrence, the structural and mechanistic basis of this process has remained elusive. Here, we report the first crystal structures of the *O*-aminoacyl-hydroxylamine synthetase component of a bacterial hydrazine synthetase, captured in binary and ternary complexes with substrates and catalytic intermediates. These structures reveal the molecular determinants of substrate recognition and, together with biochemical and computational analyses, establish a detailed mechanistic framework for the hydrazine synthetase family. Leveraging these insights, we expanded hydrazine biosynthesis through the targeted discovery of novel hydrazine synthetases and implemented a chemoenzymatic synthesis strategy. This work provides fundamental insights into hydrazine synthetases and lays the groundwork for their rational engineering as versatile biocatalysts.

## Introduction

Nitrogen-nitrogen (N-N) bonds are prevalent structural motifs found in numerous synthetic drugs and natural products^1,2^. Functional groups such as hydrazine, diazo, pyrazole, N-nitroso, hydrazide, triazene, and hydrazone play critical roles in shaping the bioactivity and chemical properties of their parent molecules. Owing to their importance, considerable effort has been devoted to developing diverse synthetic strategies for N-N bond construction^2–6^. In parallel, elucidating how nature builds N-N bonds and identifying the enzymes responsible for these transformations provide exciting opportunities to harness biocatalysts for producing N-N-containing chemicals and pharmaceuticals. Notably, only in recent years have dedicated N-N bond-forming enzymes begun to be uncovered^7–15^.

A conserved biosynthetic strategy for intermolecular N-N bond formation has emerged as a central mechanism for generating hydrazine precursors in diverse bacterial metabolites (Fig. 1)^8,16–25^. The key step involves the coupling of an amino acid (or an aminoacyl-tRNA) with a hydroxylamine to generate an unusual *O*-aminoacyl-hydroxylamine intermediate, a species not observed in other biosynthetic processes (Fig. 1a). This intermediate can then undergo intramolecular rearrangement to form an N-N bond. Distinct enzyme families have independently adopted this strategy^14,15,26^, with bacterial hydrazine synthetases representing the largest and most widespread N-N bond-forming enzyme family described to date^8,16–25^. An expanding set of N-N-containing natural products has now been shown to depend on hydrazine synthetases for metabolic precursors formation (Fig. 1b).

**Fig. 1.**
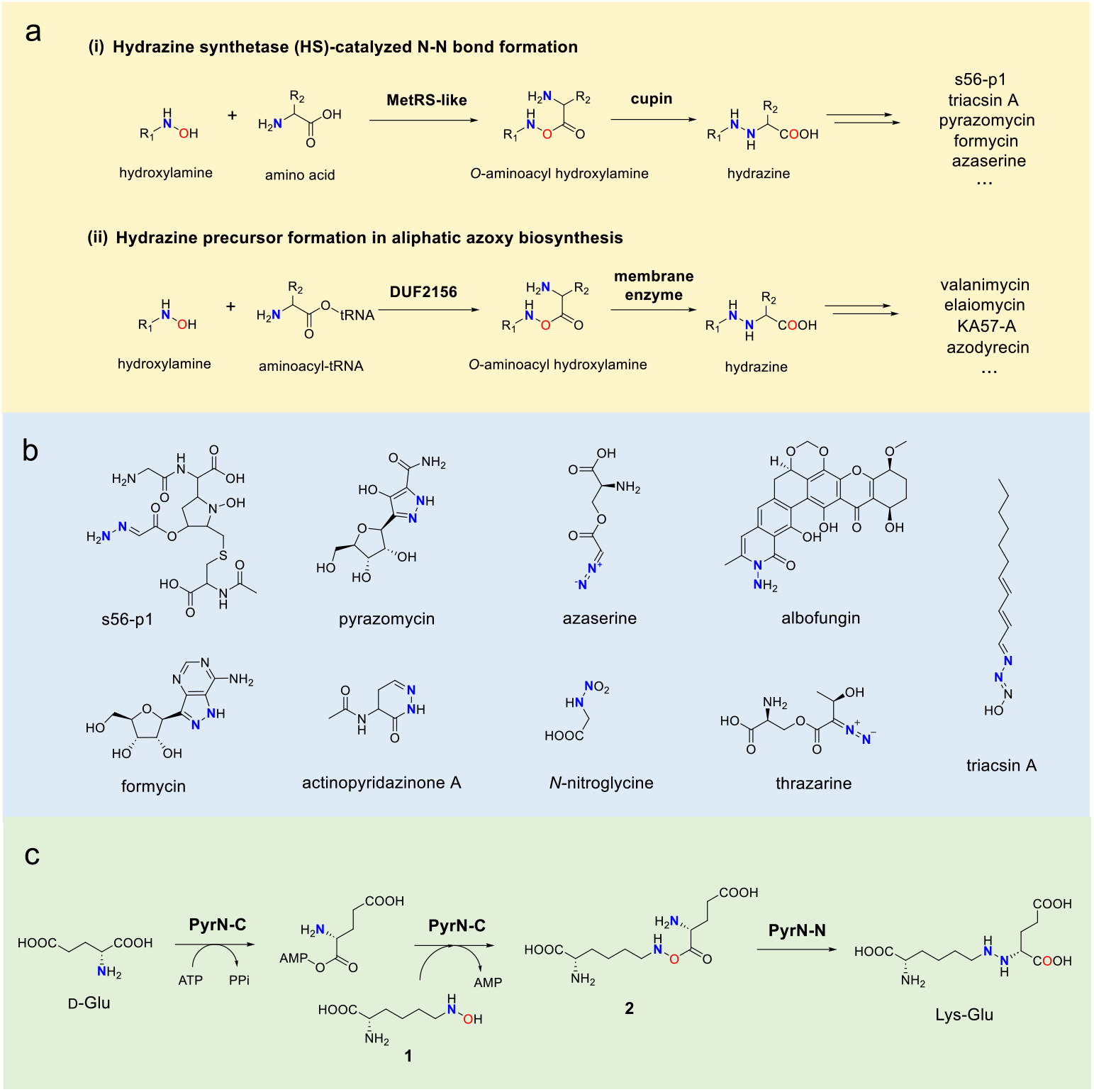
A conserved enzymatic strategy for generating hydrazine precursors in bacterial natural products with diverse N-N moieties. (a) Reaction schemes for enzyme families that employ this biosynthetic strategy, characterized by the formation of a distinctive *O*-aminoacyl hydroxylamine intermediate. (b) Representative N-N-containing natural products whose biosynthesis relies on hydrazine synthetases for N-N bond formation. (c) The multi-step N-N-forming reaction catalyzed by hydrazine synthetases (exemplified by PyrN), as revealed by this study and previous work.

Didomain hydrazine synthetases typically comprise a small N-terminal cupin domain (∼100 residues) and a large C-terminal domain (∼550 residues) homologous to class I aminoacyl-tRNA synthetases such as methionyl-tRNA synthetase (MetRS)^8,12,13^. Depending on the biosynthetic context, these domains may exist as separate proteins or as components of a multidomain fusion enzyme. Members of this family catalyze intermolecular N-N bond formation between a free amino acid and an amino-acid-derived hydroxylamine (Fig. 1a). For example, in the biosynthesis of s56-p1 and triacsins, hydrazine synthetases Spb40/Tri28 couple glycine and *N*^6^-hydroxy-l-lysine (**1**)^8,13^, whereas in the pyrazomycin pathway, PyrN conjugates glutamate to **1** (Supplementary Fig. 1)^12,27^. In vitro biochemical studies have shown that the two domains are functionally separable^12,13^. The large C-terminal MetRS-like domain is responsible for the generation of the key *O*-aminoacyl-hydroxylamine intermediate, whereas the small N-terminal cupin domain mediates the subsequent intramolecular rearrangement to form a N-N bond (Fig. 1a). Bioinformatic analyses indicate that hydrazine synthetase are widely distributed across bacterial genomes^8,12,28^.

The discovery of hydrazine synthetases, which assemble N-N bonds from simple building blocks, highlights their potential as versatile biocatalysts. However, despite their broad distribution and catalytic promise, the structural and mechanistic basis of hydrazine synthetases—particularly their large MetRS-like domain—remains poorly understood, limiting both mechanistic insight and biocatalyst engineering. In this study, we integrate structural, computational, and biochemical approaches to elucidate the catalytic mechanism of PyrN, the hydrazine synthetase from the pyrazomycin pathway. Building on these insights, we expand the hydrazine synthetase family through structure-guided mining and further demonstrate the chemoenzymatic synthesis of non-natural hydrazine-containing compounds. Together, these findings provide a comprehensive view of the structural and catalytic principles governing bacterial hydrazine synthetases.

## Results

### Structural insights into the hydrazine synthetase PyrN

Although hydrazine synthetases have been identified from many biosynthetic pathways, no crystal structure of a didomain hydrazine synthetase or its large C-terminal domain has been reported. As a result, the structural and mechanistic basis of bacterial hydrazine biosynthesis—particularly the generation of the key *O*-aminoacyl– hydroxylamine intermediate—has remained elusive. To address this gap, we attempted to crystallize PyrN from the pyrazomycin biosynthetic pathway. Despite extensive crystallization trials, we were unable to obtain structures of full-length PyrN in either apo or substrate-bound states. Given our previous detailed characterization of RHS1 (a standalone PyrN-N homolog)^12^, we instead focused on the large C-terminal domain (PyrN-C) (Supplementary Fig. 2). PyrN-C catalyzes the ATP-dependent transfer of glutamate to the *N*^6^-OH group of **1**, generating the highly unstable ester intermediate *O*-glutamyl-*N*^6^-hydroxyl-lysine (**2**), setting ready for subsequent PyrN-N-catalyzed N-N bond formation (Fig. 1c). It is worth to mention that initial in vitro reconstitution assays employed l-glutamate (l-Glu) as the substrate; however, a recent study demonstrated d-glutamate (d-Glu) to be the physiological substrate^12,27^. The earlier observation of l-Glu activity was likely an artifact resulting from the high substrate concentrations (20 mM) used in those assays.

Based on prior findings, we hypothesized that a glutamyl-adenylate intermediate is formed during catalysis and sought to capture its bound state. To prevent hydrolysis, we first solved the crystal structure of PyrN-C in complex with l-glutamyl-sulfamoyl-adenosine (GSA), a non-hydrolysable analog of l-glutamyl-adenylate (Fig. 2a-2b and Supplementary Fig. 3). The resulting dimeric structure was determined at 2.23 Å resolution (PDB: 9L8V) (Supplementary Fig. 4 and Supplementary Table 2). Interestingly, we later found that co-crystallization of PyrN-C with either d-or l-Glu and Mg-ATP (ratio 1:2:10) yielded structures with clear electron density for the corresponding d-or l-glutamyl-adenylate (DGA/LGA) intermediates (Fig. 2c-2d). These complexes were determined at 2.00 Å (PDB: 9VQJ) and 2.10 Å (PDB: 9VQ4) resolution, respectively (Supplementary Table 3). Superposition of these binary complexes revealed that both stereoisomers bind within the glutamate pocket without inducing significant conformational changes in active site residues (Supplementary Fig. 5).

**Fig. 2.**
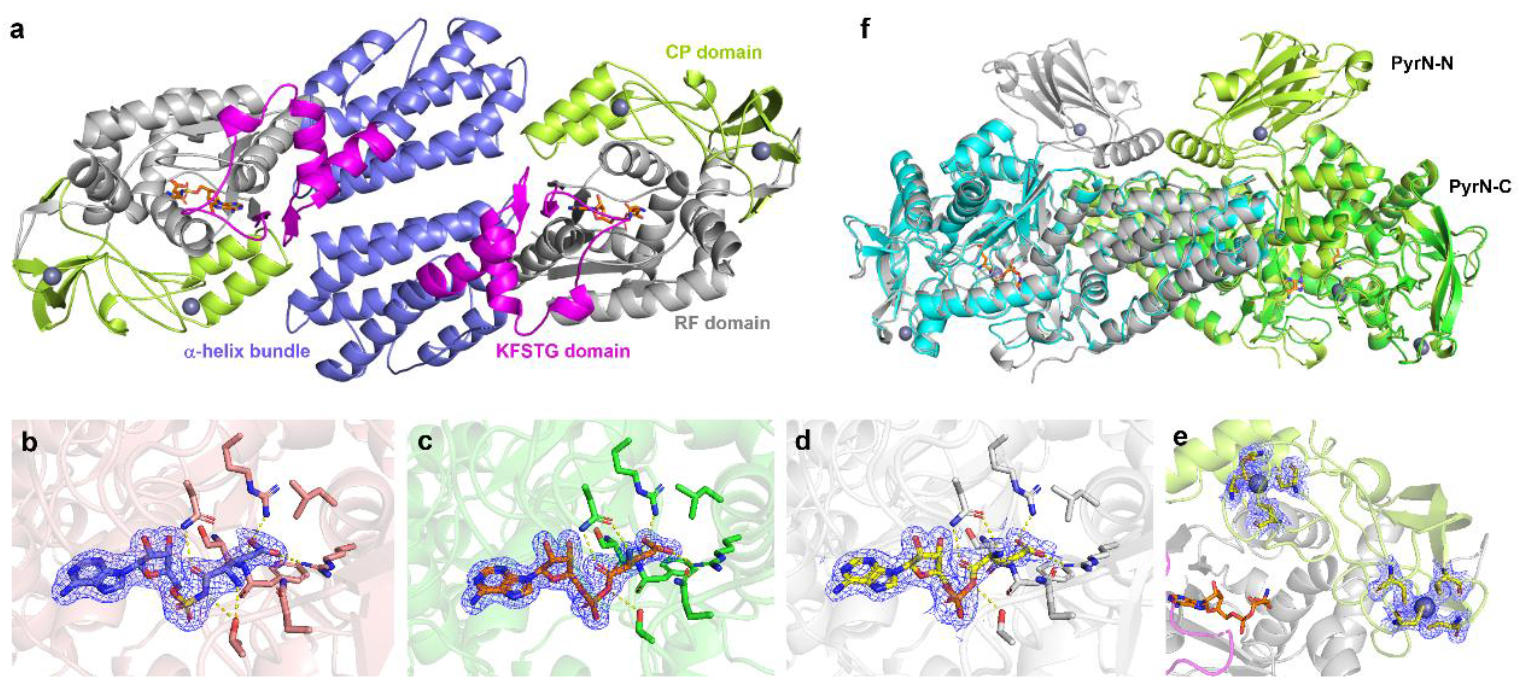
Structural studies of PyrN-C. (a) Structural organization of the co-crystal structure of PyrN-C in complex with GSA. PyrN-C features a Rossmann-fold catalytic core (gray) with an inserted connective peptide (CP) domain (lemon), the KFSTG motif-containing domain (magenta), and an α-helix bundle (marine). GSA is represented as orange stick, and zinc ions are shown as gray spheres. (b) The 2Fo-Fc electron density maps contoured to 1 σ for GSA. The blue mesh represents electron density. (c) The 2Fo-Fc electron density maps contoured to 1 σ for DGA. (d) The 2Fo-Fc electron density maps contoured to 1 σ for LGA. (e) The 2Fo-Fc electron density maps contoured to 1 σ for zinc ions and coordinating cysteine residues. (f) Superposition of PyrN-C dimer with the AlphaFold-predicted dimer structure of full-length PyrN. The crystal structure of PyrN-C is shown in green and cyan, whereas the predicted structure of PyrN is displayed in lemon and gray.

A Dali server search identified the closest structural homologs of PyrN-C as class I aminoacyl-tRNA synthetases (aaRSs).^29^ These include methionyl-tRNA synthetases (MetRSs) from *Pyrococcus abyssi* (PaMetRS; 22% sequence identity) and *Escherichia coli* (EcMetRS; 16%) (Supplementary Fig. 6),^30,31^ the isoleucyl-tRNA synthetase 2 from *Priestia megaterium* (PmIleRS2; 16%),^32^ and arginyl-tRNA synthetase from *Thermus thermophilus* (TtArgRS; 16%).^33^ Similar to the core architecture of EcMetRS^31^, PyrN-C consists of four structural domains: a Rossmann-fold (RF) catalytic core containing an inserted connective peptide (CP) domain, a domain harboring the KFSTG-motif (corresponding to the ‘KMSKS’ motif in canonical MetRSs), and an α-helix bundle (Fig. 2a). The CP domain, positioned above the catalytic core, contains two zinc-coordinating clusters: Cys256, Cys259, Cys302, Cys305, and Cys272, Cys275, Cys285, Cys288 (Fig. 2e). While canonical MetRSs exhibit variable zinc content across organisms^34^, structural alignment of PyrN-C with homologous domains from other hydrazine synthetases suggests that the number of zinc knuckles also varies within this enzyme family (see later sections).

To have a structural view of the didomain PyrN, we utilized single-particle cryo-electron microscopy (cryo-EM) to determine the structure of full-length PyrN in complex with GSA (Supplementary Fig. 7 and Fig. 8). Non-uniform refinement resulted in a reconstruction at 3.23 Å resolution. While the C-terminal domain is well-resolved with unambiguous density, no clear density is observed for the N-terminal domain. A plausible explanation is that the N-terminal domain is flexibly tethered to the C-terminal domain. Since the molecular mass of the C-terminal domain is larger than that of the N-terminal domain, particle alignment is primarily driven by the C-terminal domain, causing the signal of the N-terminal domain to be averaged out during data processing. We then generated an AlphaFold-predicted model of the didomain enzyme (Fig. 2f and Supplementary Fig. 9)^35^. This model shows that the small N-terminal cupin domain (PyrN-N) is linked to PyrN-C via a α-helix segment spanning residues 111-126.

### Identification of key residues responsible for amino acid substate recognition

Co-crystal structures of PyrN-C bound to GSA, DGA, or LGA reveal that ligand-interacting residues are exclusively located within the Rossmann-fold catalytic core and the KFSTG-motif-containing domain (Fig. 2a). In the PyrN-C-DGA co-crystal structure, the adenosine moiety of DGA is accommodated within a pocket formed by the ^147^HLGH^150^ and ^458^KFSTG^462^ motifs, which correspond to the HIGH and KMSKS motifs of class I aaRSs (Fig. 3a)^31^. The adenine base is stabilized through hydrogen bonds with the side-chain amide of Asn449 and the main-chain atoms of Tyr452 and Phe459. The ribose hydroxyl groups interact with Ser138, Gly418, Asp420, and Asn421. The α-amino and α-carboxyl groups of the glutamyl moiety form hydrogen bonds with Asn421 and water-mediated interactions involving Thr141, Gln182, and Asp420. The γ-carboxylate group engages in direct interactions with the guanidinium side chains of Arg390 and Arg425, as well as the main chain of Phe139.

**Fig. 3.**
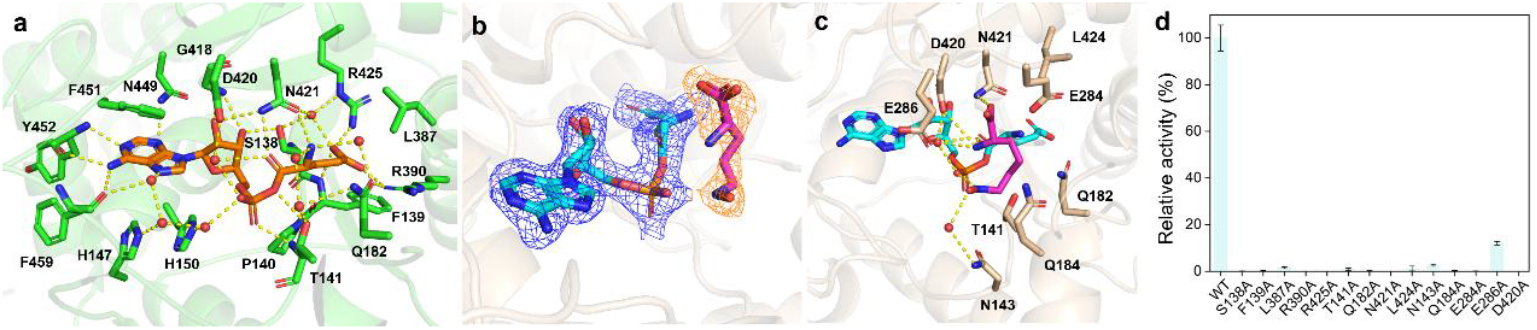
(a) Residues in the substrate-binding pocket that interact with DGA. Residue numbers are specific to the full-length PyrN. Polar interactions are shown as yellow dashed lines. Water molecules are shown as red spheres. (b) The 2Fo-Fc electron density maps contoured to 1 σ. The blue mesh and orange mesh represent electron density for LGA and **1**, respectively. LGA is shown in cyan stick, and *N*^6^-OH-l-Lys (**1**) is displayed as magenta stick. (c) The active site of the PyrN-C-LGA-**1** complex structure. (d) Site-directed mutagenesis and in vitro assays of selected active site residues of PyrN.

To examine the binding mode of another substrate *N*^6^-hydroxy-l-lysine (**1**), we soaked PyrN-C-LGA/DGA crystals with **1**. Clear electron density for **1** was observed only in PyrN-C-LGA complex (refined to 2.45 Å; PDB: 9VR2) (Fig. 3b), suggesting that **1** reacts rapidly with DGA upon binding. The ternary PyrN-C-LGA-**1** structure superimposes closely with the PyrN-C-LGA/DGA binary complexes, indicating no substantial conformational rearrangements upon **1** binding (Supplementary Fig. 10 and Fig. 11). Compound **1** is positioned near the active-site entrance, with its α-carboxyl and α-amine forming hydrogen bonds with Asn421 and Asp420, respectively (Fig. 3c). The N-OH hydroxyl group engages in a water-mediated hydrogen bond with Asn143, while the N^6^-H group interacts with a phosphate oxygen.

Collectively, structural analysis identifies Arg390, Arg425, Asn421, Thr141, Gln182, and Ser139 as key residues for glutamate recognition (Fig. 3a), while the binding mode of **1** suggests that Asn421, Asp420, Glu284, Glu286, Asn184, Asn143, and Leu424 contribute to its interaction with PyrN (Fig. 3c). To assess the functional importance of these residues, site-directed mutagenesis followed by in vitro activity assays was performed (Supplementary Fig. 12). LC-MS analysis revealed that all variants exhibited severely reduced or abolished activity, underscoring the precise tuning of the PyrN active site for substrate binding and catalysis (Fig. 3d and Supplementary Fig. 13).

### Mechanistic study of PyrN-C-catalyzed reaction via computational approaches

Because substrate **1** could not be resolved in the PyrN-C-DGA co-crystal structure, we investigated *O*-aminoacyl-hydroxylamine formation computationally, using the PyrN-C-DGA complex docked with **1**. The docking model was based on the ternary PyrN-C-LGA-**1**. For simulations, two protonation states of hydroxylamine were considered: the zwitterionic form and the neutral form. Substrate **1** in each form was independently docked into the PyrN-C-DGA complex, followed by 100 ns molecular dynamics (MD) simulations. In both cases, the substrate maintained a stable binding conformation throughout the simulations (Supplementary Fig. 14 and Supplementary Fig. 15).

Representative binding conformations from the MD trajectories were then selected for further mechanistic analysis. To determine which form of hydroxylamine is favored in the active site, we performed quantum mechanics (QM) calculations (Supplementary Fig. 16). It was found that the zwitterionic form is energetically favored by 5.9 kcal/mol over the neutral form, indicating that the zwitterionic species is the thermodynamically preferred form within the enzymatic environment. To test the reactivity of **1** in its neutral conformation, we carried out additional QM/MM simulations (Supplementary Fig. 17). For the neutral conformer, oxygen-centered nucleophilic attack exhibited an activation barrier of 22.8 kcal/mol, while nitrogen-centered nucleophilic attack required 33.6 kcal/mol. Both pathways involve prohibitively high activation energies, rendering them unlikely to occur under physiological enzymatic conditions.

Consequently, zwitterionic **1** and d-Glu-AMP were used for substrate modeling. In the subsequent QM/MM simulations, the QM region comprised **1**, d-Glu-AMP, Thr141, and Asn421. Residues Thr141 and Asn421 were included due to their hydrogen-bonding interactions with atoms involved in bond cleavage and charge redistribution (Supplementary Fig. 18). The QM/MM energy profile was calculated for the PyrN-C-catalyzed conversion of **1** and d-Glu-AMP to *O*-aminoacyl-hydroxylamine, starting from the Md-equilibrated representative structure (Fig. 4a and 4b). The results support a concerted mechanism for *O*-aminoacyl-hydroxylamine formation (Fig. 4c). Beginning from the reactant complex (RC1), our QM/MM calculations show that the reaction proceeds via: (i) nucleophilic attack by the hydroxylamine oxygen of **1** on the ester carbonyl carbon of d-Glu-AMP, (ii) simultaneous proton transfer from the hydroxylamine nitrogen to the phosphate oxygen of d-Glu-AMP, and (iii) synchronous cleavage of the C(acyl)-O(P) bond in d-Glu-AMP (Fig. 4c). This concerted step experiences a barrier of 17.0 kcal/mol and yields the ester product complex PC1, which is exothermic by 1.6 kcal/mol relative to RC1 (Fig. 4a). Importantly, the steric inaccessibility of the nitrogen atom in zwitterionic **1** rules out nitrogen-centered nucleophilic attack, confirming that oxygen-centered nucleophilic attack is the exclusive viable pathway.

**Fig. 4.**
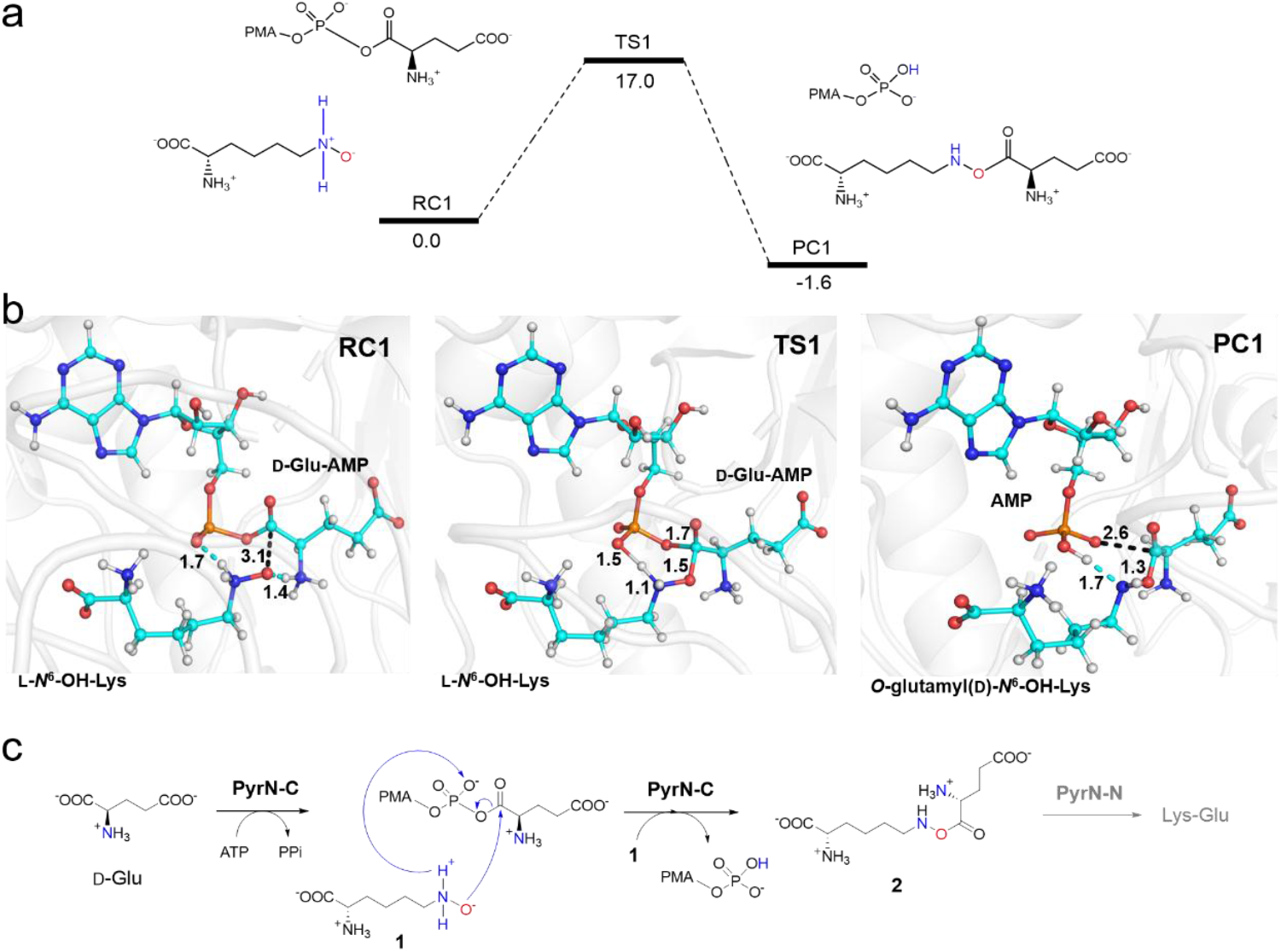
(a) QM(B3LYP-D3/B2)/MM-calculated energy profile (in kcal mol^−1^) for the oxygen nucleophilic attack of substrate d-Glu-AMP and *N*^6^-OH-l-Lys in zwitterionic form. The ZPEs are included in the relative energies. (b) QM(B3LYP-D3/B1)/MM-optimized structures of key species involved in the reaction. Key distances are given in Å. (c) Proposed catalytic mechanism for PyrN-catalyzed reaction.

Although d-Glu is the physiological substrate for PyrN, exhibiting significantly higher activity than l-Glu (Supplementary Fig. 19), the successful co-crystallization of PyrN-C with GSA or l-Glu-AMP confirmed its ability to recognize l-Glu. To investigate the structural and mechanistic basis of this pronounced activity difference, QM/MM calculations were performed on the zwitterionic PyrN-C-LGA-**1** complex (Supplementary Fig. 20). Like the d-configured substrate, l-Glu-AMP reacts via a concerted single-step pathway. Starting from the reactant complex RC2, the reaction overcomes an activation barrier of 19.9 kcal/mol to form the product complex PC2, which is exothermic by 3.2 kcal/mol. The kinetic barrier for the d-Glu-AMP reaction is thus 2.9 kcal/mol lower than that of l-Glu-AMP reaction, in agreement with the experimentally observed activity difference within error margins.

To investigate the origin of the d/l configurational energy barrier difference, we analyzed key interatomic distances from MD simulations: (1) the O-N distance between the hydroxylamine oxygen of **1** and the carbonyl carbon of Glu-AMP, and (2) the C-O distance between the same carbonyl carbon and the nitrogen of the amine group attached to the chiral carbon of Glu-AMP (Supplementary Fig. 21 and Fig. 22). QM/MM-optimized geometries revealed significantly shorter precatalytic distances for d-Glu-AMP (O-N = 2.57 Å; C-O = 3.06 Å) compared with l-Glu-AMP (O-N = 2.63 Å; C-O = 3.20 Å. Statistical analysis of MD trajectories corroborated this trend: mean distances for d-Glu-AMP were O-N = 2.84 Å and C-O = 3.35 Å, versus O-N = 3.04 Å and C-O = 3.44 Å for l-Glu-AMP.

These results support that stereochemical configuration directly modulates critical interatomic distances in the reactant state, thereby exerting a pronounced influence on the reaction energy barrier. The superior activity of d-Glu-AMP can be attributed to stereoelectronic advantages conferred by its configuration. First, the inward orientation of the hydrogen atom at d-Glu-AMP chiral center minimizes steric hindrance during nucleophilic attack by the hydroxylamine oxygen of **1**. Second, the outward projection of the ammonium group (-NH_3_^+^) on this chiral carbon positions it in proximity to the O^−^ group of **1**, enabling formation of a robust salt bridge. This electrostatic attraction both shortens the O-N distance and draws the O^−^ nucleophile toward the substrate, concomitantly reducing the C-O distance along the reaction trajectory. Together, these stereoelectronic effects enhance spatial preorganization, optimally positioning the nucleophile for efficient catalysis.

### Structure-guided targeted mining reveals new hydrazine synthetase homologs

In our previous work, enzyme mining based solely on primary amino acid sequence led to the identification of PyrN homologs capable of utilizing altered amino acid substrates, enabling the discovery of enzymes that accept serine, alanine, or tyrosine for hydrazine production^36^. Structural analysis of PyrN-C-glutamyl-adenylate complexes highlighted residues within 5 Å of the glutamyl moiety as key determinants of amino acid substrate recognition. Consistent with this observation, these residues are highly conserved among glutamate-utilizing hydrazine synthetases (Supplementary Fig. 23). Building on these insights, we attempted to identify homologs recognizing previously unreported substrates, via a structure-guided mining strategy.

To this end, we aligned the PyrN-C-DGA crystal structure with AlphaFold-predicted models of characterized homologs known to recognize glycine, serine, alanine, and tyrosine (Fig. 5a and Supplementary Fig. 24-27).^12^ Examination of eleven residues within the putative amino acid–binding pocket revealed a pattern: four positions (3, 4, 6, and 10), which interact with the α-amino and α-carboxyl groups of the aminoacyl-adenylate, were relatively conserved, whereas seven positions (1, 2, 5, 7, 8, 9, and 11) were variable and likely govern side-chain specificity (Fig. 5b). We therefore designated these variable residues as the “specificity code”. For example, glutamate-utilizing PyrN exhibits the code “SFVALRR” (N- to C-terminus). This specificity code system provides a framework for the targeted discovery and engineering of hydrazine biosynthetic enzymes with novel substrate preferences. Of note, Matsuda *et al*. recently used AlphaFold-predicted structures to analyze putative substrate-binding residues in characterized hydrazine synthetases, proposing an eight-residue specificity code (corresponding to positions 1, 2, 5, 6, 7, 8, 9, and 10 in Fig. 5b), exemplified by “SFVQALNR” in PyrN^37^. A key distinction between the two code systems is that our model mainly focused on residues potentially interacting with the amino acid side chain, and additionally includes position 11 (e.g., Arg425 in PyrN), which our co-crystal structures reveal to be critical for side-chain recognition.

**Fig. 5.**
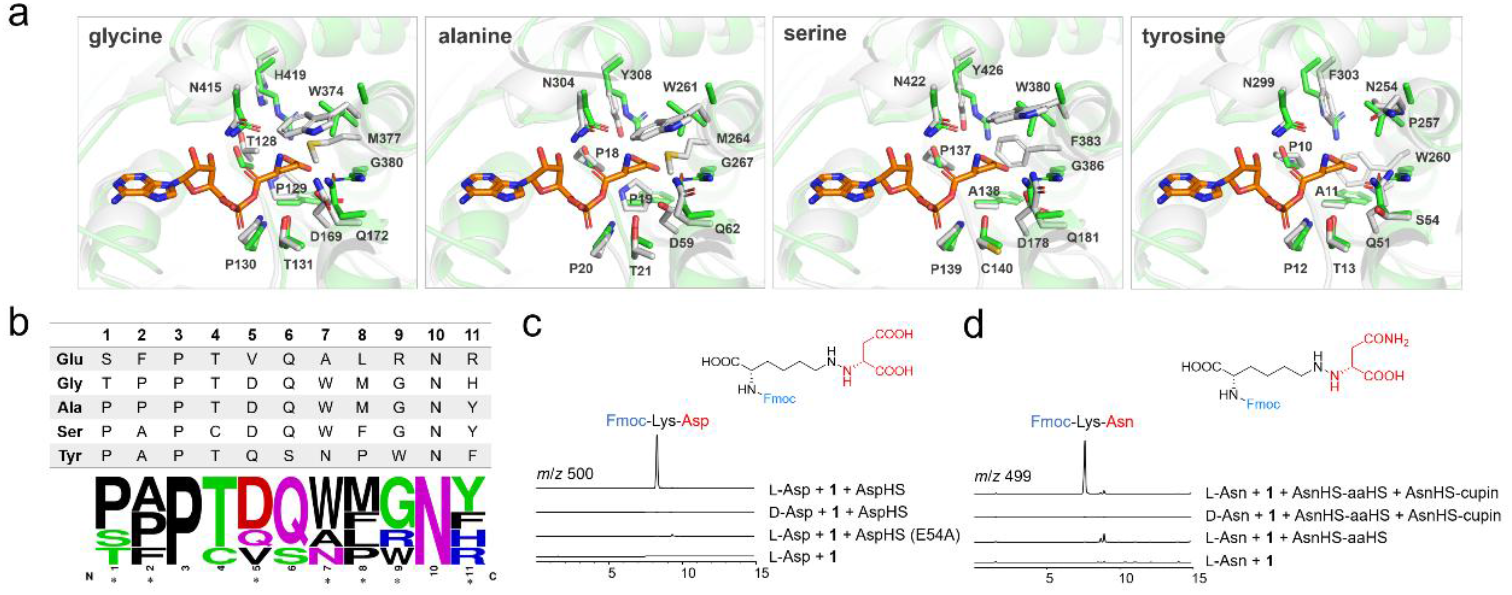
Structure-guided discovery for new hydrazine biosynthetic enzymes with altered amino acid specificity. (a) Structure alignment of PyrN-C-DGA complex with predicted structural models of characterized hydrazine synthetases that utilize Gly, Ala, Ser, and Tyr. Extended versions are provided in Supplementary Fig 24-27. (b) Sequence logo of the amino acid-binding pocket residues from panel (a). Asterisks mark the residues identified as the ‘specificity code’. (c) and (d) LC-MS analysis of the in vitro reaction mixtures of AspHS (c) and AsnHS (d) after Fmoc-Cl derivatization. Note: the EICs for the [M+H]^+^ ions of the corresponding Fmoc-derived hydrazine products (Lys-Asp or Lys-Asn) are displayed. Note: AspHS is didomain hydrazine synthetase, while AsnHS consists of two separated proteins (AsnHS-aaHS and AsnHS-cupin). AspHS (E54A) carries an inactive cupin domain due to the mutation of an essential Glu residue was used as a control.

For targeted enzyme mining, we employed two complementary strategies. First, a UniProt database search using PyrN as the query identified 884 PyrN homologs. AlphaFold structural models for these sequences were directly downloaded, aligned to the PyrN-C-DGA complex structure, and the substrate-specificity code residues were extracted using a custom Python script. Second, we retrieved PyrN-C homologs from the NCBI database that were co-localized with a putative *N*-hydroxylase gene and, in the case of standalone PyrN-C homologs, also with a cupin gene. AlphaFold models for the resulting 830 sequences were generated and analyzed in the same manner to extract their specificity codes. We then selected 20 homologs with specificity codes distinct from previously characterized hydrazine synthetases for the subsequent experimental validation (Supplementary Table 4)^12^. Synthetic genes corresponding to these homologs were co-expressed in *E. coli* together with the lysine *N*^6^-hydoxylase gene *nbtG*^38^, enabling in situ generation of **1**. For standalone PyrN-C homologs, their cognate cupin protein genes were also synthesized and co-expressed. Culture supernatants of the engineered *E. coli* strains were analyzed by LC-MS for lysine-amino acid conjugates. This approach led to the discovery of new PyrN-C or PyrN homologs with previously unreported substrate specificities (Supplementary Table 5). These include the standalone PyrN-C homolog AsnHS from *Burkholderia ubonensis*, which specifically activates l-asparagine, and the didomain PyrN homolog AspHS from *Bradyrhizobium* sp., which recognizes l-aspartic acid (Fig. 5c-5d and Supplementary Fig. 28-31). The activities of these enzymes were further validated by in vitro biochemical assays using purified proteins, followed by LC-HR-MS/MS and NMR analysis of the target products (Supplementary Fig. 32-35).

In addition to enzymes with strict specificity for a single amino acid, we also identified homologs with broadened substrate scope (Supplementary Table 5). These include the didomain enzyme Ser/GlyHS from *Chitinophaga flava*, which accepts both Ser and Gly, and the PyrN-C homolog Ala/GlyHS from *Ralstonia pseudosolanacearum*, which utilizes Ala and Gly (Supplementary Fig. 36-37). Collectively, these newly identified enzymes greatly expand the biocatalyst repertoire available for hydrazine biosynthesis.

### Production of novel hydrazines using unnatural hydroxylamines and hydrazine synthetases

The co-crystal structures of PyrN-C with glutamyl-AMP revealed a solvent-accessible, open substrate-binding cavity (Supplementary Fig. 38), suggesting that PyrN could accommodate alternative hydroxylamines for the biosynthesis of non-natural hydrazines. To test this possibility, we assayed PyrN with a panel of synthetic *N*-substituted hydroxylamines, including *N*-methylhydroxylamine (MHA), *N*-ethylhydroxylamine (EHA), *N*-hexylhydroxylamine (HHA), *N*-aminohexylhydroxylamine (AHHA), and *N*-benzylhydroxylamine (BHA). LC-HR-MS/MS analysis of the in vitro reaction mixtures demonstrated the formation of all the corresponding amine-glutamate conjugates containing N-N bonds, with relative activities quantified via a pyrophosphate release assay (Fig. 6 and Supplementary Fig. 39)^39^.

**Fig. 6.**
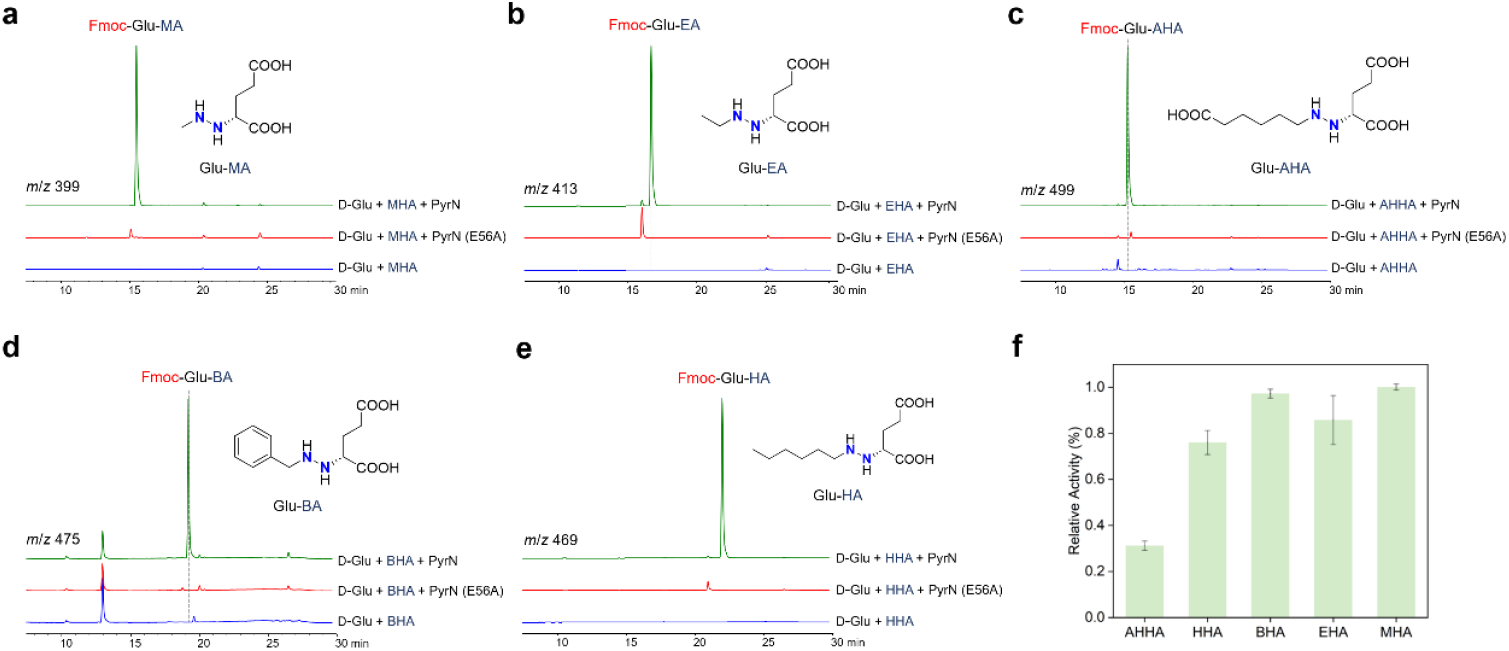
LC-MS analysis of the in vitro reaction mixtures of synthetic *N*-substituted hydroxylamines with hydrazine synthetase PyrN. The EICs for the [M+H]^+^ ions of the corresponding Fmoc-derived hydrazine products are shown. The red traces are the control reaction mixtures using the PyrN(E56A) variant enzyme. This variant contains a non-functional PyrN-N cupin domain due to the mutation of the Glu56 residue that is essential for the N-N formation step^12^. Please also see Supplementary Fig 39 for the detailed LC-HR-MS/MS analysis.

We next examined hydroxylamine specificity in two additional hydrazine synthetases: the serine-utilizing SerHS from *Microcystis aeruginosa* (Uniprot: A0A552E3D4) and glycine-utilizing GlyHS from *Rhizobium loti* (Uniprot: Q8KGM6) (Supplementary Fig. 40)^12^. Both enzymes accepted the synthetic *N*-substituted hydroxylamines, but their cupin domains displayed distinct efficiencies in processing the corresponding *O*-aminoacyl hydroxylamine intermediates (Supplementary Fig. 41-51). The cupin domain of SerHS (SerHS-N) efficiently converted these intermediates into the corresponding hydrazine products, whereas GlyHS-N exhibited lower efficiency, often accumulating amide shunt products due to the instability of *O*-aminoacyl hydroxylamines (Supplementary Fig. 46-51). Notably, GlyHS produced the expected hydrazine products only with BHA and HHA, as confirmed by comparison with synthetic standards (Supplementary Fig. 49 and Fig. 50). Together, these findings demonstrate that relaxed hydroxylamine specificity is a conserved feature of hydrazine synthetases, underscoring their potential as versatile biocatalysts for chemoenzymatic N–N bond formation and the synthesis of non-natural hydrazine derivatives.

## Discussion

Enzymes that catalyze N-N bond formation have recently attracted considerable attention due to their unusual biochemistry and their potential applications in biocatalysis and synthetic biology. A conserved biochemical logic for intermolecular N-N formation involves the generation of an N-O-linked ester intermediate, *O*-aminoacyl hydroxylamine, from an amino acid and an *N*-substituted hydroxylamine, followed by an intramolecular rearrangement that yields the N–N bond. Here, using the glutamate-utilizing hydrazine synthetase PyrN as a model, we report the first crystal structure of the *O*-aminoacyl hydroxylamine synthetase (aaHS) component of a hydrazine synthetase, including binary and ternary co-crystal structures with substrate, catalytic intermediate, and structure mimic. These structures uncover atomic-level determinants of substrate recognition and provide crucial mechanistic insights into this conserved enzymatic strategy for N-N bond formation in nature.

Together with our previous characterization of the cupin domain^12^, our results establish a detailed mechanistic framework for hydrazine synthetase-catalyzed N-N bond formation. The reaction begins with nucleophilic attack of the amino acid α-carboxylate oxygen on the α-phosphate of Mg-ATP, generating an aminoacyl-adenylate intermediate. Hydroxylamine, in its zwitterionic form, then attacks the carbonyl carbon of this intermediate, releasing AMP and producing the key O-aminoacyl hydroxylamine species. This unstable intermediate subsequently undergoes intramolecular rearrangement catalyzed by the zinc-binding cupin domain, leading to N–N bond formation.

The aaHS component of hydrazine synthetases shares clear evolutionary ancestry with class I aminoacyl-tRNA synthetases (aaRSs)^40,41^. Both enzyme families activate amino acids via an enzyme-bound adenylate intermediate and exhibit conserved structural features. However, their catalytic pathways diverge: aaRSs transfer the aminoacyl group to the 3′-OH of tRNA to form aminoacyl-tRNA, whereas aaHSs transfer it to the N-OH group of hydroxylamine, yielding *O*-aminoacyl hydroxylamine. In addition, while aaRSs employ proofreading mechanisms to ensure stringent fidelity in amino acid and tRNA selection during protein synthesis, aaHS domains display striking promiscuity toward structurally diverse synthetic hydroxylamines. This relaxed substrate tolerance, combined with the growing repertoire of amino acid–specific hydrazine synthetases, highlights their exceptional potential as versatile biocatalysts for the chemoenzymatic synthesis of N-N bond-containing compounds.

In summary, our integrated structural, computational, and biochemical studies provide a comprehensive characterization of the bacterial hydrazine synthetase family. By elucidating the substrate recognition and catalytic principles, this work lays a foundation for the discovery and rational engineering of these widespread N-N bond-forming enzymes for advanced biocatalytic applications.

## ACKNOWLEDGMENT

This work was supported by the National Key R&D Program of China (2025YFA0921000), the National Natural Science Foundation of China (32370051 and 32122005) to Y.-L.D., Guangdong Major Project of Basic and Applied Basic Research (2023B0303000026) and the Natural Science Foundation of Guangdong Province (2023A1515012763) to Z.-M.Z., Fundamental Research Funds for the Central Universities (20720240124) to B. W, and the China Postdoctoral Science Foundation (2024M752836, 2024T170784) and Postdoctoral Fellowship Program of CPSF (GZB20230658) to G. Z. We thank Jiaojiao Wang and Zhiwei Ge (Analysis Center of Agrobiology and Environmental Sciences, Zhejiang University) for the assistance of LC-HR-MS/MS data collection. We thank the staff at beamlines BL18U1 or BL10U2 in the Shanghai Synchrotron Radiation Facility (SSRF) and BL19U1 of the National Facility for Protein Science (Shanghai) for access and help with data collection.

## REFERENCES

1. Blair, L. M. & Sperry, J. Natural products containing a nitrogen-nitrogen bond. J Nat Prod 76, 794–812 (2013).

2. He, H.-Y., Niikura, H.Du, Y.-L. & Ryan, K. S. Synthetic and biosynthetic routes to nitrogen-nitrogen bonds. Chem Soc Rev 51, 2991–3046 (2022).

3. Wang, H. et al. Nitrene-mediated intermolecular N-N coupling for efficient synthesis of hydrazides. Nat Chem 13, 378–385 (2021).

4. Barbor, J. P. et al. Development of a Nickel-Catalyzed N–N Coupling for the Synthesis of Hydrazides. J Am Chem Soc 145, 15071–15077 (2023).

5. Chen, L., Deng, Z. & Zhao, C. Nitrogen-Nitrogen Bond Formation Reactions Involved in Natural Product Biosynthesis. ACS Chem Biol 16, 559–570 (2021).

6. Katsuyama, Y. & Matsuda, K. Recent advance in the biosynthesis of nitrogen-nitrogen bond-containing natural products. Curr Opin Chem Biol 59, 62–68 (2020).

7. Du, Y.-L., He, H.-Y., Higgins, M. A. & Ryan, K. S. A heme-dependent enzyme forms the nitrogen-nitrogen bond in piperazate. Nat Chem Biol 13, 836–838 (2017).

8. Matsuda, K. et al. Discovery of Unprecedented Hydrazine-Forming Machinery in Bacteria. J Am Chem Soc 140, 9083–9086 (2018).

9. Sugai, Y., Katsuyama, Y. & Ohnishi, Y. A nitrous acid biosynthetic pathway for diazo group formation in bacteria. Nat Chem Biol 12, 73–75 (2016).

10. Waldman, A. J. & Balskus, E. P. Discovery of a Diazo-Forming Enzyme in Cremeomycin Biosynthesis. J. Org. Chem. 83, 7539–7546 (2018).

11. Kawai, S. et al. Complete Biosynthetic Pathway of Alazopeptin, a Tripeptide Consisting of Two Molecules of 6-Diazo-5-oxo-l-norleucine and One Molecule of Alanine. Angew Chem Int Ed Engl 60, 10319–10325 (2021).

12. Zhao, G. et al. Molecular basis of enzymatic nitrogen-nitrogen formation by a family of zinc-binding cupin enzymes. Nat Commun 12, 7205 (2021).

13. Del Rio Flores, A. et al. Biosynthesis of triacsin featuring an N-hydroxytriazene pharmacophore. Nat Chem Biol 17, 1305–1313 (2021).

14. Shi, J. et al. Conserved Enzymatic Cascade for Bacterial Azoxy Biosynthesis. J. Am. Chem. Soc. 145, 27131–27139 (2023).

15. Zheng, Z. et al. Reconstitution of the Final Steps in the Biosynthesis of Valanimycin Reveals the Origin of Its Characteristic Azoxy Moiety. Angewandte Chemie 136, e202315844 (2024).

16. Twigg, F. F. et al. Identifying the Biosynthetic Gene Cluster for Triacsins with an N-Hydroxytriazene Moiety. Chembiochem 20, 1145–1149 (2019).

17. Zhao, G. et al. The Biosynthetic Gene Cluster of Pyrazomycin-A C-Nucleoside Antibiotic with a Rare Pyrazole Moiety. Chembiochem 21, 644–649 (2020).

18. Ren, D. et al. Identification of the C-Glycoside Synthases during Biosynthesis of the Pyrazole-C-Nucleosides Formycin and Pyrazofurin. Angew Chem Int Ed Engl 58, 16512–16516 (2019).

19. Zhang, M. et al. Comparative Investigation into Formycin A and Pyrazofurin A Biosynthesis Reveals Branch Pathways for the Construction of C-Nucleoside Scaffolds. Appl Environ Microbiol 86, e01971–19 (2020).

20. Van Cura, D. et al. Discovery of the Azaserine Biosynthetic Pathway Uncovers a Biological Route for α-Diazoester Production. Angew Chem Int Ed Engl 62, e202304646 (2023).

21. Shikai, Y., Kawai, S., Katsuyama, Y. & Ohnishi, Y. In vitro characterization of nonribosomal peptide synthetase-dependent O-(2-hydrazineylideneacetyl)serine synthesis indicates a stepwise oxidation strategy to generate the α-diazo ester moiety of azaserine. Chem. Sci. 14, 8766–8776 (2023).

22. Wei, Z.-W., Niikura, H., Wang, M. & Ryan, K. S. Identification of the Azaserine Biosynthetic Gene Cluster Implicates Hydrazine as an Intermediate to the Diazo Moiety. Org Lett 25, 4061– 4065 (2023).

23. Li, W. et al. Discovery of a Bacterial Hydrazine Transferase That Constructs the N-Aminolactam Pharmacophore in Albofungin Biosynthesis. J Am Chem Soc 146, 13399–13405 (2024).

24. Wang, Y. et al. Nitramine Formation via a Cryptic Non-Ribosomal Peptide Synthetase-Dependent Strategy in N-Nitroglycine Biosynthesis. Angew Chem Int Ed Engl 64, e202507866 (2025).

25. Shikai, Y., Muramatsu, H., Igarashi, M., Katsuyama, Y. & Ohnishi, Y. Identification of a l-Threonine-Utilizing Hydrazine Synthetase for Thrazarine Biosynthesis in Streptomyces coerulescens MH802-fF5. Chembiochem e2500298 (2025) doi:10.1002/cbic.202500298.

26. Garg, R. P., Qian, X. L., Alemany, L. B., Moran, S. & Parry, R. J. Investigations of valanimycin biosynthesis: elucidation of the role of seryl-tRNA. Proc Natl Acad Sci U S A 105, 6543–6547 (2008).

27. Zheng, Z., Ren, D., Ko, Y. & Liu, H. Revision of the Formycin A and Pyrazofurin Biosynthetic Pathways Reveals Specificity for d-Glutamic Acid and a Cryptic N-Acylation Step During Pyrazole Core Formation. J Am Chem Soc 147, 11425–11431 (2025).

28. Matsuda, K., Nakahara, Y., Choirunnisa, A. R., Arima, K. & Wakimoto, T. Phylogeny-guided Characterization of Bacterial Hydrazine Biosynthesis Mediated by Cupin/methionyl tRNA Synthetase-like Enzymes. Chembiochem 25, e202300838 (2024).

29. Holm, L. Dali server: structural unification of protein families. Nucleic Acids Res 50, W210– W215 (2022).

30. Crepin, T., Schmitt, E., Blanquet, S. & Mechulam, Y. Three-dimensional structure of methionyl-tRNA synthetase from Pyrococcus abyssi. Biochemistry 43, 2635–2644 (2004).

31. Crepin, T. et al. Use of analogues of methionine and methionyl adenylate to sample conformational changes during catalysis in Escherichia coli methionyl-tRNA synthetase. J Mol Biol 332, 59–72 (2003).

32. Brkic, A. et al. Antibiotic hyper-resistance in a class I aminoacyl-tRNA synthetase with altered active site signature motif. Nat Commun 14, 5498 (2023).

33. Shimada, A., Nureki, O., Goto, M., Takahashi, S. & Yokoyama, S. Structural and mutational studies of the recognition of the arginine tRNA-specific major identity element, A20, by arginyl-tRNA synthetase. Proc Natl Acad Sci U S A 98, 13537–13542 (2001).

34. Mechulam, Y. et al. Crystal structure of Escherichia coli methionyl-tRNA synthetase highlights species-specific features. J Mol Biol 294, 1287–1297 (1999).

35. Abramson, J. et al. Accurate structure prediction of biomolecular interactions with AlphaFold 3. Nature 630, 493–500 (2024).

36. Zhao, G. et al. Molecular basis of enzymatic nitrogen-nitrogen formation by a family of zinc-binding cupin enzymes. Nat Commun 12, 7205 (2021).

37. Matsuda, K., Nakahara, Y., Choirunnisa, A. R., Arima, K. & Wakimoto, T. Phylogeny-guided Characterization of Bacterial Hydrazine Biosynthesis Mediated by Cupin/methionyl tRNA Synthetase-like Enzymes. Chembiochem 25, e202300838 (2024).

38. Binda, C. et al. An unprecedented NADPH domain conformation in lysine monooxygenase NbtG provides insights into uncoupling of oxygen consumption from substrate hydroxylation. J Biol Chem 290, 12676–12688 (2015).

39. Maruyama, C., Niikura, H., Takakuwa, M., Katano, H. & Hamano, Y. Colorimetric Detection of the Adenylation Activity in Nonribosomal Peptide Synthetases. Methods Mol Biol 1401, 77– 84 (2016).

40. Ling, J., Reynolds, N. & Ibba, M. Aminoacyl-tRNA synthesis and translational quality control. Annu Rev Microbiol 63, 61–78 (2009).

41. Ibba, M. & Soll, D. Aminoacyl-tRNA synthesis. Annu Rev Biochem 69, 617–650 (2000).

