## Supporting Information for "Structural and mechanistic insights into bacterial hydrazine biosynthesis"

### MATERIALS AND METHODS

#### General methods

DNA primers were purchased from Tsingke Biological Technology (Supplementary Table 1). Reagents were purchased from Merck, Cambridge Isotope Laboratories, New England BioLabs, and Bio Basic Inc. DNA manipulations in *Escherichia coli* strains were carried out according to standard procedures (Ref.). Ampicillin (100 μg $mL^{-1}$), kanamycin (50 μg $mL^{-1}$), spectinomycin (50 μg $mL^{-1}$) and chloramphenicol (25 μg $mL^{-1}$) were used for the selection of recombinant *E. coli* strains.

#### Protein expression and purification

For the construction of protein expression vectors, DNA fragments encoding the target genes were either amplified by PCR or synthesized by Tsingke Biological Technology (China). These fragments were cloned into the expression vectors pET28a or pCDFDuet-1, and the recombinant plasmids were introduced into *E. coli* BL21 (DE3) for protein expression. Site-directed mutagenesis was performed using the Q5 Site-Directed Mutagenesis Kit (NEB), and mutations were confirmed by DNA sequencing.

For protein expression, *E. coli* cells harboring the corresponding expression vectors were grown overnight at 37°C and 200 rpm in 5 mL of Luria–Bertani (LB) broth supplemented with 50 μg/mL kanamycin (for pET28a-derived vectors) or 50 μg/mL spectinomycin (for pCDFDuet-1-derived vectors). A 2.5 mL aliquot of the overnight culture was used to inoculate 750 mL of LB broth containing the appropriate antibiotic. The culture was incubated at 37°C and 200 rpm until the optical density at 600 nm (OD600) reached 0.4–0.6, at which point protein expression was induced by adding isopropyl β-D-1-thiogalactopyranoside (IPTG) to a final concentration of 0.1 mM.

For protein purification, cells were harvested after 20 hours of incubation at 16°C, then resuspended in lysis buffer (300 mM NaCl, 10 mM imidazole, 1 mM DTT, 50 mM Tris-HCl, pH 8.0). The cells were lysed by sonication on ice, and the lysate was centrifuged at 15,000 rpm for 40 min at 4°C to separate soluble and insoluble fractions. His-tagged proteins were purified using nickel-nitrilotriacetic acid (Ni-NTA) resin. The resin was first washed with washing buffer (300 mM NaCl, 50 mM imidazole, 1 mM DTT, 50 mM Tris-HCl, pH 8.0), and the target protein was eluted using elution buffer (300 mM NaCl, 250 mM imidazole, 1 mM DTT, 50 mM Tris-HCl, pH 8.0).

Purified protein fractions were analyzed by SDS-PAGE, then dialyzed overnight against 1 L of storage buffer (150 mM NaCl, 10% glycerol, 50 mM Tris-HCl, pH 8.0). The dialyzed proteins were concentrated and stored at -80°C for future use. Protein concentration was determined at 280 nm using a Nano Spectrophotometer, with extinction coefficients calculated using the online ExPASy ProtParam tool (https://web.expasy.org/protparam/).

**Protein crystallization and Structure determination**

For the expression of PyrN and PyrN-C (two versions of PyrN-C were constructed: 123-668 of PyrN for co-crystallization with GSA, and 109-668 of PyrN for co-crystallization with LGA and DGA), DNA fragments containing the coding regions were amplified from the genome of *Streptomyces candidus* NRRL 3601 by PCR, using primers listed in Supplementary Table 1. Following Ni-NTA purification, PyrN-C was dialyzed for 4 hours in a buffer containing 20 mM Tris (pH 8.0), 150 mM NaCl, and 1mM DTT. Tobacco Etch Virus (TEV) protease was then added at a 1:50 ratio (TEV:protein) and incubated at 4°C overnight to remove the $His_6$-tag.

For crystallization of PyrN-C (123-668 of PyrN), the TEV-cleaved PyrN-C was dialyzed into a buffer containing 20 mM Tris-HCl (pH 8.0), 50 mM NaCl, 1 mM DTT, then concentrated and loaded onto a HiLoad Superdex 16/600 200pg size-exclusion column pre-equilibrated with a buffer containing 20 mM Tris-HCl (pH 8.0), 50 mM NaCl, and 1mM DTT. The protein was eluted at a flow rate of 1 ml $min^{-1}$, and fractions containing purified protein were pooled and concentrated to a final concentration of ~20 mg $ml^{-1}$.

Preliminary screenings were performed with crystallization screen kits (Crystal Screen HT, Index HT Screen, PEG/ion, SalRX from Hampton Research, and MCSG-1, MCSG-2, MCSG-3, MCSG-2 from Anatrace) by sitting-drop vapor diffusion at 16 °C. The protein (20 mg $ml^{-1}$ and 10 mg $ml^{-1}$) was first incubated with GSA at a ratio of 1:4 for 30 min, and then used to set 2 μl 1:1 ratio drops over 50 μl of reservoir solution. Crystals of the PyrN-C–GSA complex were grown by sitting-drop vapor diffusion using a reservoir solution consisting of 0.1M MES monohydrate (pH 6.5), 13% PEG20000, 4%-(±)1,3-butanediol. Crystals appeared after approximately 1 week, and were cryo-protected with the reservoir solution containing with 15% ethylene glycol and flash-frozen in liquid nitrogen before X-ray data collection.

To obtain the PyrN-C structures in complex with reaction intermediates, the protein (109-668 of PyrN) was incubated with ATP, L/D-Glu at a molar ration of 1 : 2 : 10 on ice for 1 h, and then clarified by centrifugation at 12000 rpm at 4°C for 5 min. Crystals were obtained using hanging drop vapor diffusion method at 20°C by mixing 1 μL of the reaction solution with 1 μL of the reservoir buffer (0.1M Tris (pH9.0), 14% (v/v) PEG8000 and 0.2M $MgCl_2$). Crystals of PyrN-C grown in the presence of L/D-Glu and ATP were soaked with 5 mM of **1** added in the crystallization drops to obtain crystals of PyrN-C-LGA/DGA-**1** complex.

The data of PyrN-C complex were collected at Beam line BL18U1 or BL10U2 in the Shanghai Synchrotron Radiation Facility (SSRF) and BL19U1 of the National Facility for Protein Science in Shanghai. The datasets all indexed, integrated, and scaled using the XDS package[1] or the Aquarium[2]. The co-crystalized structure was determined using molecular replacement using PHASER[3] in Phenix[4]. The models were then refined with iterative cycles of manually building in COOT[5] and refinement in Phenix.

#### In vitro biochemical assays

For the in vitro assays of PyrN or its variants, the reaction mixture (100 µl) contained 3 mM $N^6$-OH-L-Lys (**1**), 10 µM PyrN (or its variants), 5 mM ATP, 2 mM D-Glu, 10 mM $MgCl_2$, and 1 mM DTT in 50 mM Tris-HCl buffer (pH 8.3).

For the in vitro assays of GlyHS or SerHS, the reaction mixture (100 µl) contained 3 mM synthetic *N*-substituted hydroxylamines, 25 µM GlyHS or SerHS, 5 mM ATP, 20 mM L-Gly or L-Ser, 10 mM $MgCl_2$, and 1 mM DTT in 50 mM Tris-HCl buffer (pH 8.3).

The reaction mixtures were incubated for 3 h at 30 °C and then quenched with two volumes of acetonitrile. The supernatants were collected and subjected to LC-MS analysis following pre-column Fmoc-Cl derivatization, which was performed as described previously.[6] LC-MS was performed using an Agilent 1260 II-6125 system, with an Agilent Eclipse XDB-C18 column (5 µm, 4.6 mm ID × 250 mm). Elution was carried out at 1 mL/min with a mobile-phase gradient consisting of water and acetonitrile (v/v): 85:15 for 0–20 min, and 5:95 for 20–25 min, both containing 0.05% (v/v) formic acid. The detection wavelength was set to 263 nm. LC-HR-ESI-MS/MS was conducted using a Waters UPLC (Waters Corp., USA) coupled to an AB TripleTOF 5600plus mass spectrometer (AB SCIEX, USA).

#### In vivo characterization of hydrazine synthetases

The genes or gene pairs identified through Uniprot or NCBI database mining were codon-optimized and synthesized by Tsingke Biological Technology. For fused didomain or tridomain enzyme genes, they were cloned into the pET28a vector using the NdeI/XhoI sites and co-introduced into *E. coli* strain BL21(DE3) along with pCDFDuet-nbtG for protein expression and metabolite production. For standalone cupin and aaHS proteins, the aaHS genes were cloned into pET28a via the NdeI/XhoI sites, while the cupin genes were inserted into the NdeI/XhoI sites of pCDFDuet-nbtG, creating pCDFDuet-nbtG-cupin vectors. These two vectors (pET28a-aaHS and pCDFDuet-nbtG-cupin) were then co-introduced into *E. coli* for protein expression and metabolite production.

For LC-MS analysis, the culture broth supernatants from each strain were mixed with two volumes of acetonitrile and subjected to Fmoc-chloride derivatization. LC-MS was performed using an Agilent 1260 II-6125 system, with an Agilent Poroshell 120 EC-C18 column (4 µm, 4.6 mm ID × 150 mm). Elution was carried out at 1 mL/min with a mobile-phase gradient consisting of water and acetonitrile (v/v): 85:15 for 0–20 min, and 5:95 for 20–25 min, both containing 0.05% (v/v) formic acid. MS signals corresponding to the twenty Fmoc-lysine-amino acid conjugates were searched across all samples to assess the substrate specificities of the synthesized enzymes. LC-HR-ESI-MS/MS was conducted as described above.

**System setup for computational studies.**

The initial structure of the enzyme was prepared based on the determined crystal structures reported in this study. Here, we assigned the protonation states of titratable residues (His, Glu, Asp) based on pKa values from the PROPKA software[7] and a careful visual inspection of local hydrogen-bonded networks. Histidine residues His173, His181, His271, His369, His540 and His658 in PyrN-C were protonated at the δ position, while all the other histidine residues were protonated at the ε position. Glutamic residues Glu284 in PyrN-C was protonated, while all the other glutamic residues were deprotonated. All of the aspartic acid residues were deprotonated. The Zn(II)-containing site were parametrized using MCPB[8]. Thereafter, the substrate $N^6$-OH-L-Lys was docked into the generated pocket of PyrN-C-DGA structure using the AutoDock Vina[9] tool in Chimera[10] and the lowest binding energy conformer was selected to conduct further molecular dynamics (MD) simulation.

**Classical MD simulations**

The Amber ff14SB[11] force field was selected for treating the amino acid residues of the PyrN-C protein, while the general AMBER force field (GAFF)[12] was employed for the substrate. Besides, the RESP calculations[13] at the B3LYP[14–17]/def2-SVP[18] level of theory was used to define the partial atomic charges of the substrate. The parmchk utility in Amber 20 was used to load the missing parameters of the substrate. Then, we added the sodium ions to the surface of the protein to balance the total charge of the complex systems. Finally, the whole system was solvated in a rectangular box with TIP3P[19] waters, with a minimum distance of 15 Å from the protein surface. After the setup of the system, it was totally minimized by the combined steepest descent and conjugate gradient methods. The system was gently annealed from 0 to 300 K under a canonical ensemble for 50 ps with a small restraint of 25 kcal/mol/Å on the protein. To achieve a uniform density after the heating dynamics, 1 ns of density equilibration was performed under the NPT ensemble, where the target temperature of 300 K was maintained by using the Langevin thermostat[20] with a collision frequency of 2 $ps^{-1}$, while the target pressure of 1.0 atm was controlled by using the Berendsen barostat[21] with a pressure relaxation time of 1 ps. Afterward, we removed all the restraints on the protein and further equilibrated the system for 4 ns under the NPT ensemble to get the stable temperature and pressure. At last, we performed a productive MD simulation under the NPT ensemble for 100 ns. The periodic boundary conditions were used for all MD simulations, while the time-step is 2 fs. The covalent bonds containing hydrogen atoms were constrained using SHAKE[22] to enable an integration step of 2 fs. Nonbonded interactions were treated with Particle Mesh Ewald (PME)[23] with the cutoff set at 8 Å. Representative snapshots from the MD trajectory were used as starting structures for the QM/MM calculations.

#### QM calculation

All QM cluster-continuum model calculations were performed using the Gaussian 16 software. The QM model is comprised of the truncated substrate D-Glu-AMP and $N^6$-OH-L-Lys (**1**). The valence charge of models in zwitterionic form of reactant complex and neutral form of reactant complex are all 0 (Supplementary Fig. 16). The geometries of interested species were fully optimized in water solution in conjunction with the SMD[24] continuum solvation model at the B3LYP[14–17]/def2-TZVP[18] level of theory.

#### QM/MM calculation

For the subsequent QM/MM calculations, we selected the representative snapshot based on the analysis of the distance between target O and target C and the corresponding hydrogen-bond network during the classical MD trajectory. The representative snapshots from MD simulations were selected using the K-means clustering algorithm. Representative snapshots from the most populated cluster were used for the following QM/MM geometric optimization. As shown in Supplementary Fig. 14, the time evolution of the root mean square deviation (RMSD) shows that the MD simulations are well converged in 100 ns. All the QM/MM calculations were performed by the ChemShell[25], in which the turbomole[26] is invoked for the QM region while the DL_POLY[27] is used for the MM region. Besides, the electronic embedding scheme[28] was employed to account for the polarizing effect of the enzyme environment on the QM region, while the hydrogen link atoms with the charge-shift model was used to deal with the QM/MM boundary. The QM/MM system contains the whole protein and solvation waters within 8 Å of protein. Here, the QM region was studied with the hybrid B3LYP[14–17] density functional. For geometry optimization, the double - ζ basis set def2-SVP[18] (labeled as B1) was used. Transition states were located with relaxed potential energy surface (PES) scans followed by full TS optimizations using the dimer optimizer implemented in the DL-FIND code[29]. The energies of all species were further corrected with a larger basis set def2-TZVP[18] (labeled as B2), together with zero-point energies (ZPE) at the B3LYP/B1 level. For this PyrN-C protein, the QM region consists of residues T141, N421 and substrates D-Glu-AMP and $N^6$-OH-L-Lys. The dispersion corrections computed with Grimme's D3 method with BJ-damping[30,31] were included in QM regions in all QM/MM calculations. Atoms included in the QM regions are further demonstrated in Supplementary Fig. 18.

#### Enzyme mining

Using the EFI-Enzyme Similarity Tool[32], we retrieved 1,000 homologous protein sequences for each target enzyme (PyrN, Afn8, Tri28, TyrHS, and SerHS) from the UniProt database. Using the EFI-Genome Neighborhood Tool, we retrieved information on the protein-coding genes located upstream and downstream of each homologous protein, with the neighborhood size set to 20. Pfam protein family annotations were obtained for all neighboring genes. Homologous proteins were classified as putative hydrazine synthetases if their genomic neighborhoods contained genes encoding proteins known to be essential for N-N bond formation, specifically those belonging to the Cupin_2 (PF07883), Lys_Orn_oxygenase (PF13434), and tRNA-synth_1g (PF09334) families. After removing redundant sequences, a total of 990 homologous proteins were identified. The corresponding PDB structures for these homologs were subsequently batch-downloaded based on their UniProt identifiers, yielding 884 protein structures in total. Each homologous protein structure was then aligned to the PyrN-C reference structure to facilitate comparative analysis of the key amino acid residues involved in substrate binding and active site interactions.

For NCBI database mining, Candidate hydrazine synthetases were identified by phmmer (HMMER 3.3.2)[33] searches of the NCBI nr database (downloaded from https://ftp.ncbi.nlm.nih.gov/blast/db/ in Jun 3. 2024). The database includes all segments from nr.00.tar.gz to nr.125.tar.gz (inclusive), totaling 126 compressed archives. The amino acid sequence of PyrN was used as a query, with an E-value threshold of $10^{e-10}$, yielding 108,121 unique sequences. Redundancy reduction was performed using CD-HIT (v4.8.1)[34] with a sequence identity cutoff of 70%, resulting in 4,801 candidate sequences.

For functional context assessment, the genomic neighborhoods of all 4,801 candidate proteins were analyzed. A sequence was tentatively annotated as a hydrazine synthetase if the proteins containing Cupin_2 (PF07883), Lys_Orn_oxygenase (PF13434), and tRNA-synth_1g (PF09334) families—all essential for N-N bond formation—were present within ±20 kb of the candidate locus. This screening resulted in 279 sequences meeting the criterion. These 279 sequences were aligned using ClustalW (https://www.genome.jp/tools-bin/clustalw), and a HMM profile was constructed using hmmbuild (HMMER v3.3). The resulting HMM profile was then used to search the nr database with hmmsearch (HMMER v3.3), applying an *E*-value threshold of $1.0 \times 10^{-125}$. After further redundancy reduction with CD-HIT (identity threshold 0.96), 830 hydrazine synthetase sequences were retained for downstream analysis. Subsequently, structures of the 830 homologs were predicted using AlphaFold, and each predicted structure was aligned to the PyrN-MetRS structure to enable comparative analysis of key amino acid residues involved in substrate binding and active site interactions.

**Supplementary Table 1. Oligonucleotides used in this study**

| Primer name | Sequence (5′→3′) | Discriptation |
|---|---|---|
| PyrN-NdeI-F | agcagcCATATGatcgtcagtgagatcccc | Primers for PyrN expression |
| PyrN-XhoI-R | agcagcCTCGAGtgtcggtgtcggtgtcggtgt | |
| PyrN-C-SNA-TEV-NdeI-F1 | agcagcCATATGGAAAACCTGTACTTCCAATCCAATGCAgaacaggctgtggaacggc | Primers for PyrN-C expression |
| PyrN-C-XhoI-R | agcagcCTCGAGctgcctcccaggtgctgttcg | |
| PyrN-S138A-F | CTGCTGCCGgcgTTCCCCACGCCGAAC | Primers for the point mutation of PyrN |
| PyrN-S138A-R | GGGGAAcgcCGGCAGCAGCAGCACGGG | |
| PyrN-F139A-F | CTGCCGTCGgcgCCCACGCCGAACGGTGAAC | Primers for the point mutation of PyrN |
| PyrN-F139A-R | CGGCGTGGGcgcCGACGGCAGCAGCAGCACG | |
| PyrN-R390A-F | CGGCGTTCGAGCTGGCCGCAgcgTTCCTCACCGCGCTCGACGGATTCGC | Primers for the point mutation of PyrN |
| PyrN-R390A-R | CGTCGAGCGCGGTGAGGAAcgcTGCGGCCAGCTCGAACGCCGAGTACA | |
| PyrN-R425A-F | GCTTCGACAACGCGTTCCTGgcgGCGTTCGCGTTTCCGGCGGTGC | Primers for the point mutation of PyrN |
| PyrN-R425A-R | ACCGCCGGAAACGCGAACGCcgcCAGGAACGCGTTGTCGAAGCCGAAGAACAGCACG | |
| PyrN-T141A-F | TGCTGCTGCCGTCGTTCCCCgcgCCGAACGGTGAACTGCACCTCGGG | Primers for the point mutation of PyrN |
| PyrN-T141A-R | GTGCAGTTCACCGTTCGGcgcGGGGAACGACGGCAGCAGCAGCAC | |
| PyrN-Q182A-F | TGCTCGGCACGGTCGGCCATgcgAGCCAGGTGTCCGCGGCCGCGG | Primers for the point mutation of PyrN |
| PyrN-Q182A-R | CGGCCGCGGACACCTGGCTcgcATGGCCGACCGTGCCGAGCAGCAGGTG | |
| PyrN-N421A-F | GTGCTGTTCTTCGGCTTCGACgcgGCGTTCCTGCGAGCGTTCGCGT | Primers for the point mutation of PyrN |
| PyrN-N421A-R | GCGAACGCTCGCAGGAACGCcgcGTCGAAGCCGAAGAACAGCACGGTGCG | |
| PyrN-L424A-F | GACAACGCGTTCgcgCGAGCGTTCGCGTTTCCG | Primers for the point mutation of PyrN |
| PyrN-L424A-R | GAACGCTCGcgcGAACGCGTTGTCGAAGCCGAAG | |
| PyrN-N143A-F | CCCACGCCGgcgGGTGAACTGCACCTC | Primers for the point mutation of PyrN |
| PyrN-N143A-R | TTCACCcgcCGGCGTGGGGAACGACGGC | |

| | | |
|---|---|---|
| PyrN-Q184A-F | CATCAGAGCgcgGTGTCCGCGGCCGCGGAGGCG | Primers for the point mutation of PyrN |
| PyrN-Q184A-R | CGCGGACACcgcGCTCTGATGGCCGACCGTG | |
| PyrN-E286A-F | GGCATCGAGTGCgcgTTGTGCGCGTTGCCC | Primers for the point mutation of PyrN |
| PyrN-E286A-R | GCACAAcgcGCACTCGATGCCGGCCGTCTG | |
| PyrN-E284A-F | GCCGGCATCgcgTGCCAGTTGTGCGCGTTG | Primers for the point mutation of PyrN |
| PyrN-E284A-R | CAACTGGCAcgcGATGCCGGCCGTCTGGTTCG | |
| PyrN-D420A-F | GGCTTCgcgAACGCGTTCCTGCGAG | Primers for the point mutation of PyrN |
| PyrN-D420A-R | CGCGTTcgcGAAGCCGAAGAACAGC | |
| PyrN-E56A-F | CACGACCTGgcgGTCTGGGTGATGCTCGAC | Primers for the point mutation of PyrN |
| PyrN-E56A-R | CACCCAGACcgcCAGGTCGTGGTGGTTGTG | |
| GlyHS-NdeI-F | agcagcCATATGCGTAAATCTACCTTCTCGCGC | Primers for GlyHS expression |
| GlyHS-XhoI-R | agcagcCTCGAGtcaAGCTTCCTCTTCGTGCAACAC | |
| SerHS-NdeI-F | agcagcCATATGCTAATAAGGAAATTTAATATTGCTG | Primers for SerHS expression |
| SerHS-XhoI-R | agcagcCTCGAGtcaCAAGCTCGGGGTTG | |
| AlaHS-NdeI-F | agcagcCATATGatcacgcgtgccttcgacc | Primers for AlaHS expression |
| AlaHS-HindIII-R | agcagcAAGCTTctgttcgaggacgagcaccgcg | |
| GlyHS-E69A-F | GATGAGATCgcgGCTTTCGTGGTTCTGAGC | Primers for the point mutation of GlyHS |
| GlyHS-E69A-R | CACGAAAGCcgcGATCTCATCATGACGATG | |
| SerHS-E55A-F | CATGAAGGCgcgACTTTCTTCATCATTCAAGGTAAGG | Primers for the point mutation of SerHS |
| SerHS-E55A-R | GAAAGTcgcGCCTTCATGGTGATTGTGGCGC | |
| AspHS-E54A-F | CACGATTCCgcgATCTGGATCGTTGTTGCTG | Primers for the point mutation of SerHS |
| AspHS-E54A-F | CCAGATcgcGGAATCGTGGTGGTTGTGCG | |

**Supplementary Table 2. Data collection and refinement statistics of PyrN-C-GSA**

| Data set | **PyrN-C** (PDB: 9L8V) |
|---|---|
| **Data collection** | |
| Space Group | P121 |
| *a, b, c* (Å) | 50.01 243.81 100.62 |
| α, β, γ (°) | 90 99.04 90 |
| Wavelength (Å) | 0.9793 |
| Resolution (Å) | 34.80-2.23 |
| $R_{merge}$ (%) | 15.6 (122.5) |
| $CC_{1/2}$ | 99.6 (57.0) |
| *Average I/σ(I)* | 9.5 (1.5) |
| Completeness (%) | 99.9(99.9) |
| **Refinement** | |
| Number of measured | 790130 |
| Number of unique reflections | 115199 |
| Redundancy | 6.9 (6.9) |
| $R_{work}$ / $R_{free}$ (%) | 18.73/23.18 |
| No. atoms | 17564 |
| Average B value (Å2) | 40.11 |
| RMSD from ideal values | |
| Bonds (Å) | 0.007 |
| Angle (°) | 0.913 |
| Ramachandran plot statistics | |
| Most favorable | 97.69 |
| allowed | 2.31 |
| Disallowed | 0 |

Values in parentheses are for the highest resolution shell.

**Supplementary Table 3. Data collection and refinement statistics for PyrN-AMP-D-Glu, PyrN-AMP-L-Glu-1, and PyrN-AMP-L-Glu**

| | PyrN-AMP-D-GLU (PDB:9VQJ) | PyrN-AMP-L-GLU $N^6$-OH Lys (PDB:9VR2) | PyrN-AMP-L-GLU (PDB:9VQ4) |
|---|---|---|---|
| **Data collection** | | | |
| Space group | P $2_1$ | P $2_1$ | P $2_1$ |
| Cell dimensions | | | |
| $a$, $b$, $c$ (Å) | 101.24, 50.43, 106.32 | 101.58, 50.53, 106.61 | 106.518, 50.29, 114.127 |
| $\alpha$, $\beta$, $\gamma$ (°) | 90.00, 90.23, 90.00 | 90.00, 89.82, 90.00 | 90.00, 92.52, 90.00 |
| Resolution (Å) | 47.15-2.00(2.04-2.00)[a] | 50.00-2.45 (2.51-2.45) | 50.00-2.10(2.15-2.10) |
| $R_{merge}$ | 0.106(0.600) | 0.184(0.317) | 0.104(0.527) |
| $I/\sigma(I)$ | 11.20(2.70) | 11.87(5.89) | 18.22(3.90) |
| $CC_{1/2}$ | 0.996(0.833) | 0.972(0.938) | 0.990(0.836) |
| Completeness (%) | 100.0(100.0) | 99.6(98.0) | 99.8(99.0) |
| Redundancy | 6.40(4.80) | 6.63(6.68) | 6.67(6.73) |
| **Refinement** | | | |
| No. reflections | 73173 | 40316 | 87158 |
| $R_{work}$ / $R_{free}$ | 0.173/0.210 | 0.169/0.217 | 0.168/0.210 |
| No. atoms | | | |
| Protein | 8125 | 7551 | 8184 |
| Ligand | 68 | 91 | 68 |
| Water | 870 | 647 | 739 |
| $B$ factors | | | |
| protein | 19.2 | 20.90 | 34.70 |
| Ligand | 13.6 | 20.40 | 34.00 |
| Water | 25.4 | 22.30 | 38.90 |
| r.m.s deviations | | | |
| Bond lengths (Å) | 0.003 | 0.010 | 0.003 |
| Bond angles (°) | 0.73 | 1.29 | 0.75 |

[a]Values in parentheses are for highest-resolution shell.

**Supplementary Table 4.** PyrN homologues selected for gene synthesis and the subsequent activity assay through in vivo approach.

| Proteins | Size (aa) | Organism | Description |
|---|---|---|---|
| WP_211346831 | 141 | *Actinokineospora cianjurensis* DSM 45657 | Cupin |
| WP_121394552 | 501 | | MetRS |
| WP_307240806 | 123 | *Kineosporia succinea* DSM 44388 | Cupin |
| WP_307240804 | 513 | | MetRS |
| WP_184911088 | 122 | *Kitasatospora gansuensis* DSM 44786 | Cupin |
| WP_184911086 | 512 | | MetRS |
| WP_093785142 | 127 | *Actinacidiphila guanduensis* CGMCC 4.2022 | Cupin |
| WP_093785150 | 515 | | MetRS |
| WP_312890654 | 135 | *Kutzneria kofuensis* DSM 43851 | Cupin |
| WP_184869844 | 513 | | MetRS |
| WP_314244933 | 136 | *Streptomyces* sp. DSM 40907 | Cupin |
| WP_314244924 | 511 | | MetRS |
| MCP4619998 | 679 | *Bradyrhizobium* sp | Cupin-MetRS |
| WP_189916778 | 115 | *Kitasatospora xanthocidica* JCM 4862 | Cupin |
| WP_189916774 | 543 | | MetRS |
| WP_203996267 | 107 | *Virgisporangium aurantiacum* NBRC 16421 | Cupin |
| WP_203996269 | 481 | | MetRS |
| WP_098134181 | 671 | *Bacillus toyonensis* JAS03 | Cupin-MetRS |
| WP_200520331 | 126 | *Bradyrhizobium diazoefficiens* | Cupin |
| WP_200520330 | 524 | | MetRS |
| WP_313740208 | 134 | *Pseudomonas* sp | Cupin |
| WP_313740207 | 521 | | MetRS |
| WP_245241356 | 137 | *Micromonospora profundi* TRM95458 | Cupin |
| WP_306272881 | 459 | | MetRS |
| WP_067598371 | 651 | *Nocardiopsis listeri* NBRC 13360 | Cupin-MetRS |
| WP_042441710 | 657 | *Streptacidiphilus jiangxiensis* CGMCC 4.1857 | Cupin-MetRS |
| HEX4062545 | 127 | *Streptosporangiaceae bacterium* | Cupin |

| | | | |
|---|---|---|---|
| HEX4062544 | 519 | | MetRS |
| A0A505DEG1 | 780 | *Streptomyces sporangiiformans* | Cupin-MetRS-PCP |
| A0A365Y092 | 655 | *Chitinophaga flava* | MetRS- Cupin |
| A0A0U3GF74 | 682 | *Pseudoalteromonas rubra* | Cupin-MetRS |
| Q8XYD9 | 114 | *Ralstonia nicotianae* ATCC BAA-1114 | Cupin |
| Q8XYE0 | 496 | | MetRS |
| A0A1H1KCL4 | 109 | *Paraburkholderia tuberum* | Cupin |
| A0A1H1KCS5 | 527 | | MetRS |
| A0A5J6J753 | 147 | *Streptomyces vinaceus* CGMCC 4.1305 | Cupin |
| A0A5J6JFZ2 | 547 | | MetRS |
| WP_246732448 | 101 | *Bradyrhizobium yuanmingense* CGMCC 1.3531 | Cupin |
| WP_157286713 | 551 | | MetRS |
| WP_048988468 | 116 | *Burkholderia cenocepacia* | Cupin |
| WP_060213334 | 531 | | MetRS |
| A0A3Q9K6F8 | 356 | *Streptomyces lydicus* CGMCC 4.1412 | Cupin |
| A0A3S9Y4V2 | 524 | | MetRS |
| A0A3A8Q8X4 | 116 | *Corallococcus interemptor* DSM 10883 | Cupin |
| A0A3A8Q8Y3 | 545 | | MetRS |
| WP_095402836 | 123 | *Burkholderia ubonensis* DSM17311 | Cupin |
| WP_095402837 | 506 | | MetRS |
| A0A7X5ZUU5 | 270 | *Sphingomonas leidyi* DSM 4733 | Cupin |
| A0A7X5UYN5 | 525 | | MetRS |

**Supplementary Table 5.** The substrate binding pocket residues of selected PyrN homologs and the amino acid substrate(s).

| NO. | Enzyme | Substrate[a] | Substrate binding pocket residues | | | | | | | | | | |
|---|---|---|---|---|---|---|---|---|---|---|---|---|---|
| | | | 1 | 2 | 3 | 4 | 5 | 6 | 7 | 8 | 9 | 10 | 11 |
| 1 | PyrN | Glu | S138 | F139 | P140 | T141 | V179 | Q182 | A384 | L387 | R390 | N421 | R425 |
| 2 | WP_095402837 | Asn | P9 | P10 | P11 | T12 | D50 | Q53 | G256 | A259 | Y262 | N294 | W298 |
| 3 | MCP4619998 | Asp | A144 | F145 | P146 | T147 | L185 | M188 | Y387 | H390 | R393 | N431 | R435 |
| 4 | A0A505DEG1 | Thr | S133 | A134 | P135 | T136 | D174 | Q177 | W376 | Y379 | S382 | G417 | H421 |
| 5 | WP_307240804 | Tyr | P10 | A11 | P12 | T13 | Q51 | S54 | P257 | W260 | Y263 | N298 | F302 |
| 6 | A0A7W7S6L6 | Tyr | P10 | A11 | P12 | T13 | Q51 | S54 | T257 | W260 | Y263 | N299 | F303 |
| 7 | A0A7W9NL65 | N.D[b] | P11 | A12 | P13 | T14 | G50 | Q55 | H253 | G256 | W259 | A298 | F302 |
| 8 | WP_030394523 | Tyr/Phe | A9 | L10 | P11 | T12 | D50 | E53 | T269 | L272 | T275 | G314 | M320 |
| 9 | WP_200520330 | Ala/Ser | P11 | P12 | P13 | T14 | V53 | Q55 | A253 | L257 | H260 | N286 | Y290 |
| 10 | WP_313740208 | Ala/Ser | P12 | P13 | P14 | T15 | D53 | Q56 | P254 | I258 | Q261 | N289 | Y293 |
| 11 | WP_042441710 | Ala/Gly | P134 | P135 | P136 | T137 | D175 | Q178 | A372 | L376 | T379 | N404 | Y408 |
| 12 | HEX4062544 | Ala/Gly | T54 | A55 | P56 | T57 | D95 | Q98 | W294 | M297 | G300 | N342 | F346 |
| 13 | Q8XYE0 | Ala/Gly | V12 | M13 | P14 | T15 | D53 | E56 | A259 | M262 | V265 | N298 | F302 |
| 14 | A0A1H1KCS5 | Ala/Gly | A11 | P12 | P13 | C14 | D52 | G55 | W254 | G257 | C260 | N295 | Y299 |
| 15 | A0A5J6JFZ2 | Ala/Gly | A9 | P10 | P11 | N12 | D50 | S53 | G259 | G262 | A265 | C299 | H303 |
| 16 | A0A365Y092 | Ser/Gly | A18 | M19 | P20 | T21 | D59 | E62 | G268 | S271 | R274 | S303 | W307 |
| 17 | WP_306272881 | Ala | P21 | P22 | P23 | T24 | D62 | Q65 | N267 | M269 | R273 | N303 | Y307 |
| 18 | WP_067598371 | Ala | E9 | S10 | M11 | T137 | D175 | Q178 | W376 | L379 | T382 | N407 | Y411 |
| 19 | A0A0U3GF74 | Gly | T134 | P135 | P136 | T137 | D175 | Q178 | W377 | M380 | A383 | N421 | F425 |
| 20 | WP_314244924 | N.D[b] | P9 | G10 | P11 | T12 | H50 | Q53 | D253 | T256 | W259 | A298 | F302 |
| 21 | WP_189916774 | Gly | S15 | N16 | P17 | T18 | A56 | N59 | C262 | L265 | R268 | N296 | R300 |

[a]It is worth to mention that, although product formation has been detected, the possibility that their native substrates are other non-proteinogenic amino acids or alternative hydroxylamines cannot be excluded.

[b]N.D, not detected. It is possible that these enzymes are not efficiently expressed in *E. coli* host, or that they require non-proteinogenic amino acids not available in *E. coli*.

#### Supplementary Fig. 1

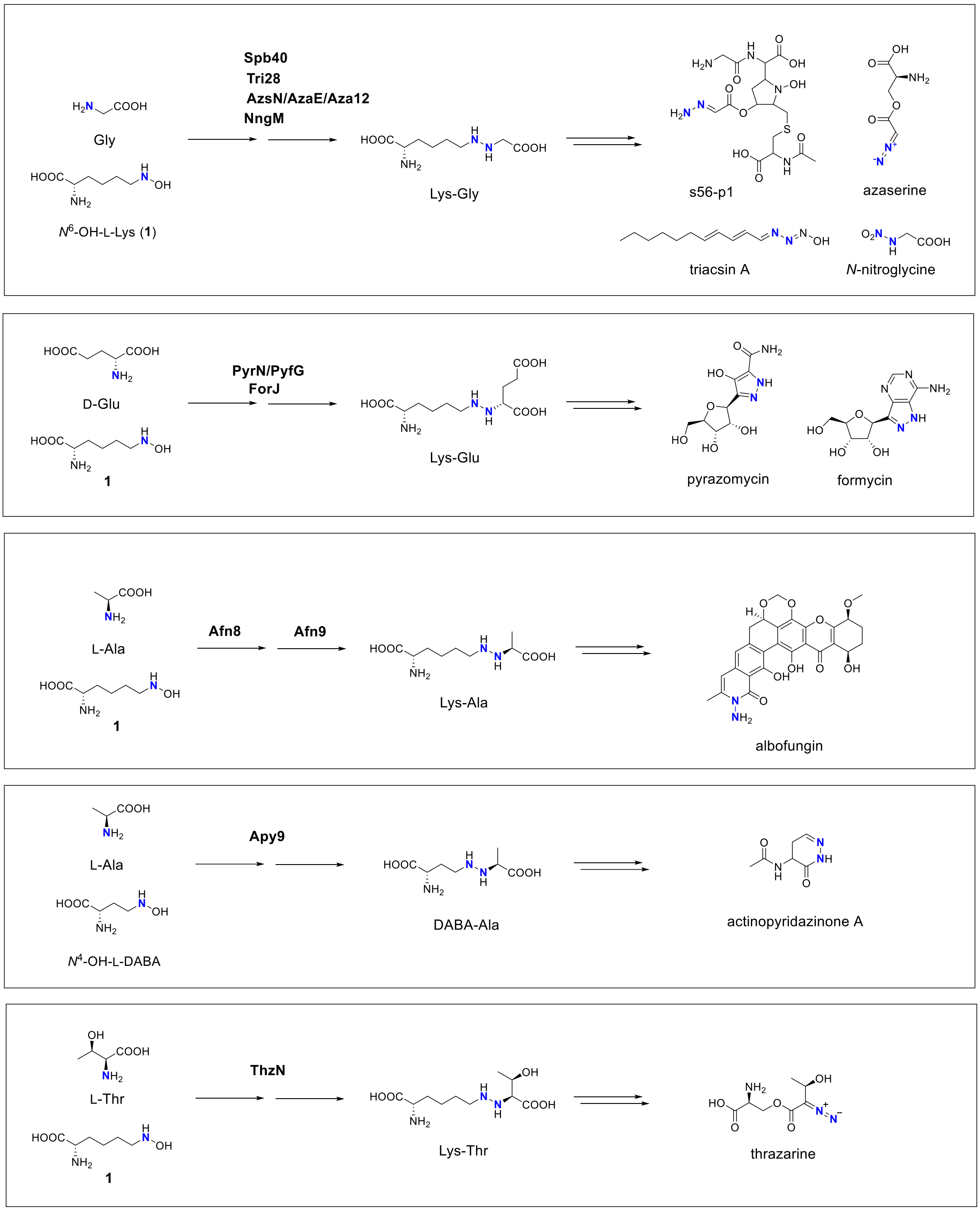

**Supplementary Fig 1.** Reaction schemes for selected hydrazine synthetase (HS)-catalyzed N-N bond formation in natural product biosynthesis.

**Supplementary Fig. 2**

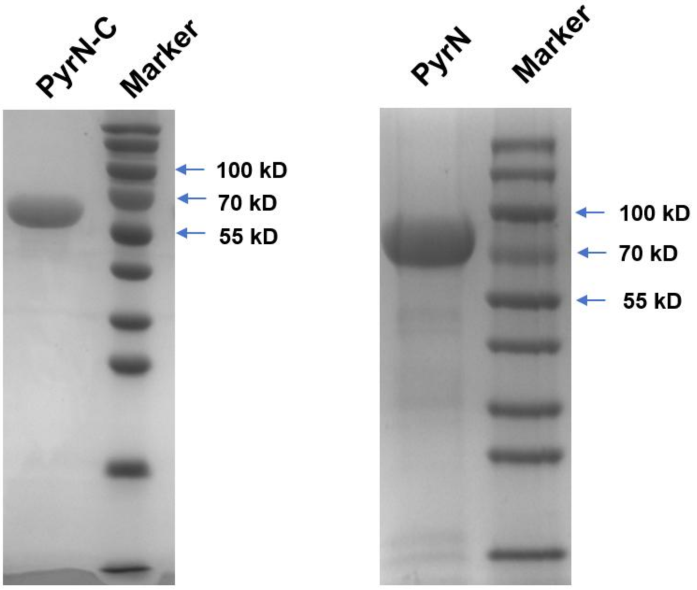

**Supplementary Fig 2.** SDS-PAGE analysis of PyrN-C and PyrN (full-length). Calculated molecular weight for monomeric PyrN-C and $His_6$-PyrN (full-length) are 61.4 kDa and 74.9 kDa, respectively.

#### Supplementary Fig. 3

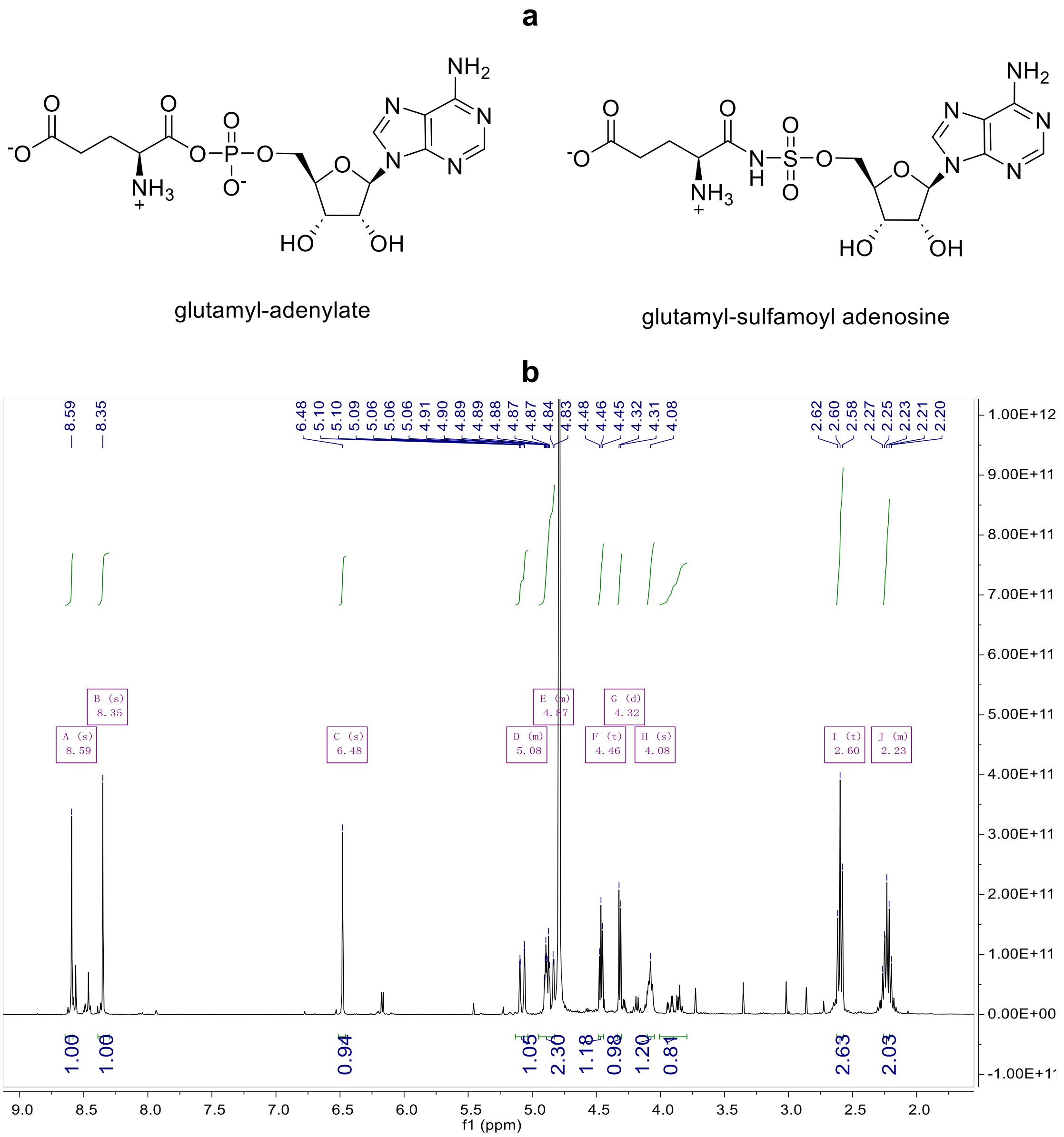

**Supplementary Fig 3.** The structure of L-glutamyl-sulfamoyl adenosine (GSA) and glutamyl-adenylate (a) and the $^{1}$H NMR spectrum of GSA (b). GSA contains an amide ester bond that mimics the phosphate ester linkage in glutamyl-adenylate. Chemical synthesis of GSA was performed similarly as described previously. HRMS of GSA: $[M+H]^+$ calculated for 476.1200, and found as 476.1197. $^{1}$H NMR (400 MHz, $D_2O$): δ 8.59 (s, 1H), 8.35 (s, 1H), 6.48 (s, 1H), 5.16 – 5.03 (d, 1H), 4.96 – 4.82 (m, 2H), 4.46 (t, $J$ = 4.9 Hz, 1H), 4.32 (d, $J$ = 5.4 Hz, 1H), 4.08 (s, 1H), 2.60 (m, 2H), 2.30 – 2.17 (m, 2H).

**Supplementary Fig. 4**

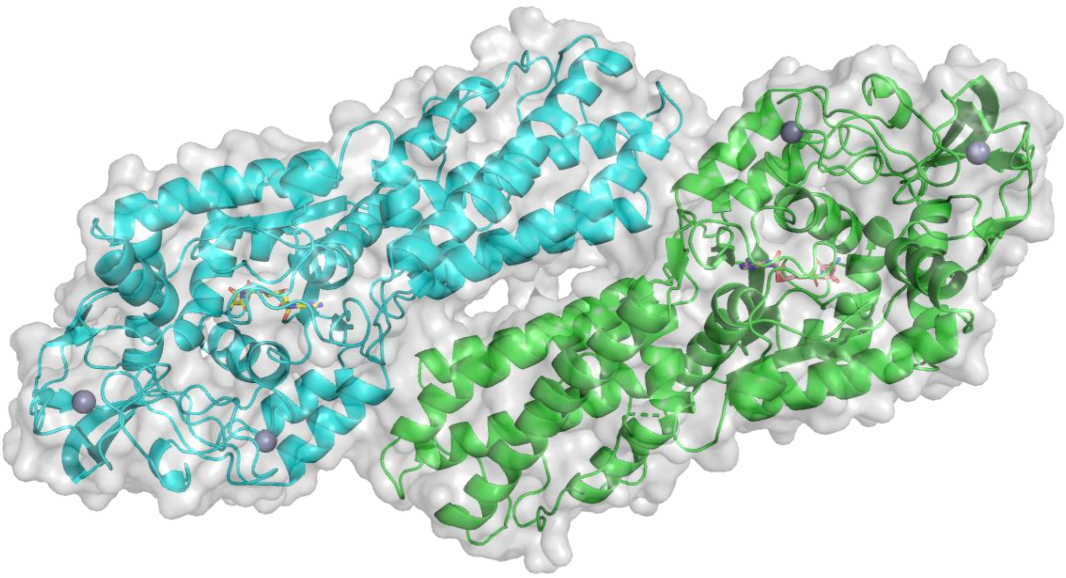

**Supplementary Fig 4.** The homodimeric structure of PyrN-C in complex with GSA.

**Supplementary Fig. 5**

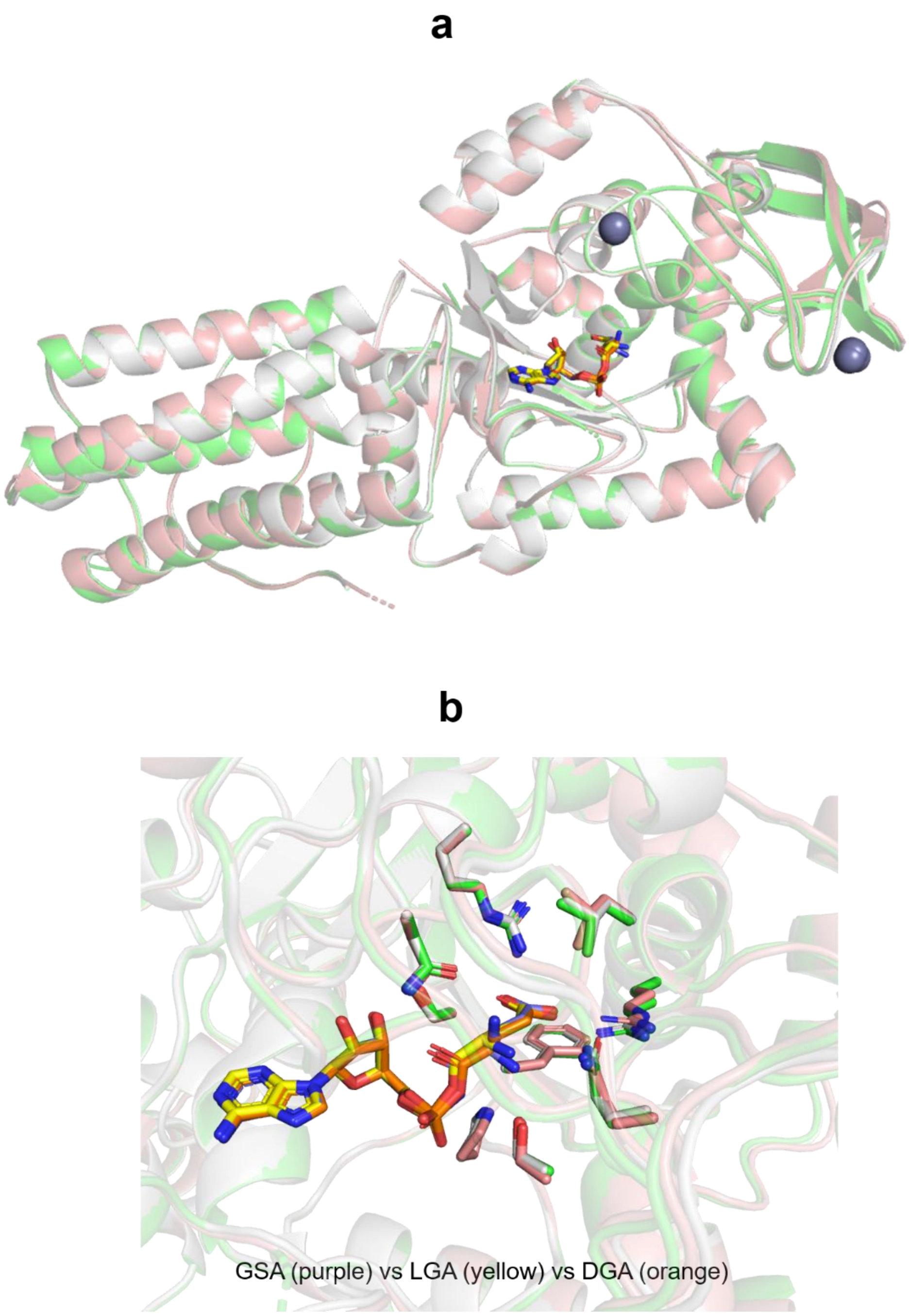

**Supplementary Fig 5.** Superposition of L-glutamyl-sulfamoyl-adenosine (GSA)-bound (salmon), D-glutamyl-adenylate (DGA)-bound (green), and L-glutamyl-adenylate (LGA)-bound (light gray) PyrN-C crystal structures. (a) Overlay of the three complex structures. (b) Overlay of GSA, DGA and LGA and residues within 4 Å of the glutamyl unit.

**Supplementary Fig. 6**

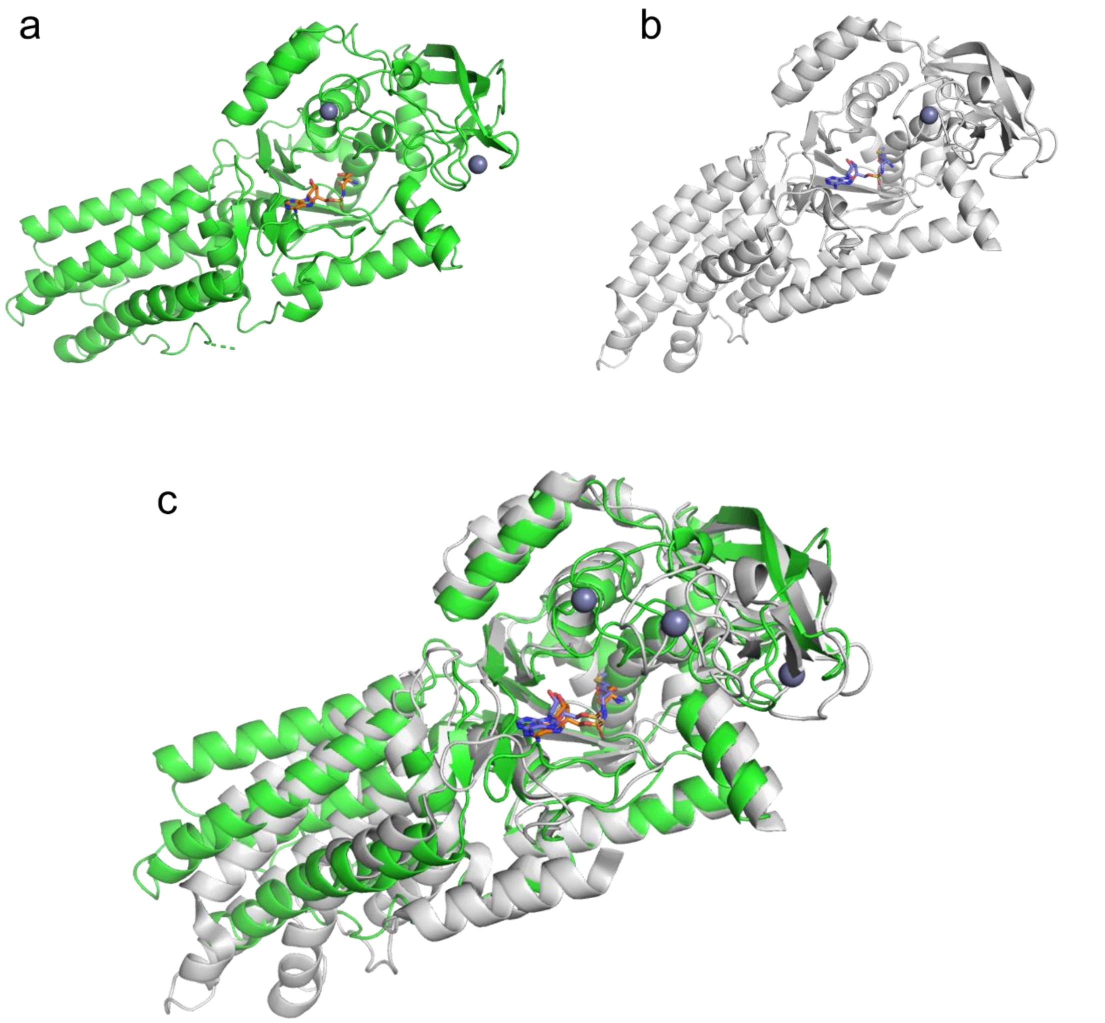

**Supplementary Fig 6.** Comparison of the crystal structures of PyrN-C with that of *E. coli* MetRS. (a) The structure of PyrN-C complexed with GSA solved in this study. The two structural zinc ions are displayed in gray sphere. (b) The structure of *E. coli* MetRS complexed with MSA (methionyl-sulphamoyl-adenosine) (PDB code: 1PFY).[35] Note: the *E. coli* MetRS contains single zinc knuckle in the CP domain. (c) Superposition of PyrN-C (green) and *E. coli* MetRS (light gray).

**Supplementary Fig. 7**

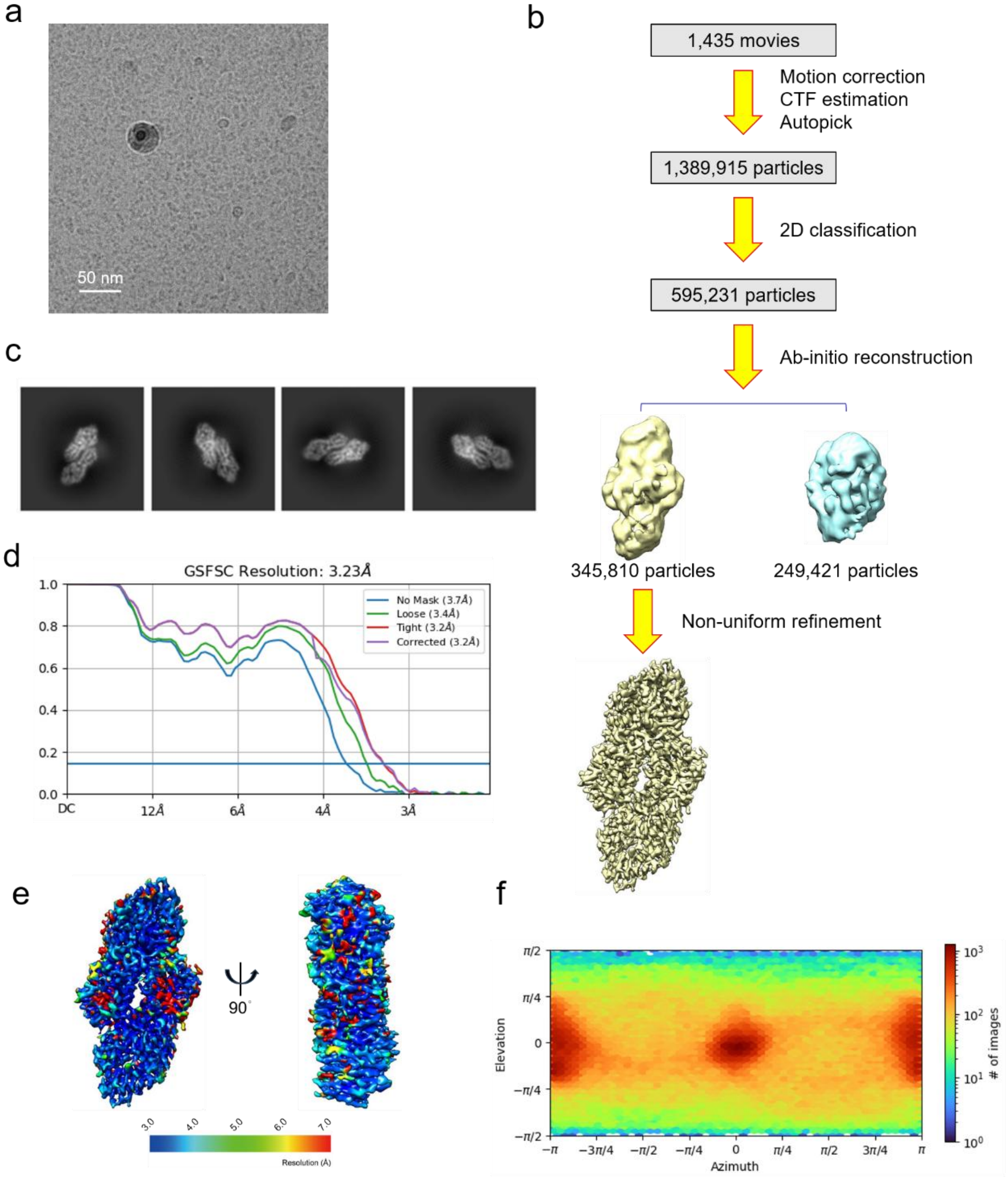

**Supplementary Fig 7.** Cryo-EM single-particle analysis of full-length PyrN. (a) A cryo-EM micrograph of PyrN. (b) The cryo-EM data processing pipeline. (c) The 2D averages of PyrN. (d) The Fourier shell correlation (FSC) curves. (e) The local resolution of the final map. (f) The orientation distribution of particles.

Note: 2D classification and *ab initio* reconstruction revealed that the particles exhibit C2 symmetry, indicating that full-length PyrN forms a dimer in solution. This finding is consistent with the dimeric state observed by gel filtration chromatography (Supplementary Fig. 8). Non-uniform refinement resulted in a reconstruction at 3.23 Å resolution. As the N-terminal domain was not resolved in the cryo-EM map, subsequent structural analyses focused on higher-resolution crystal structures.

#### Supplementary Fig. 8

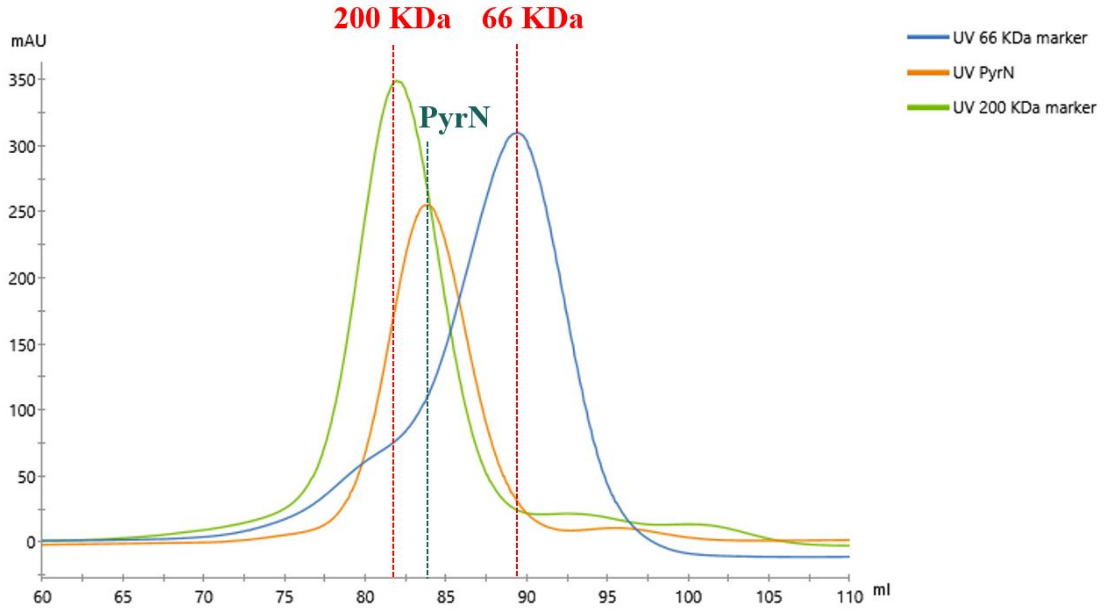

**Supplementary Fig 8.** Gel filtration analysis of PyrN (full-length). Calculated molecular weight for monomeric $His_6$-PyrN (full-length) is 74.9 kDa. The results indicated that the full-length didomain PyrN is a homodimer in solution. Note: Protein markers used for comparison are bovine serum albumin (66 kDa) and β-amylase from sweet potato (200 kDa).

**Supplementary Fig. 9**

**(a)**

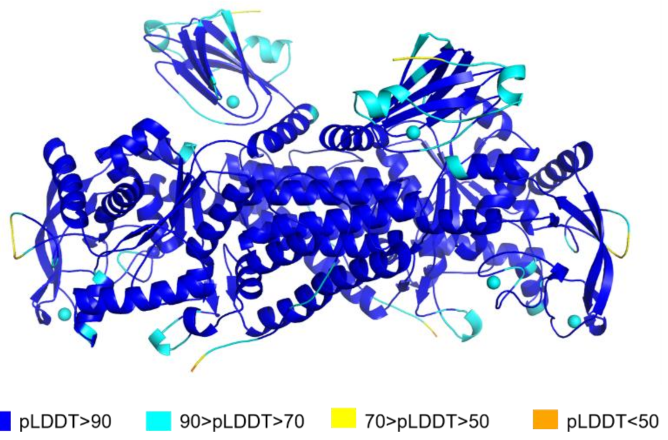

**(b)**

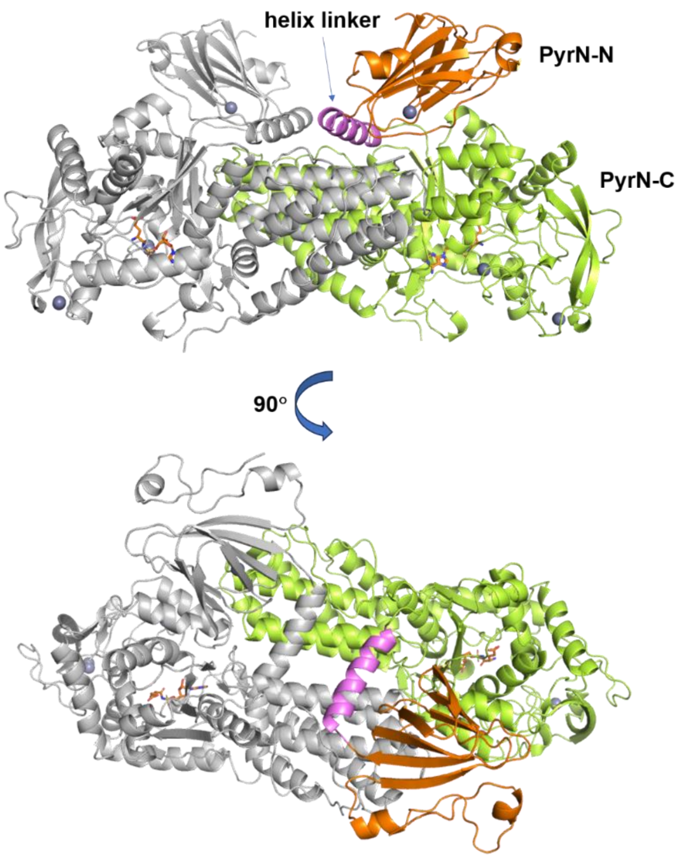

**Supplementary Fig 9.** AlphaFold-predicted structure of the full-length PyrN dimer. (a) Predicted Local Distance Difference Test (pLDDT) values for PyrN dimer generated by AlphaFold3[36]. (b) Domain organization of the AlphaFold-predicted structure of PyrN. PyrN-N (cupin) is shown in orange, PyrN-C is shown in lemon, and the helix linker is shown in pink (One subunit is colored, whereas the other subunit is displayed in grey).

**Supplementary Fig. 10**

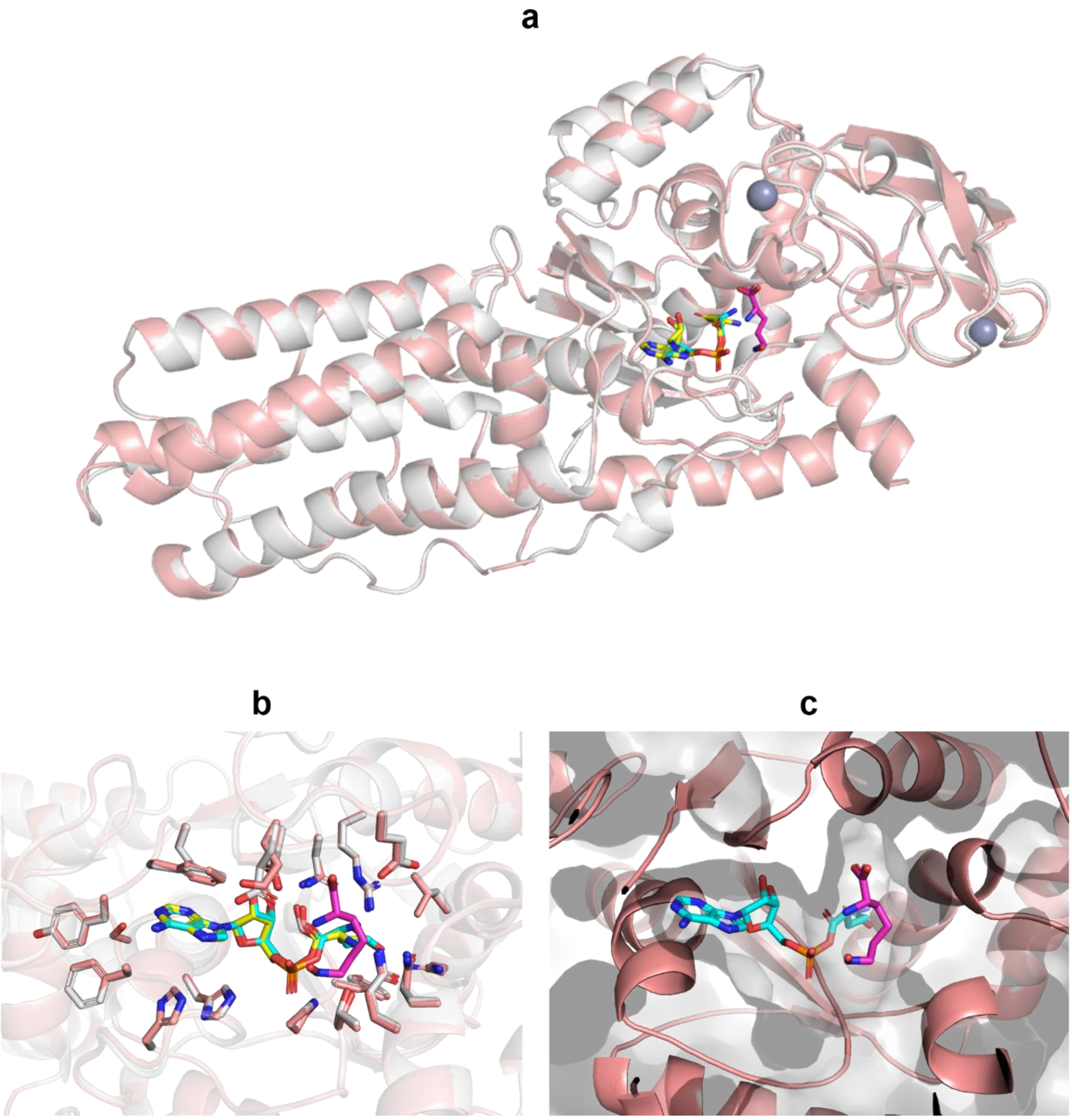

**Supplementary Fig 10.** Superposition of L-glutamyl-adenylate (LGA)-bound (light gray) and LGA-**1** bound (wheat) PyrN-C crystal structures. LGA is shown in yellow stick in the LGA-PyrN-C structure, and **1** is shown in magenta in the LGA-**1**-PyrN-C structure. (a) Overlay of the two complex structures. (b) Overlay of LGA, LGA and **1**, and active site residues within 4 Å. (c) Active site cavity of the LGA-**1**-PyrN-C complex structure.

**Supplementary Fig. 11**

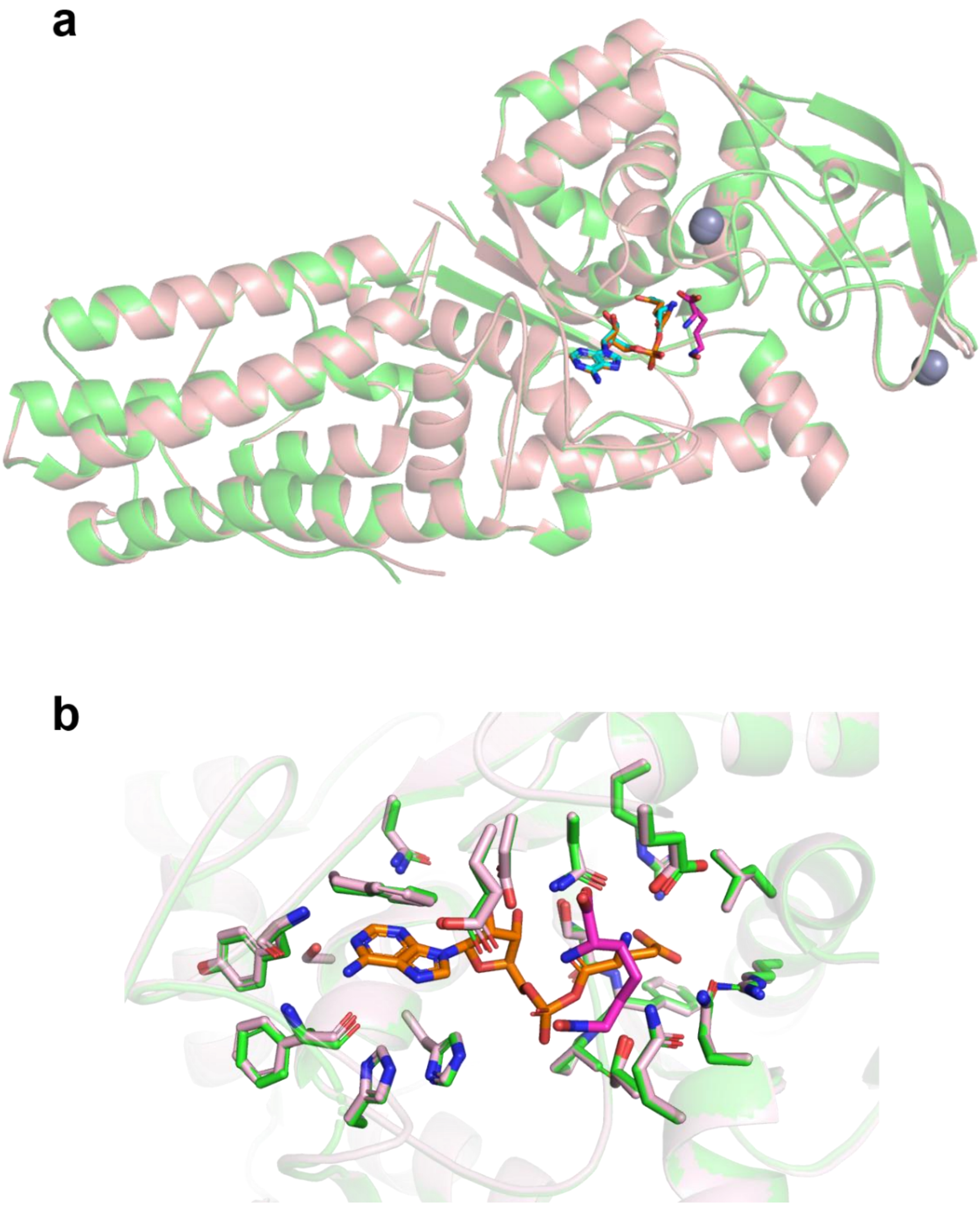

**Supplementary Fig. 11** Superposition of D-glutamyl-adenylate (DGA)-bound (green) and LGA-**1** bound (wheat) PyrN-C crystal structures. DGA is shown in orange stick in the DGA-PyrN-C structure, and **1** is shown in magenta in the LGA-**1**-PyrN-C structure.

**Supplementary Fig. 12**

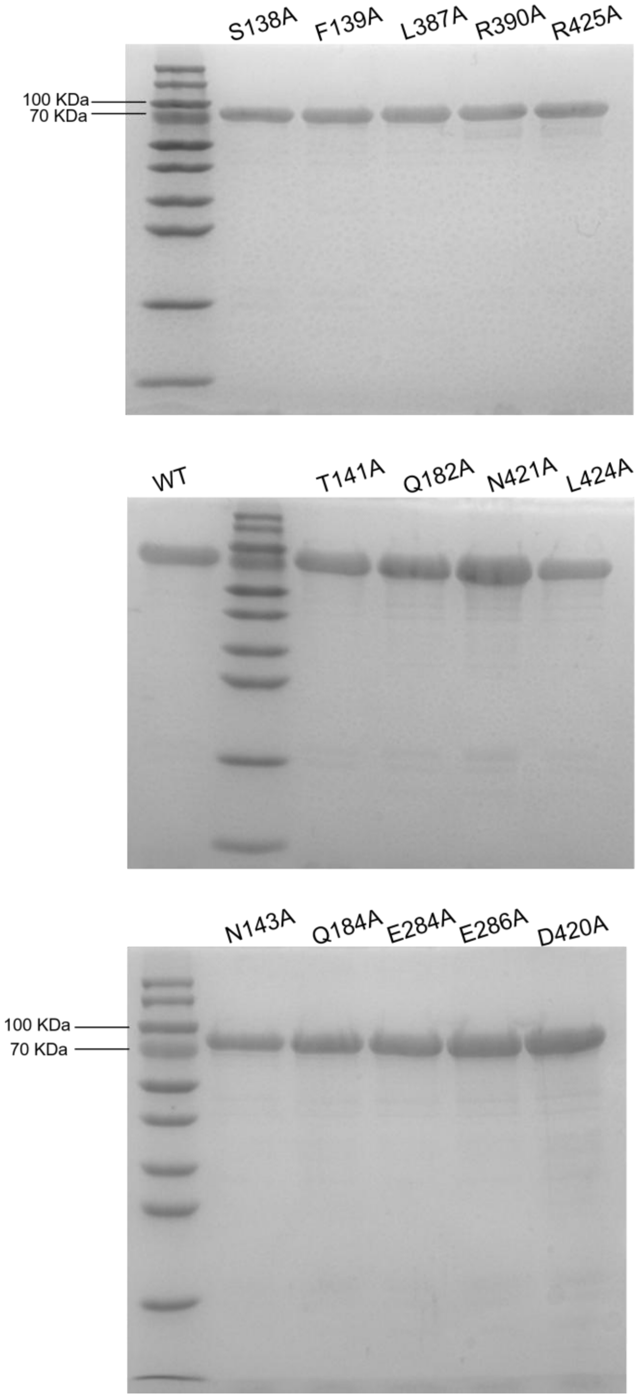

**Supplementary Fig. 12** The SDS-PAGE gel image of PyrN and its variants. Note: calculated molecular weight for $His_6$-PyrN is 74.9 kDa.

**Supplementary Fig. 13**

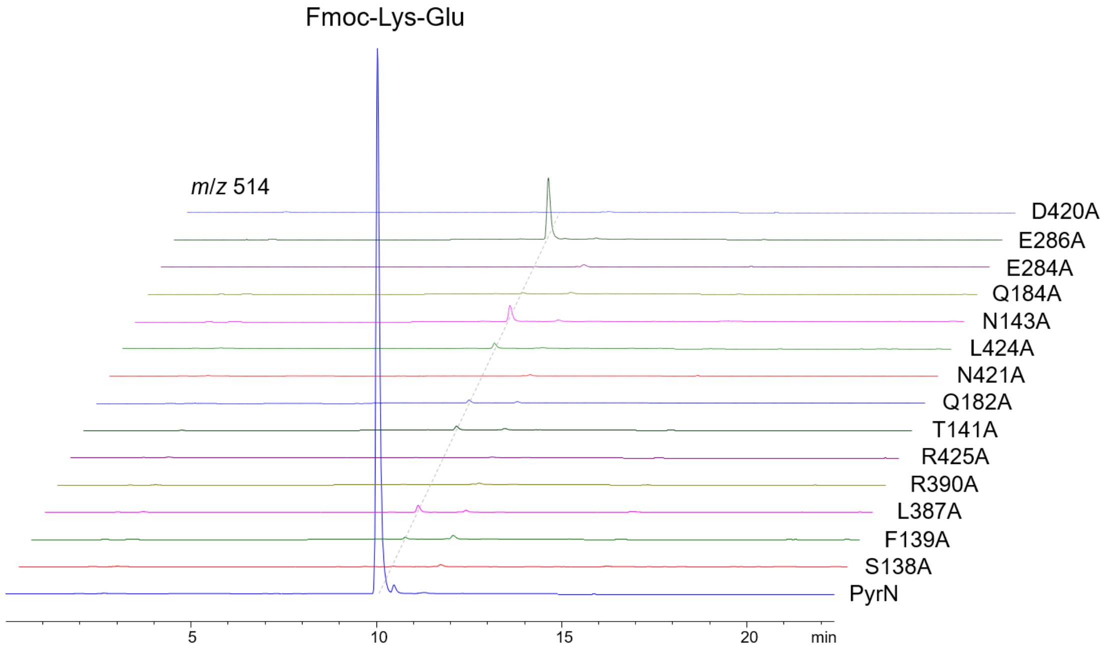

**Supplementary Fig 13.** LC-MS analysis of the in vitro reaction mixtures of PyrN and its variants after pre-column Fmoc-Cl derivatization. The extracted ion chromatographs (EIC = *m*/*z* 514) for the $[M+H]^+$ ion of Fmoc-Lys-Glu were displayed.

**Supplementary Fig. 14**

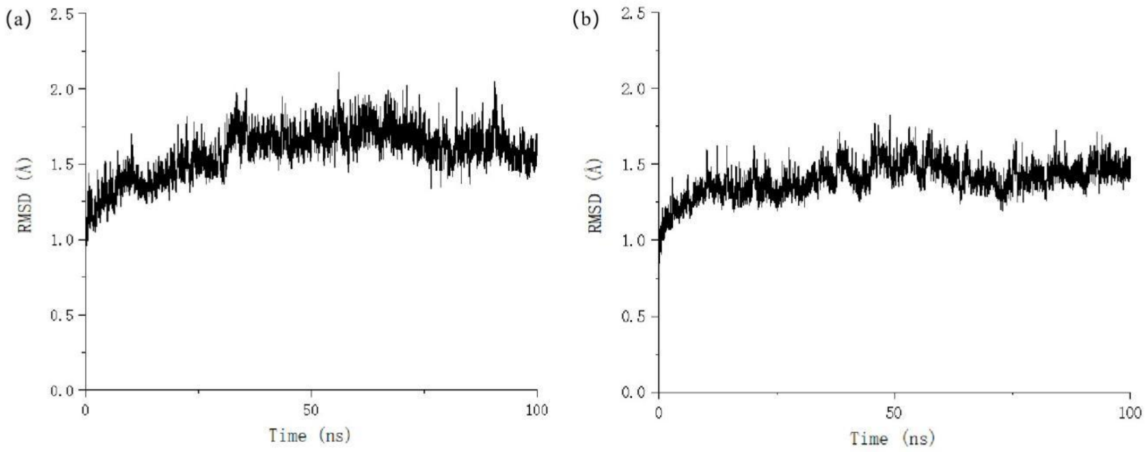

**Supplementary Fig 14.** Time evolution of the root mean square deviations (RMSD) for the protein backbone and substrate $N^6$-OH-L-Lys (**1**) of (a) zwitterionic form, (b) neutral form.

Note: it is important to note that the reaction mechanism between hydroxylamine analogues and ester groups has long been controversial.[2] It is generally accepted that three potential hydroxylamine forms exist: (1) the zwitterionic form, (2) the neutral form, and (3) the neutral form undergoing proton transfer to yield the zwitterionic form. The third scenario can be ruled out due to the prohibitive strain associated with disrupting the N-O-H three-membered ring.

**Supplementary Fig. 15**

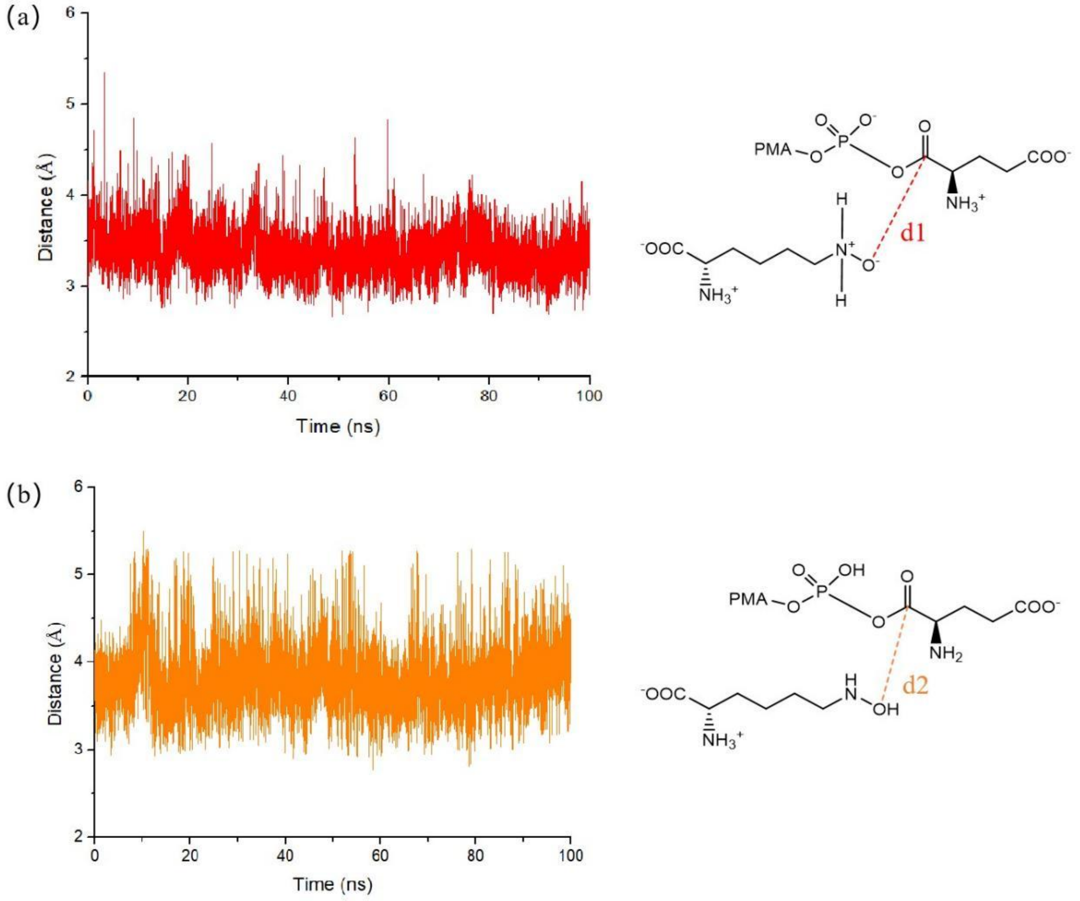

**Supplementary Fig 15.** Distance fluctuation between the hydroxylamine O atom of $N^6$-OH-L-Lys (**1**) and the ester carbonyl C atom of D-Glu-AMP during the 100 ns MD simulation. (a) $N^6$-OH-L-Lys (**1**) in zwitterionic form denote as d1 (in red). (b) $N^6$-OH-L-Lys (**1**) in neutral form denote as d2 (in orange)

**Supplementary Fig. 16**

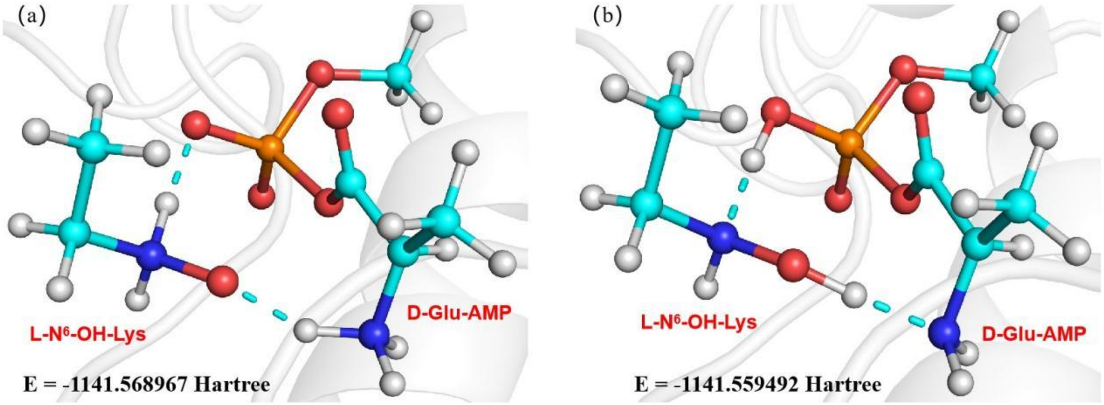

**Supplementary Fig 16.** QM (B3LYP-D3/B2) calculated absolute energy for the optimized reactant complex in (a) zwitterionic form, (b) neutral form.

#### Supplementary Fig. 17

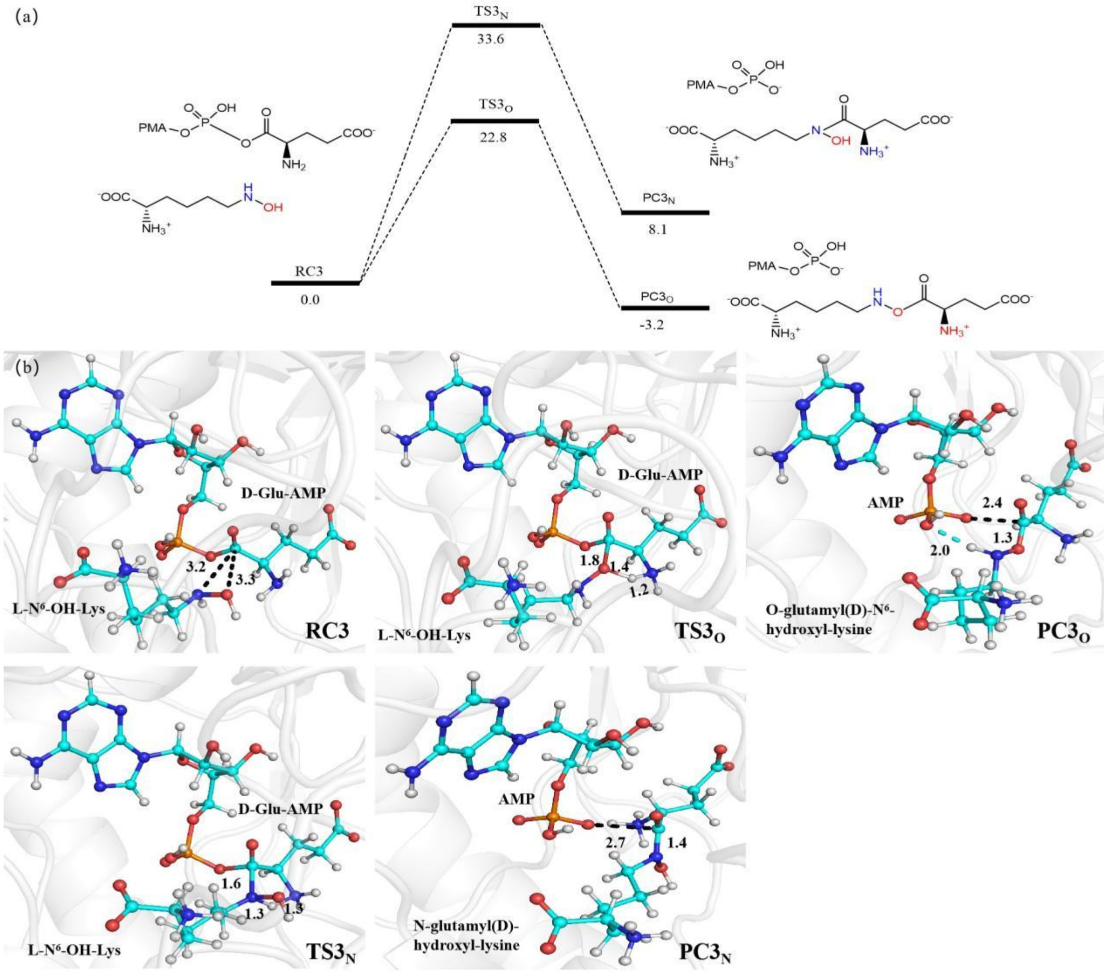

**Supplementary Fig 17.** (a) QM(B3LYP-D3/B2)/MM-calculated energy profile (in kcal mol$^{-1}$) for the oxygen nucleophilic attack vs nitrogen nucleophilic attack of substrate D-Glu-AMP and $N^6$-OH-L-Lys (**1**) in neutral form. The ZPEs are included in the relative energies. (b) QM(B3LYP-D3/B1)/MM-optimized structures of key species involved in the reaction. Key distances are given in Å.

**Supplementary Fig. 18**

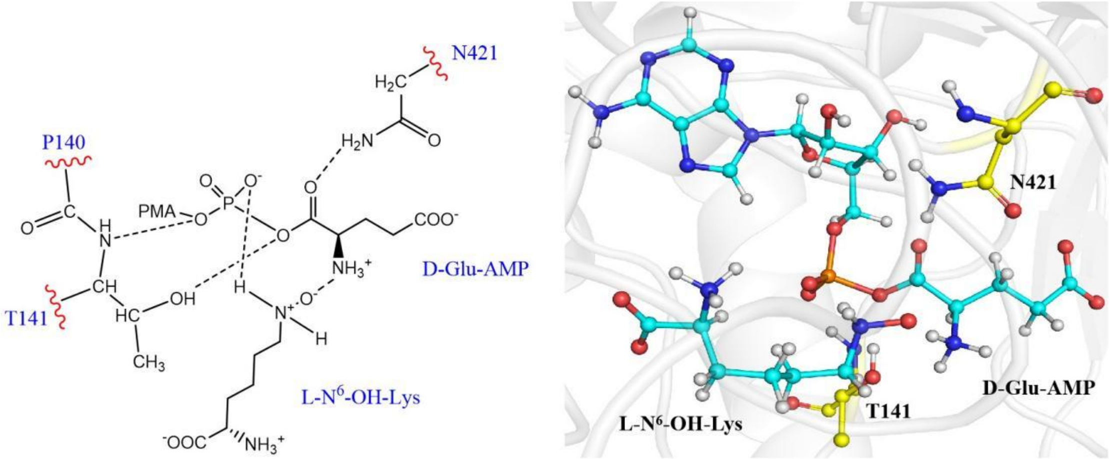

**Supplementary Fig 18.** QM regions used in QM/MM calculations.

**Supplementary Fig. 19**

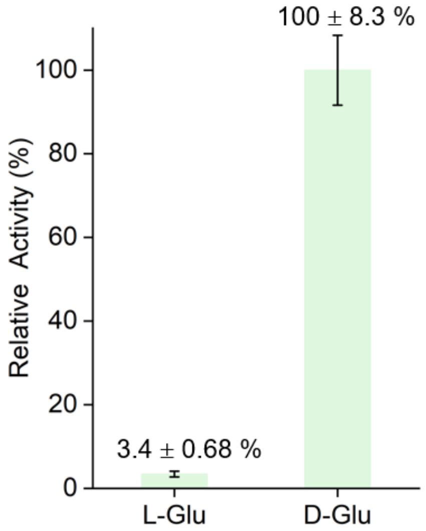

**Supplementary Fig 19.** Relative activities of PyrN toward L-Glu and D-Glu (2 mM each) determined using a pyrophosphate assay. Pyrophosphate released upon glutamyl-AMP formation was quantified by following a standard protocol[37].

#### Supplementary Fig. 20

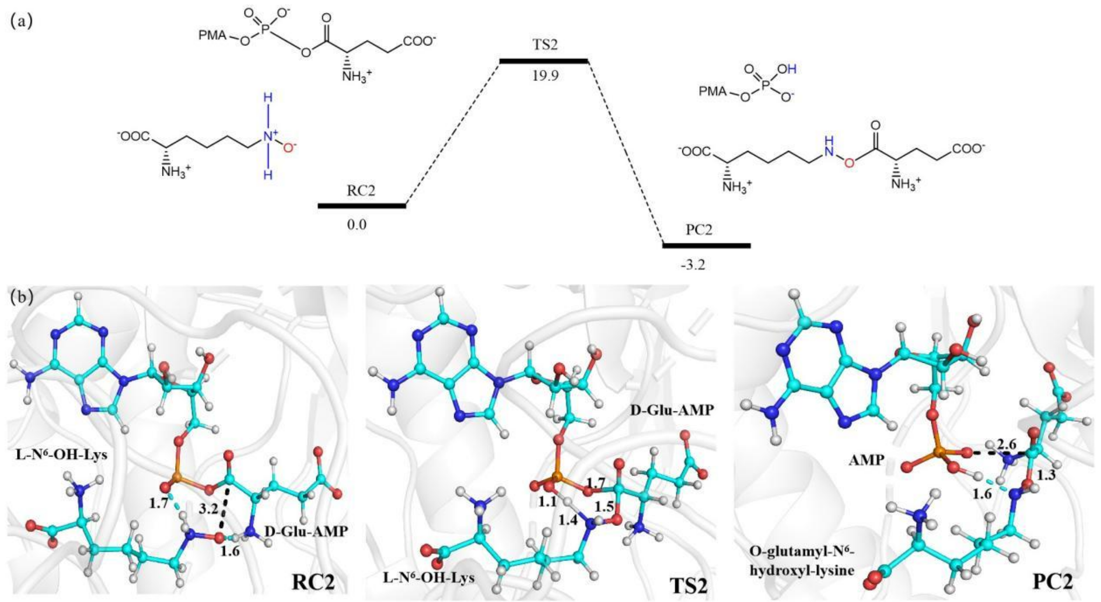

**Supplementary Fig 20.** (a) QM(B3LYP-D3/B2)/MM-calculated energy profile (in kcal mol$^{-1}$) for the oxygen nucleophilic attack of substrate L-Glu-AMP and $N^6$-OH-L-Lys (**1**) in zwitterionic form. The ZPEs are included in the relative energies. (b) QM(B3LYP-D3/B1)/MM-optimized structures of key species involved in the reaction. Key distances are given in Å.

**Supplementary Fig. 21**

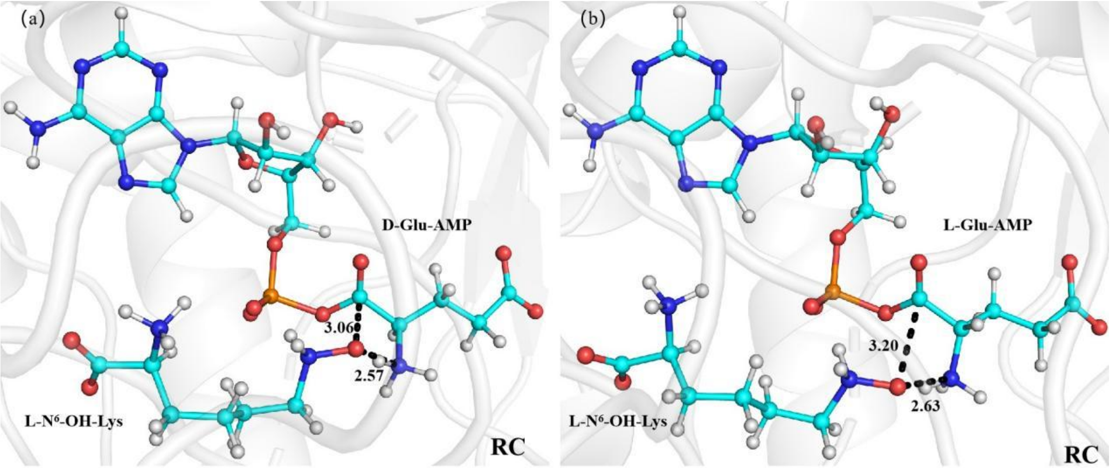

**Supplementary Fig 21.** QM(B3LYP-D3/B1)/MM-optimized structures of reactant complex in (a) D configuration, (b) L configuration. Key distances are given in Å.

#### Supplementary Fig. 22

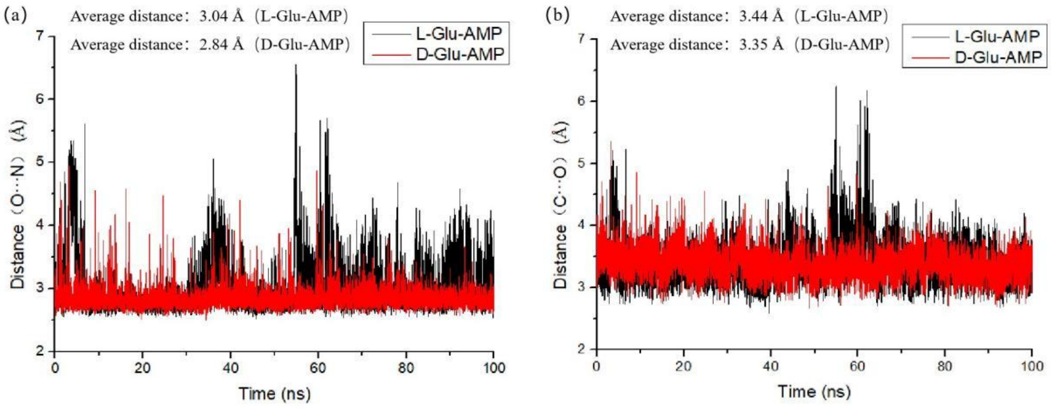

**Supplementary Fig. 22** The fluctuation of (a) O-N, (b) C-O distances in D-configured or L-configured substrate during the 100ns MD simulation.

**Supplementary Fig. 23**

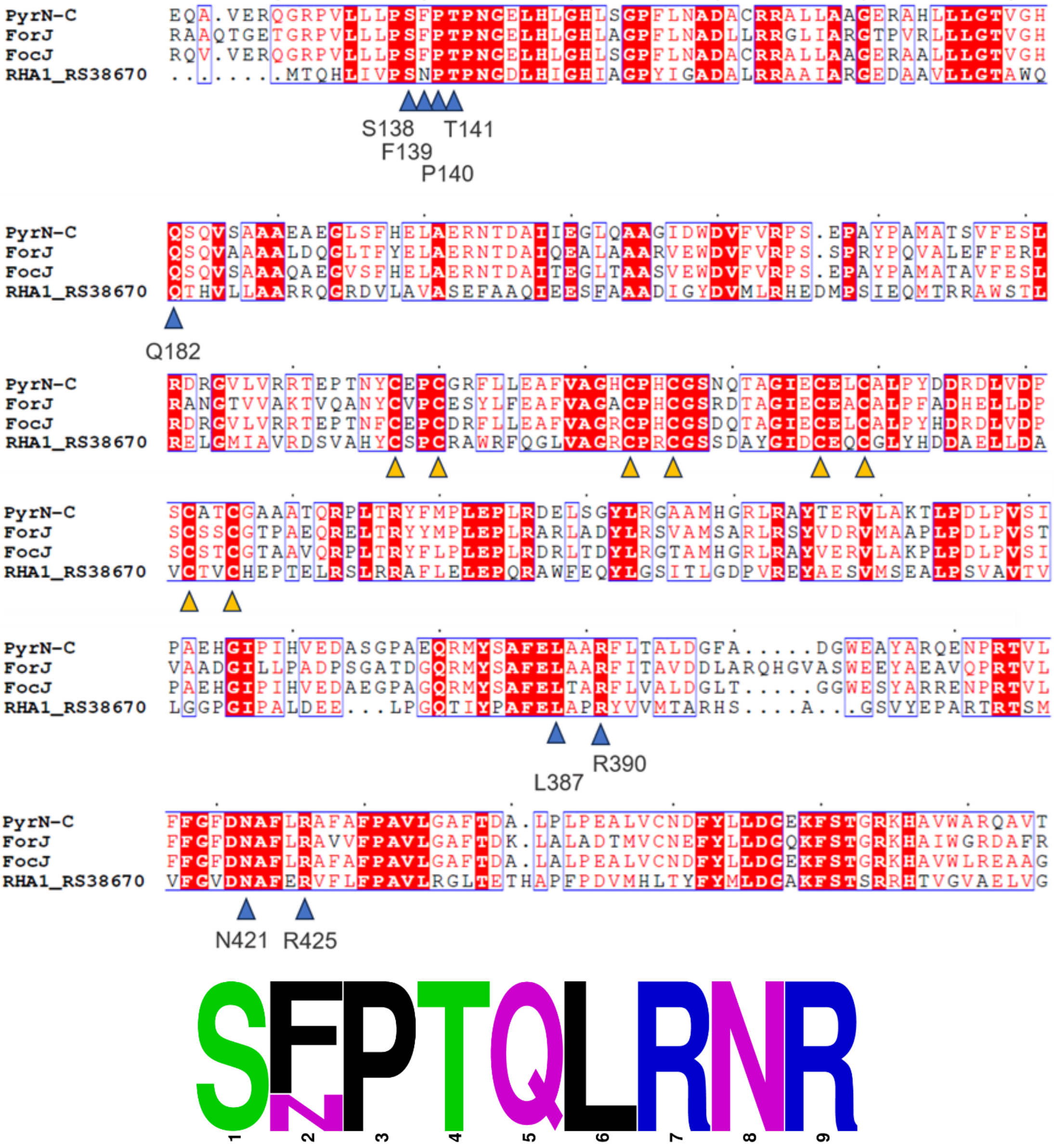

**Supplementary Fig. 23** Sequence alignment of the PyrN-C catalytic core region with other available glutamate-utilizing enzymes. Residues within 5 Å of GSA in the complex structure with PyrN-C are indicated with blue triangles. The sequence logo of these site residues is display at the bottom. Conserved zinc-coordination cysteines are indicated with yellow triangles.

Note: ForJ and FocJ are the PyrN homologues from the biosynthetic gene clusters of formycin in *Streptomyces kaniharaensis* ATCC 21070 and *Nocardia interforma* ATCC 21072, respectively[38,39]. RHA1_RS38670 is a PyrN homologue from *Rhodococcus jostii* RHA1[6].

**Supplementary Fig. 24**

a

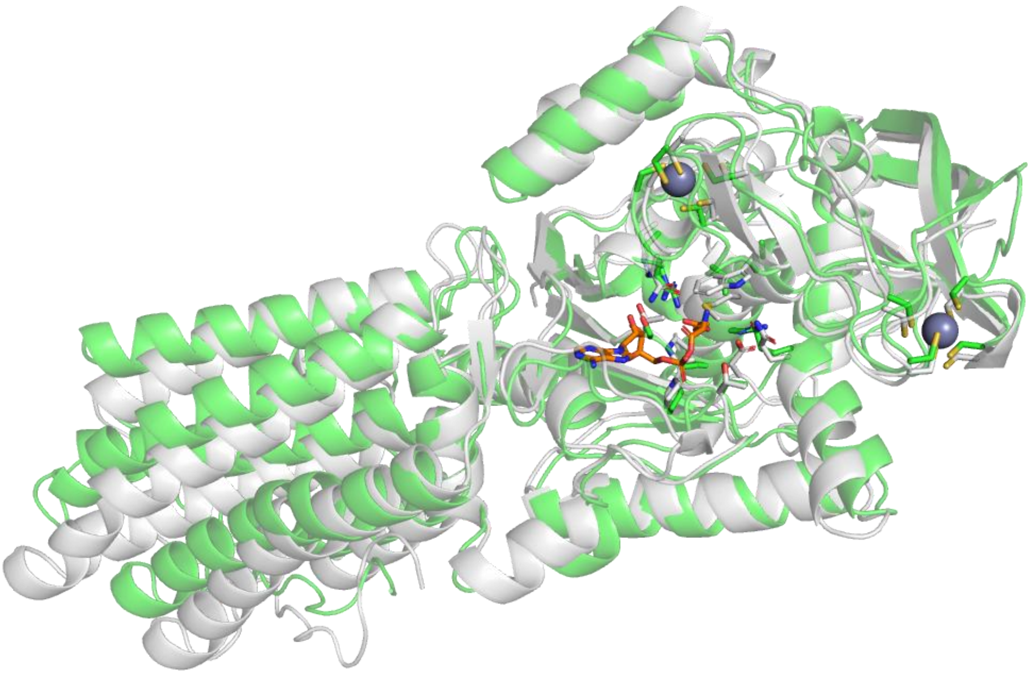

b

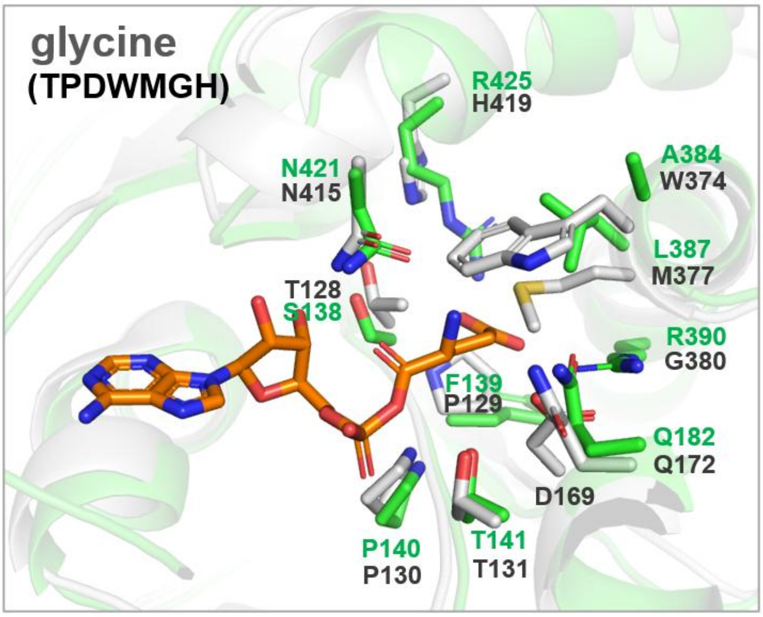

**Supplementary Fig. 24** Comparison of the amino acid binding site residues between PyrN and glycine-utilizing enzyme. (a) Superposition of the AlphaFold-predicted structure of Tri28-C (gray) onto the co-crystal structure of PyrN-C with DGA (green). DGA is shown in orange stick, and zinc ions are displayed in gray sphere. (b) Comparison of the amino acid substrate binding site residues between PyrN and glycine-utilizing Tri28. Based on our specificity code system, the 7-letter code for Tri28 is 'TPDWMGH'.

**Supplementary Fig. 25**

a

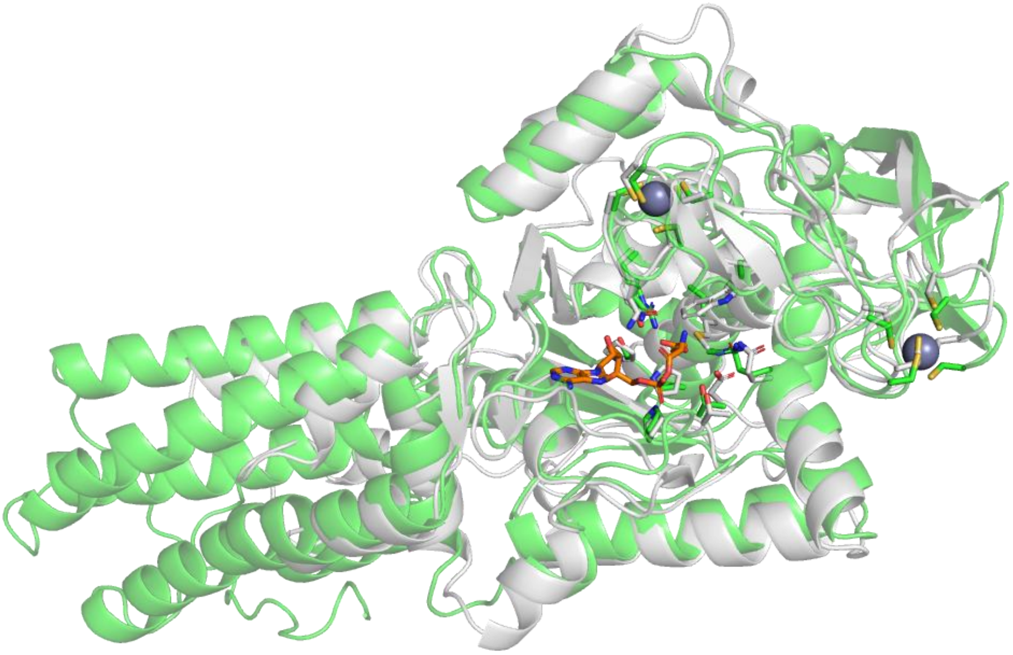

b

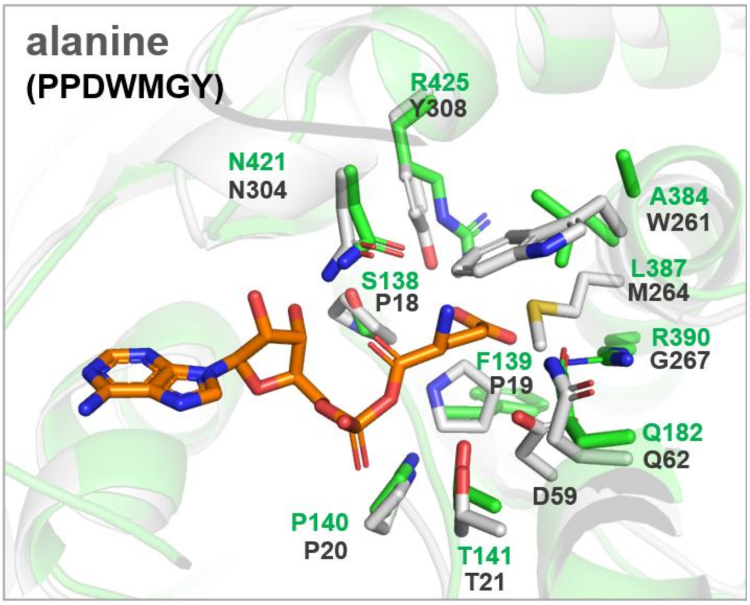

**Supplementary Fig. 25** Comparison of the amino acid binding site residues between PyrN and alanine-utilizing enzyme Afn8.[40] (a) Superposition of the AlphaFold-predicted structure of Afn8 (gray) onto the co-crystal structure of PyrN-C with DGA (green). (b) Comparison of the amino acid substrate binding site residues between PyrN and Afn8. Note: Afn8 is a standalone homolog of PyrN-C from the biosynthetic pathway of albofungin in *Streptomyces monomycini*, and it was demonstrated to accept alanine.[40] The specificity code for Afn8 is ‘PPDWMGY’.

**Supplementary Fig. 26**

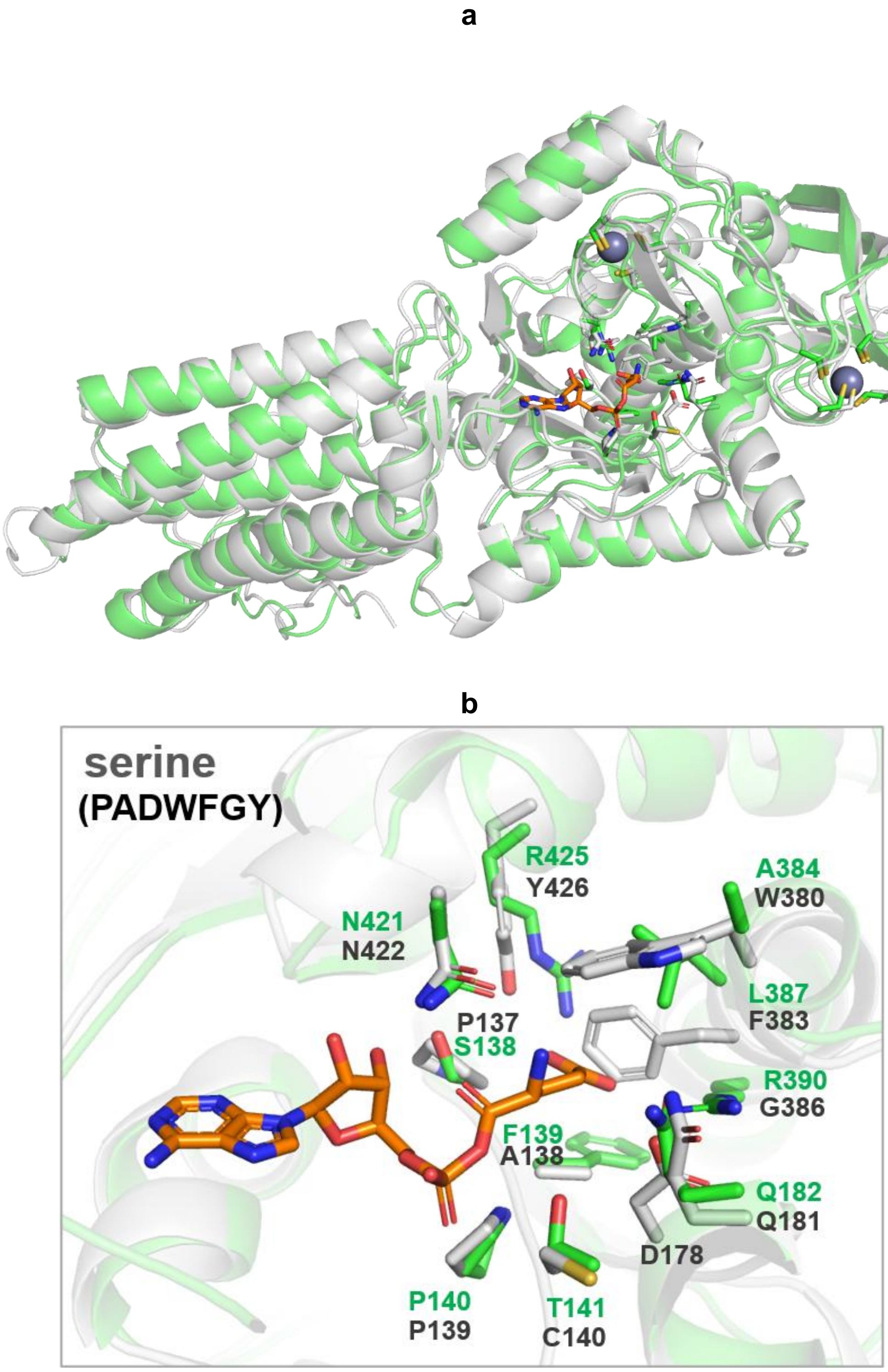

**Supplementary Fig. 26** Comparison of the amino acid binding site residues between PyrN and serine-utilizing enzyme SerHS (Uniprot code: A0A552E3D4). (a) Superposition of the AlphaFold-predicted structure of SerHS (gray) onto the co-crystal structure of PyrN-C with DGA (green). (b) Comparison of the amino acid substrate binding site residues between PyrN and SerHS. Note: SerHS is a di-domain homolog of PyrN from *Microcystis aeruginosa*, and was demonstrated to accept serine.[6] The specificity code for SerHS is 'PADWFGY'.

**Supplementary Fig. 27**

a

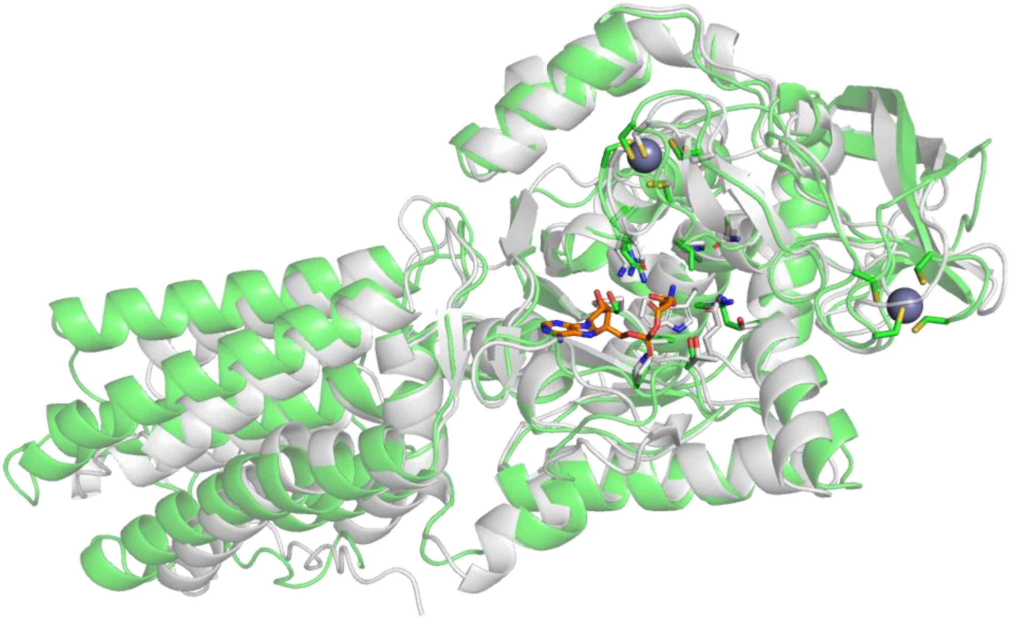

b

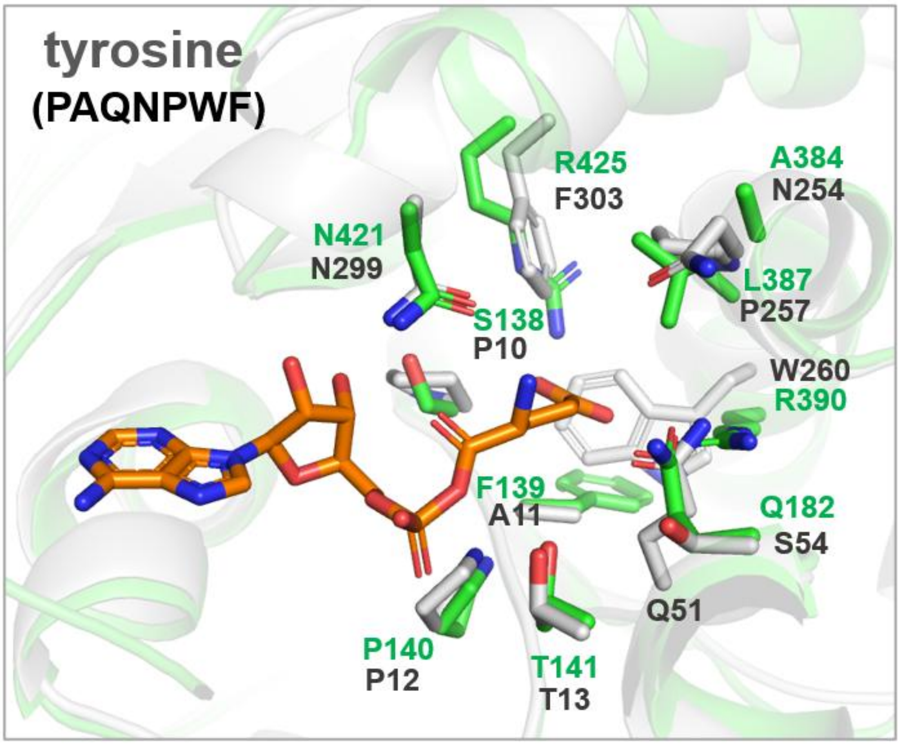

**Supplementary Fig. 27** Comparison of the amino acid binding site residues between PyrN and tyrosine-utilizing enzyme TyrHS (Uniprot code: D5UDN9). (a) Superposition of the AlphaFold-predicted structure of TyrHS (gray) onto the co-crystal structure of PyrN-C with GSA (green). (b) Comparison of the amino acid substrate binding site residues between PyrN and TyrHS. Note: TyrHS is a standalone homolog of PyrN-C from *Cellulomonas flavigena* and was demonstrated to accept tyrosine.[6] The specificity code for TyrHS is ‘PAQNPWF’.

**Supplementary Fig. 28**

a

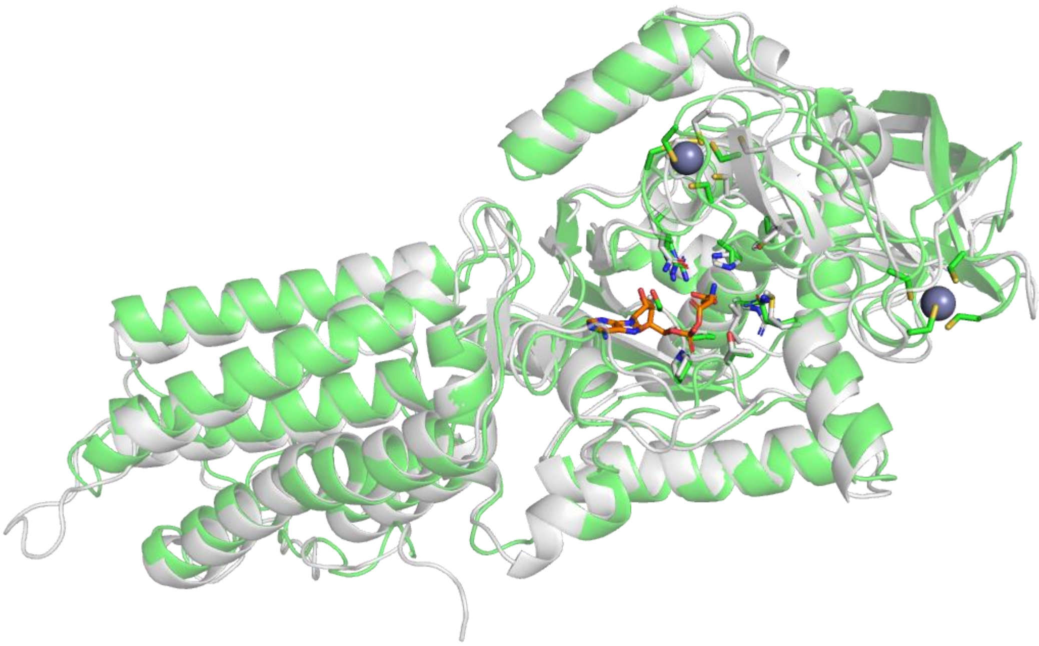

b

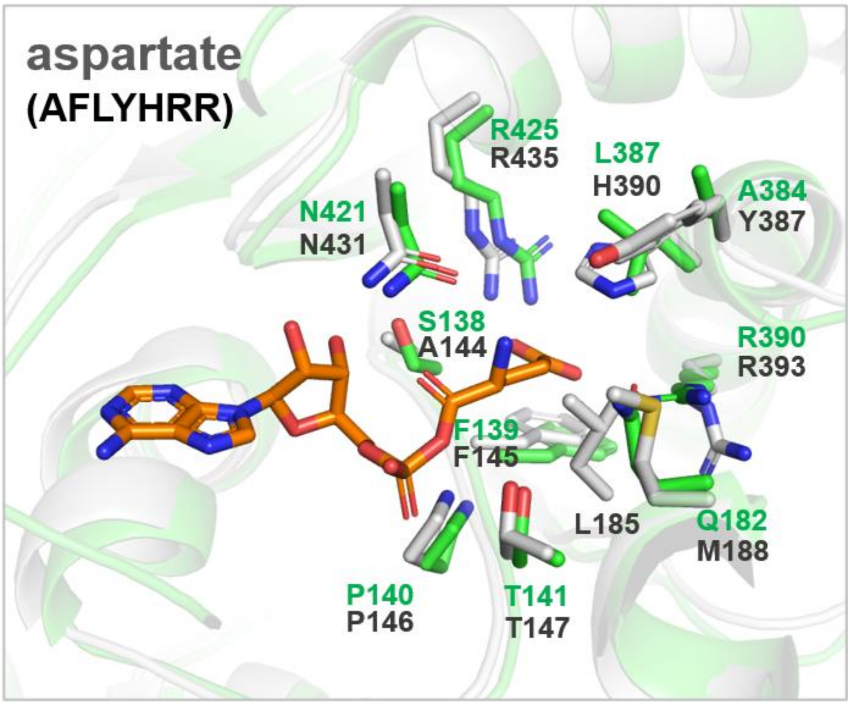

**Supplementary Fig. 28** Comparison of the amino acid binding site residues between PyrN and newly-identified aspartate-utilizing enzyme AspHS (NCBI: MCP4619998). (a) Superposition of the AlphaFold-predicted structure of AspHS (gray) onto the co-crystal structure of PyrN-C with GSA (green). (b) Comparison of the amino acid substrate binding site residues between PyrN and AspHS. Note: AspHS is a PyrN homolog from *Bradyrhizobium* sp. The specificity code for AspHS is 'AFLYHRR'.

**Supplementary Fig. 29**

**a**

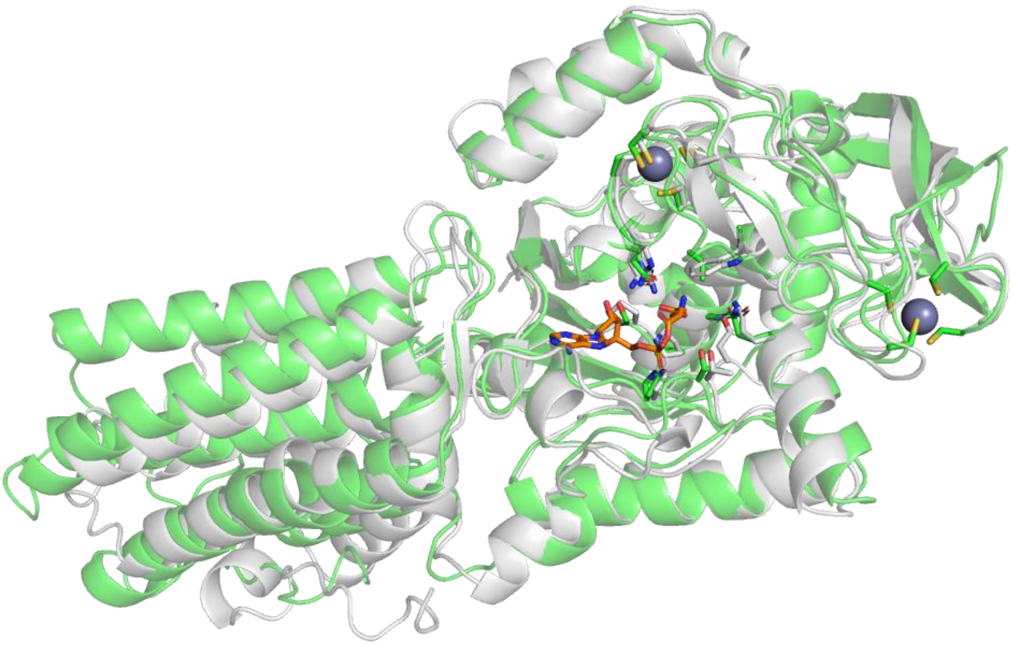

**b**

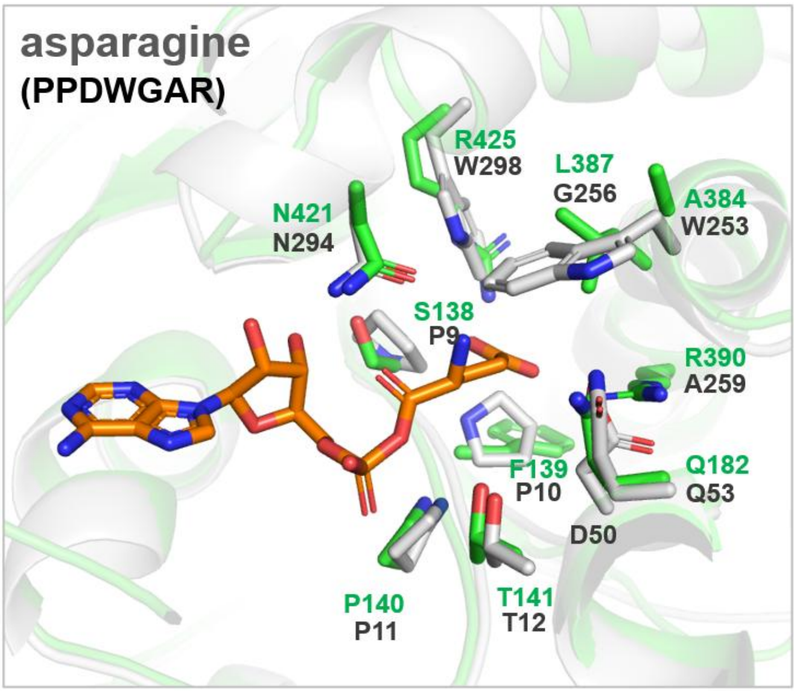

**Supplementary Fig. 29** Comparison of the amino acid binding site residues between PyrN and newly-identified asparagine-utilizing enzyme AsnHS (NCBI: WP_095402837). (a) Superposition of the AlphaFold-predicted structure of AsnHS (gray) onto the co-crystal structure of PyrN-C with GSA (green). (b) Comparison of the amino acid substrate binding site residues between PyrN and AsnHS. Note: AsnHS is a PyrN-C homolog from *Burkholderia ubonensis*. The specificity code for AsnHS is ‘PPDWGAW’.

**Supplementary Fig. 30**

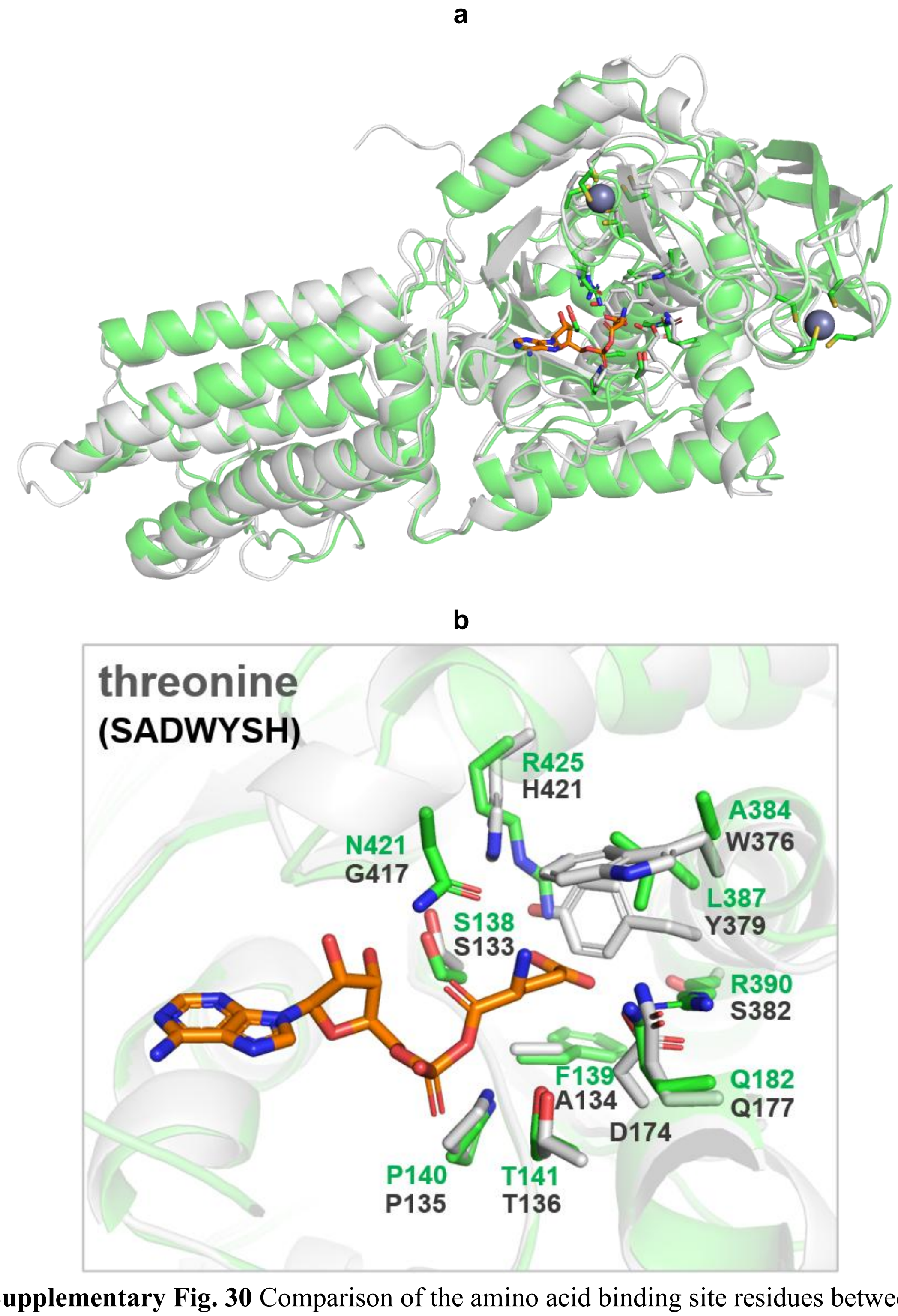

**Supplementary Fig. 30** Comparison of the amino acid binding site residues between PyrN and newly-identified threonine-utilizing enzyme ThrHS (NCBI: WP_119101482). (a) Superposition of the AlphaFold-predicted structure of ThrHS (gray) onto the co-crystal structure of PyrN-C with GSA (green). (b) Comparison of the amino acid substrate binding site residues between PyrN and ThrHS. Note: ThrHS is a tri-domain enzyme (PyrN-PCP homolog) from *Streptomyces sporangiiformans*. The specificity code for ThrHS is ‘SADWYSH’.

#### Supplementary Fig. 31

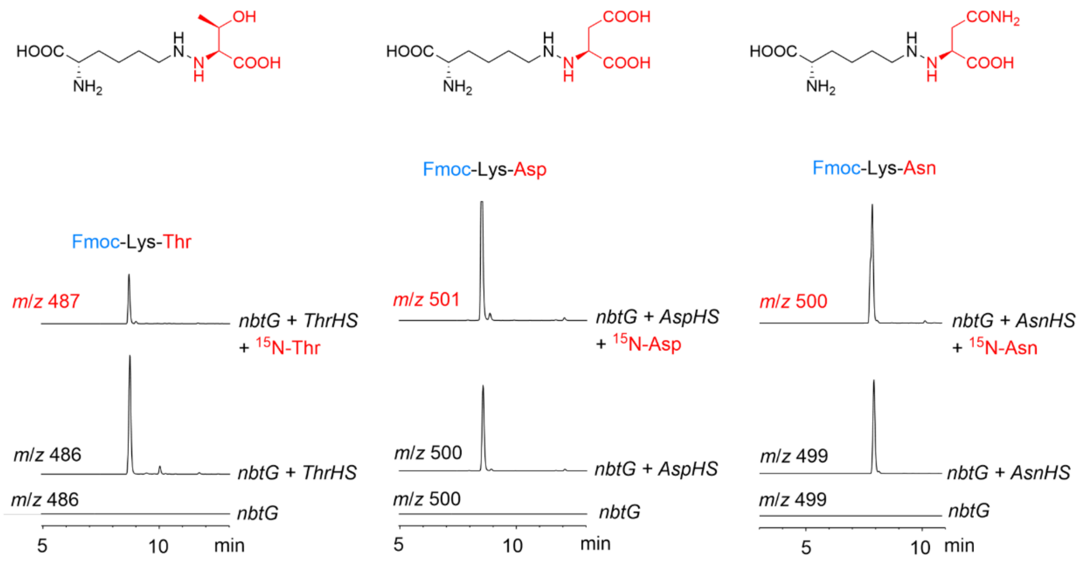

**Supplementary Fig. 31** Identification of new PyrN homologs that utilizes unreported amino acid substrate using an in vivo biotransformation assay. The EICs for the $[M+H]^+$ ions of Fmoc-derived hydrazine products are shown. When $^{15}N$-labeled amino acids were used, enriched isotopic peaks were detected (shown in red). Note: a threonine-utilizing hydrazine synthetase was also reported by another group during the preparation of this manuscript[41].

**Supplementary Fig. 32**

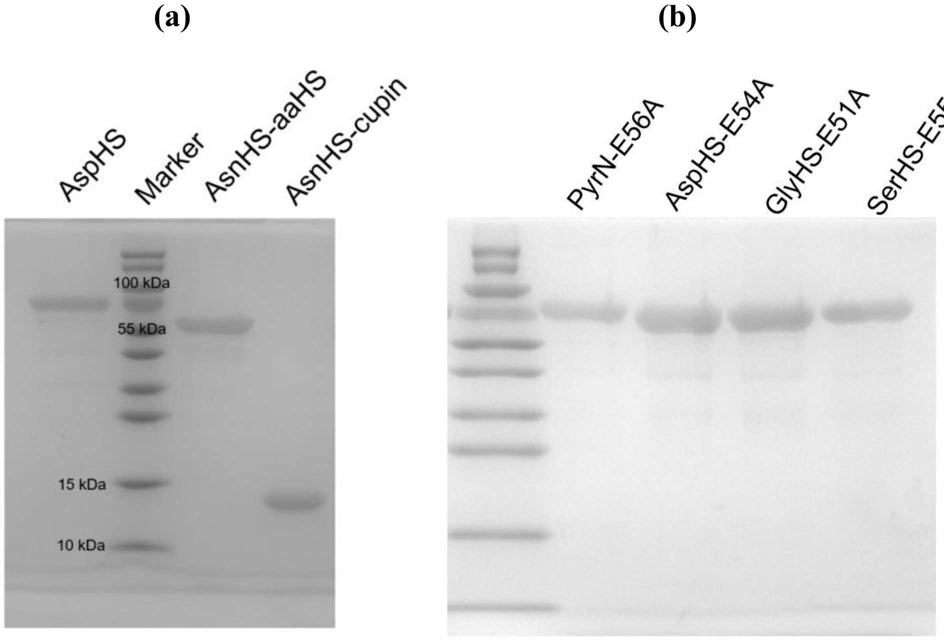

**Supplementary Fig 32.** The SDS-PAGE gel image of protein used in in vitro assays. (a) SDS-PAGE of isolated didomain AspHS from *Bradyrhizobium* sp. and standalone AsnHS-aaHS/AsnHS-cupin from *Burkholderia ubonensis*. Note: the NCBI accession code for AspHS is MCP4619998, and for AsnHS-aaHS/AsnHS-cupin are WP_095402837/WP_095402836. aaHS: aminoacyl-hydroxylase synthetase. (b) SDS-PAGE of hydrazine synthetase variants that carry an inactive cupin domain due to the mutation of an essential Glu residue[6]. These variants were used as negative control in the in vitro assays as indicated.

#### Supplementary Fig. 33

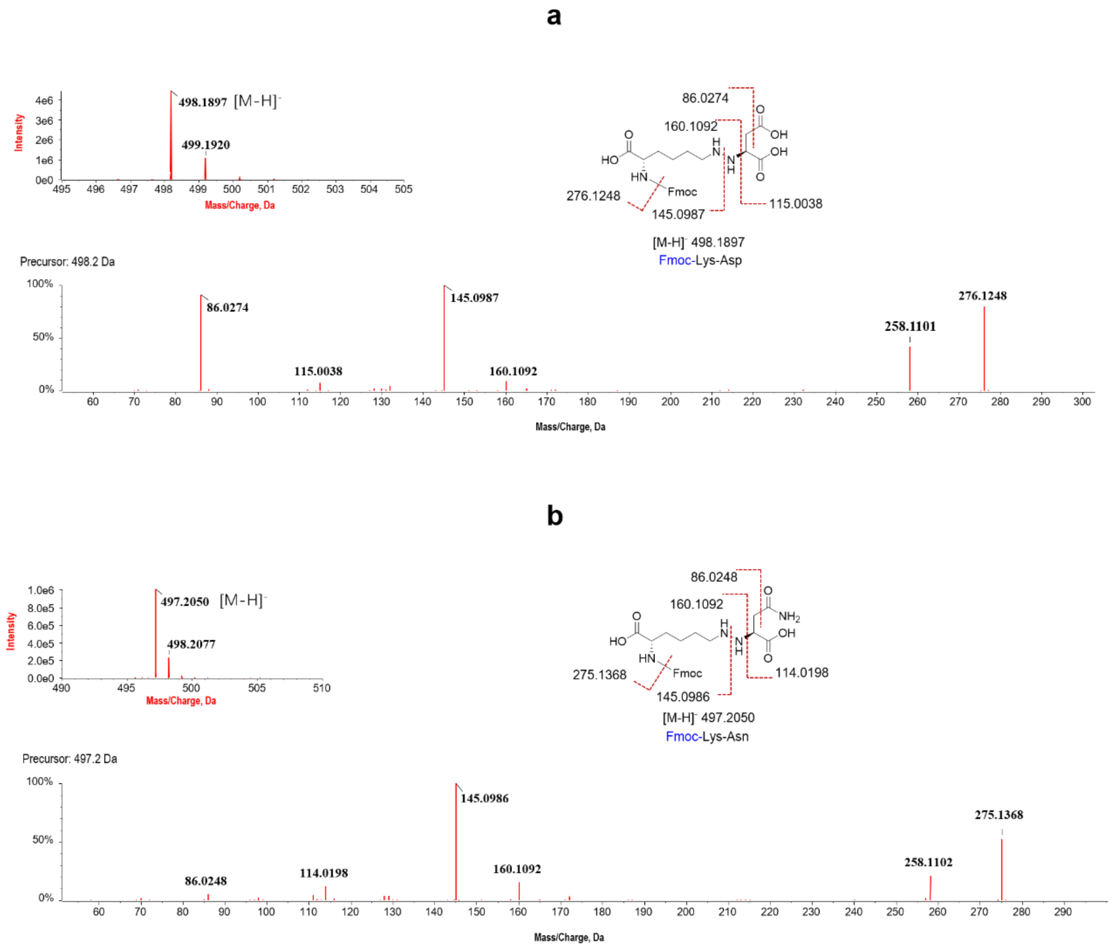

**Supplementary Fig. 33** LC-HR-MS/MS analysis (under negative mode) of the hydrazine products from newly-discoveried AspHS and AsnHS. Selected diagnostic fragment signals are indicated in the chemical structures of the corresponding products.

**Supplementary Fig. 34**

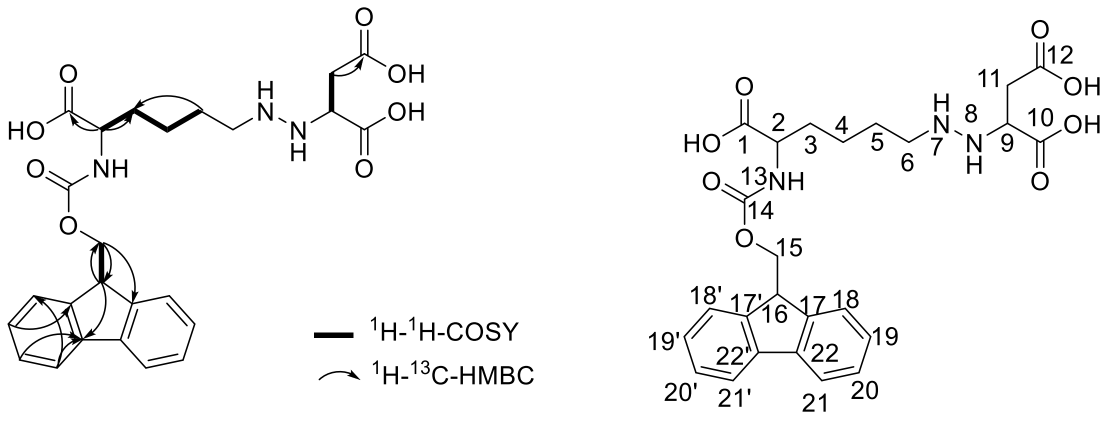

| NMR data of Compound Lys-Asp-Fmoc[a] | | | | | |
|---|---|---|---|---|---|
| NO. | $\delta_H$ (*J* in Hz) | $\delta_C$ | NO. | $\delta_H$ (*J* in Hz) | $\delta_C$ |
| 1 | - | 174.94 | 14 | - | 158.66 |
| 2 | 3.56 (1H, m) | 55.73 | 15 | 4.60 (2H, d, 5.4) | 68.49 |
| 3 | 1.73,1.84 (1H, m) | 31.64 | 16 | 4.27 (1H, d, 5.5) | 48.55 |
| 4 | 1.34 (2H, m) | 23.42 | 17, 17′ | - | 145.26 |
| 5 | 1.29 (2H, m) | 30.75 | 18, 18′ | 7.63 (2H, d, 7.5) | 125.86 |
| 6 | 3.19 (2H, m) | 27.88 | 19, 19′ | 7.33 (2H, tt, 7.4 1.4) | 128.26 |
| 9 | 3.81 (1H, brs) | 59.68 | 20, 20′ | 7.40 (2H, t, 7.4) | 128.89 |
| 10 | - | 174.03 | 21,21′ | 7.82 (2H, d, 7.5) | 121.09 |
| 11 | 2.57 (2H, brs) | 36.96 | 22,22′ | - | 142.77 |
| 12 | - | 175.46 | | | |

[a] $^{1}H$ (600 MHz) and $^{13}C$ (150 MHz) NMR Data in MeOD.

**Supplementary Fig. 34** Structural assignment of isolated Lys-Asp-Fmoc based on NMR analysis. 1D and 2D NMR spectra are shown in Supplementary Fig. 52.

**Supplementary Fig. 35**

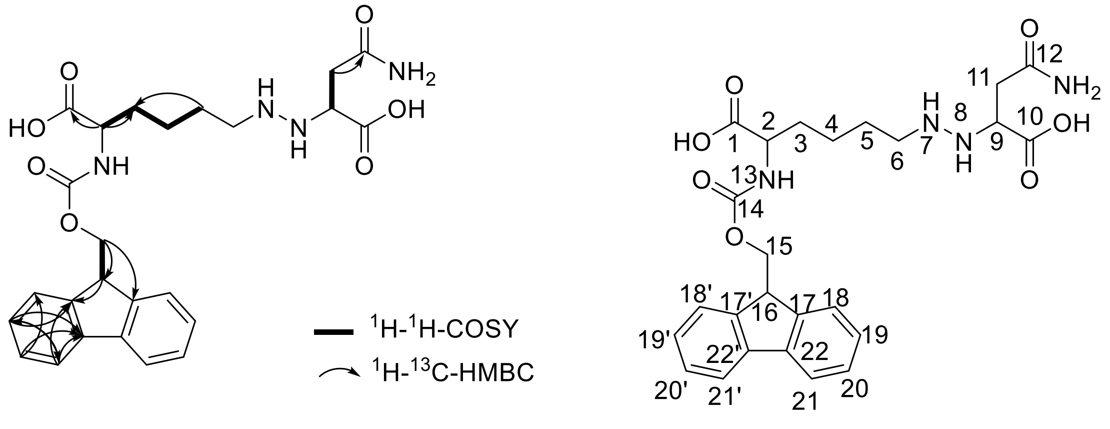

Lys-Asn-Fmoc

The NMR data for compound Lys-Asn-Fmoc.

| NMR data of Compound Lys-Asn-Fmoc[b] | | | | | |
|---|---|---|---|---|---|
| NO. | $\delta_H$ (*J* in Hz) | $\delta_C$ | NO. | $\delta_H$ (*J* in Hz) | $\delta_C$ |
| 1 | - | 174.10 | 14 | - | 158.78 |
| 2 | 3.56 (1H,m) | 55.41 | 15 | 4.60 (2H, m) | 68.49 |
| 3 | 1.72,1.84 (1H, brs) | 31.64 | 16 | 4.27 (1H, t, 5.6) | 48.94 |
| 4 | 1.34 (2H, brs) | 23.42 | 17, 17′ | - | 145.23 |
| 5 | 1.29 (2H, brs) | 30.75 | 18, 18′ | 7.61 (2H, d, 7.5) | 125.84 |
| 6 | 3.19 (2H, m) | 50.57 | 19, 19′ | 7.32 (2H, td, 7.4 1.1) | 128.25 |
| 9 | 3.81 (1H, m) | 60.09 | 20, 20′ | 7.40 (2H, t, 7.4) | 128.88 |
| 10 | - | 175.16 | 21,21′ | 7.81 (2H, d, 7.6) | 121.08 |
| 11 | 2.51 (2H, brs) | 37.53 | 22,22′ | - | 142.77 |
| 12 | - | 175.56 | | | |

b $^{1}$H (600 MHz) and $^{13}$C (150 MHz) NMR Data in MeOD.

**Supplementary Fig. 35** Structural assignment of isolated Lys-Asn-Fmoc based on NMR analysis. 1D and 2D NMR spectra are shown in Supplementary Fig. 53.

**Supplementary Fig. 36**

a

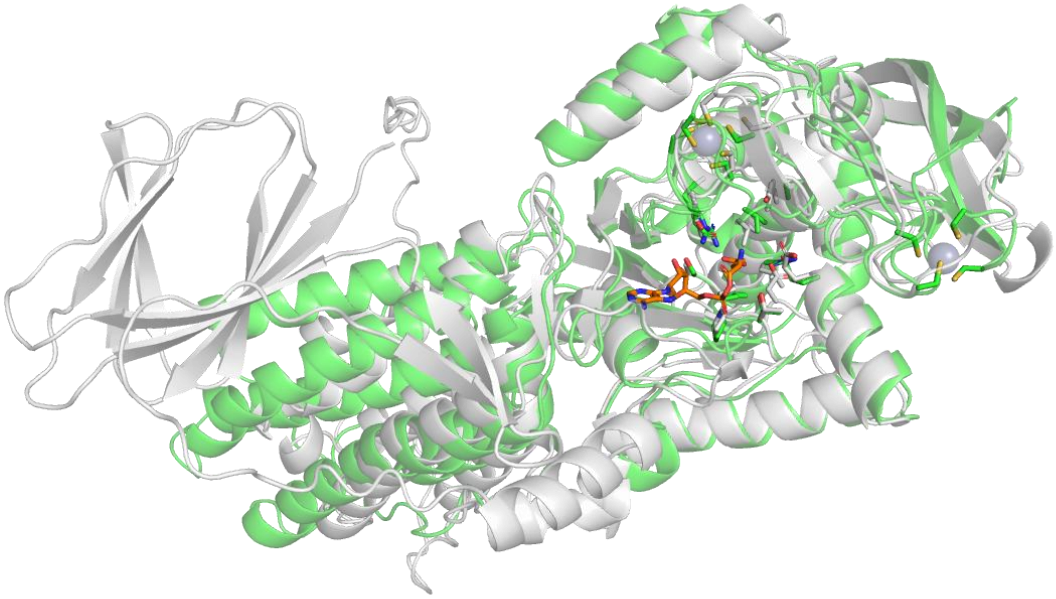

b

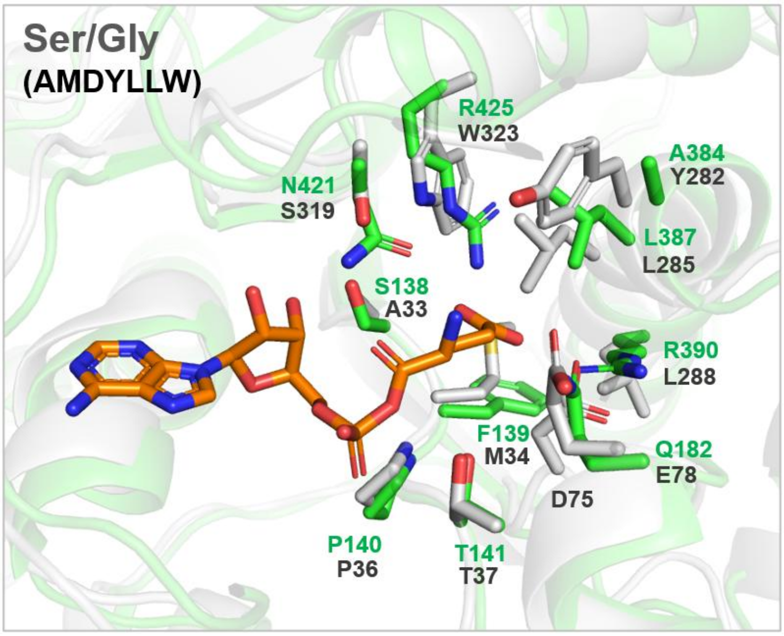

c

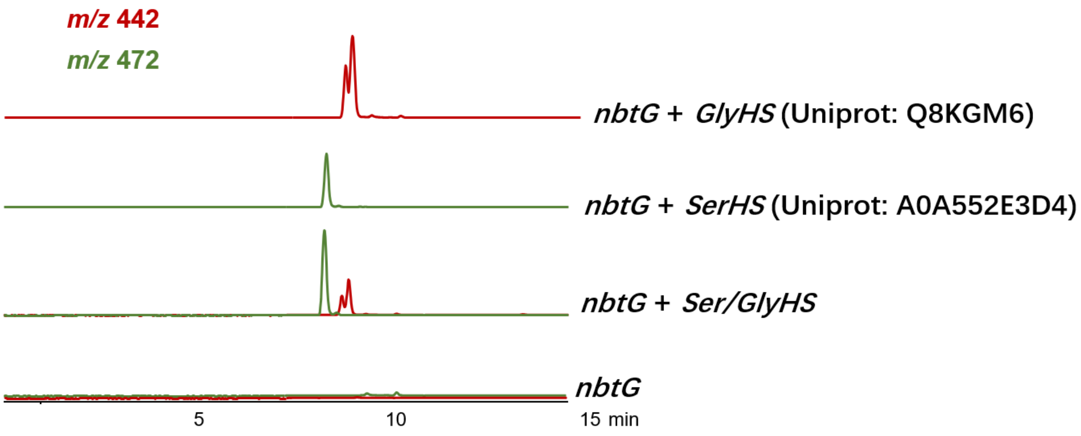

**Supplementary Fig. 36** Characterization of the newly-identified Ser/Gly-utilizing enzyme Ser/GlyHS (NCBI: RBL92032). (a) Superposition of the AlphaFold-predicted structure of Ser/GlyHS (gray) onto the co-crystal structure of PyrN-C with DGA (green). (b) Comparison of the amino acid substrate binding site residues between PyrN and Ser/GlyHS. Note: Ser/GlyHS is a PyrN homolog from *Chitinophaga flava*, with the cupin domain at C-terminus and the PyrN-C-like domain at N-terminus. The specificity code for Ser/GlyHS is 'AMDYLLW'. (c) Determination of the amino acid specificity of Ser/GlyHS using an in vivo biotransformation assay. The EICs for the $[M+H]^+$ ions (*m*/*z* 542, 572) of Fmoc-derived hydrazine products Lys-Gly and Lys-Ser are shown. The product control Lys-Gly and Lys-Ser were also generated by using the previously characterized GlyHS (Q8KGM6) and SerHS (A0A552E3D4) as controls.[6]

**Supplementary Fig. 37**

a

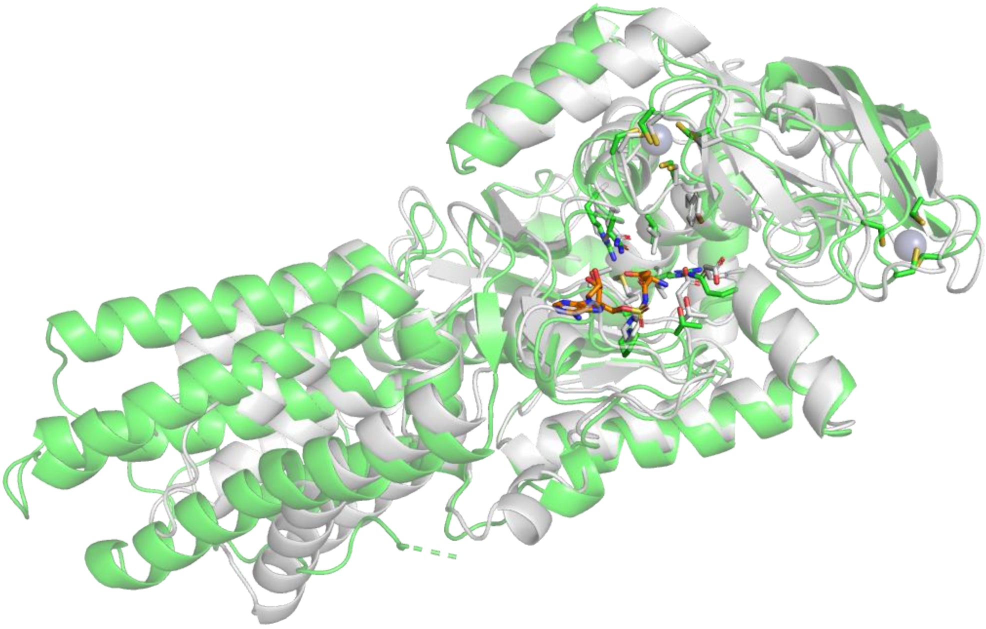

b

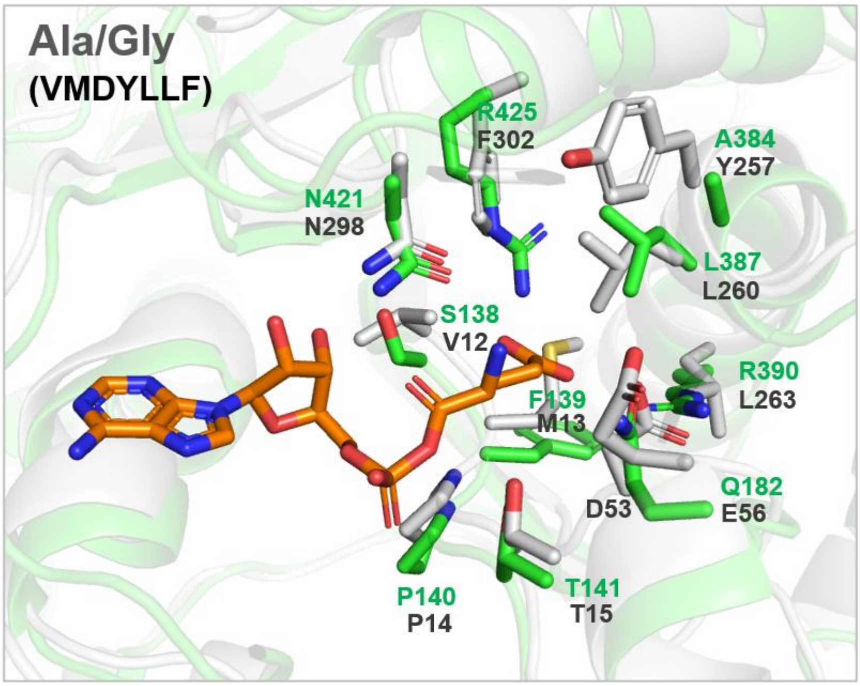

c

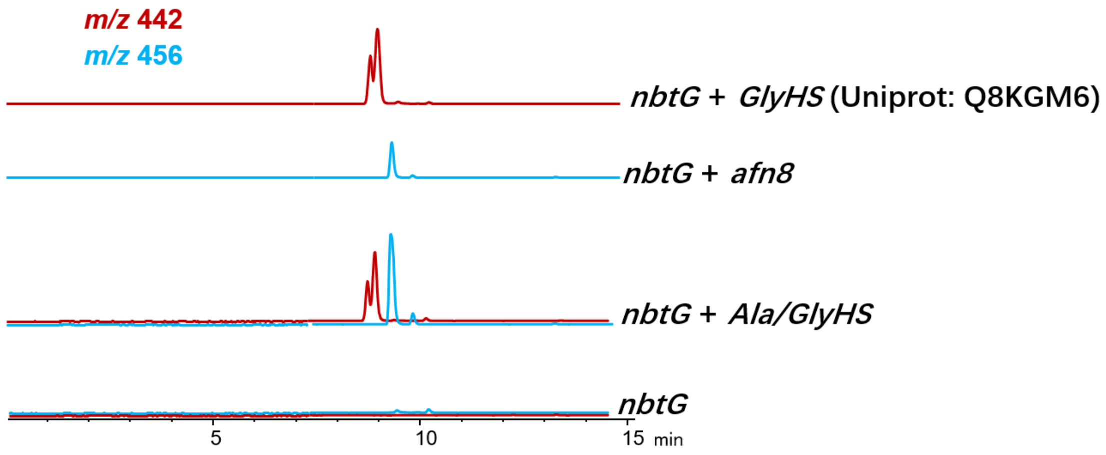

**Supplementary Fig. 37** Characterization of the newly-identified Ala/Gly-utilizing enzyme Ala/GlyHS (NCBI: WP_011001756). (a) Superposition of the AlphaFold-predicted structure of Ala/GlyHS (gray) onto the co-crystal structure of PyrN-C with DGA (green). (b) Comparison of the amino acid substrate binding site residues between PyrN and Ala/GlyHS. Note: Ala/GlyHS is a PyrN-C homolog from *Ralstonia pseudosolanacearum*. The specificity code for Ala/GlyHS is 'VMDYLLF'. (c) Determination of the amino acid specificity of Ala/GlyHS using an in vivo biotransformation assay. The EICs for the $[M+H]^+$ ions (*m*/*z* 442, 456) of Fmoc-derived hydrazine products Lys-Gly and Lys-Ala are shown. The product control Lys-Gly and Lys-Ala were also generated by using the previously characterized GlyHS (Q8KGM6)[6] and Afn8.[40]

**Supplementary Fig. 38**

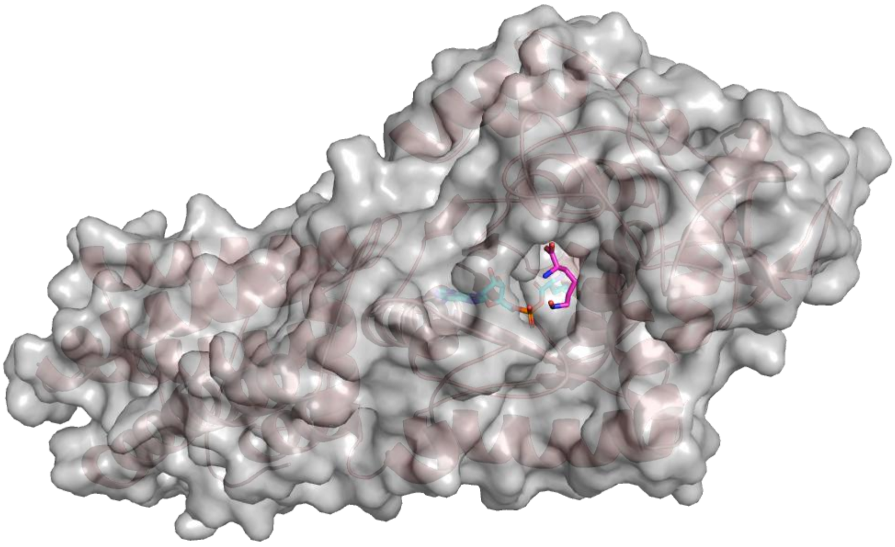

**Supplementary Fig. 38** The co-crystal structure of PyrN-C in complex with LGA and OH-Lys reveals a solvent-accessible, open substrate-binding cavity.

#### Supplementary Fig. 39

**(a)**

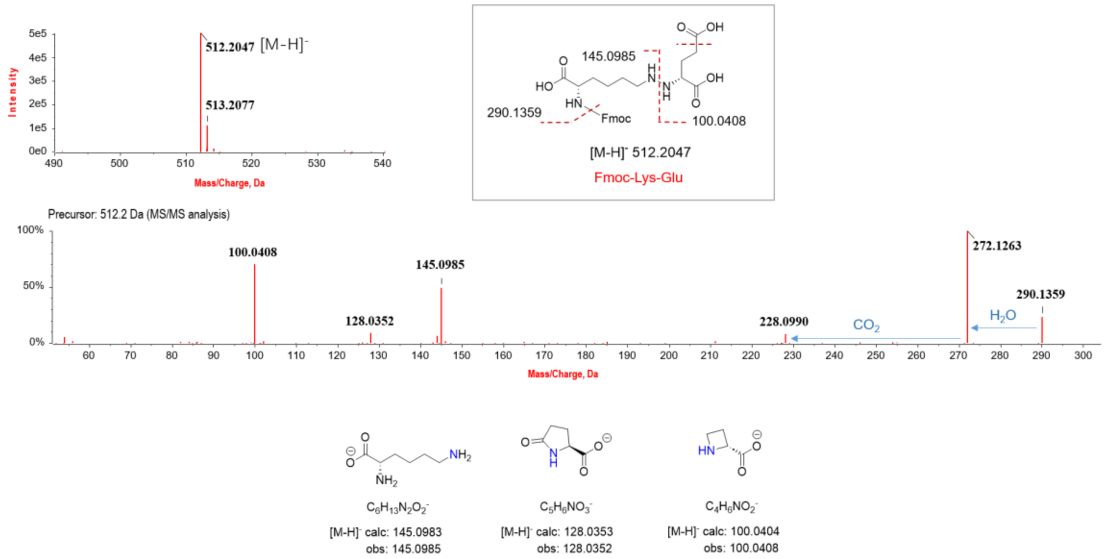

**(b)**

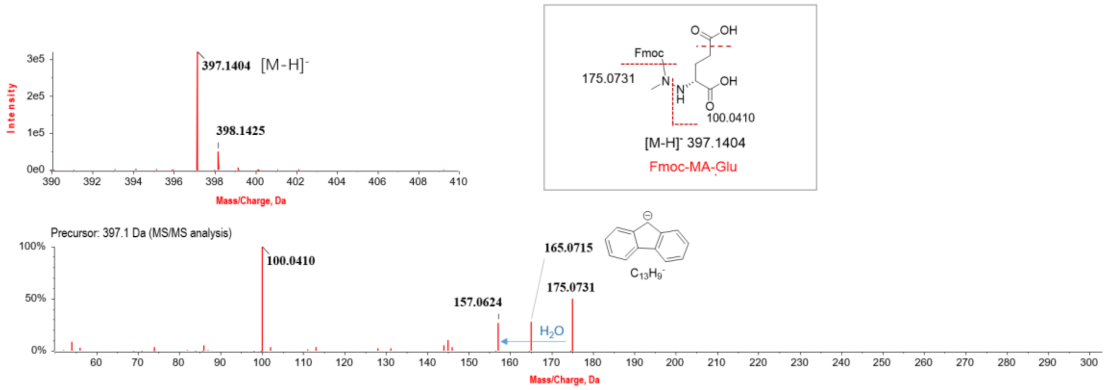

**(c)**

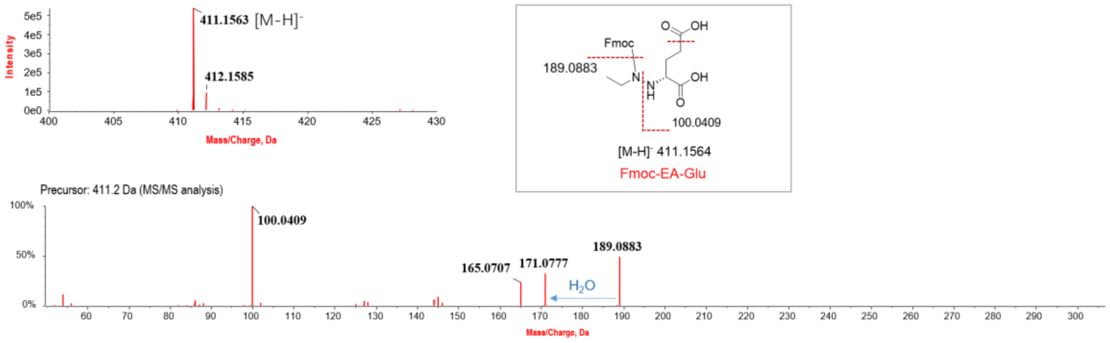

**(d)**

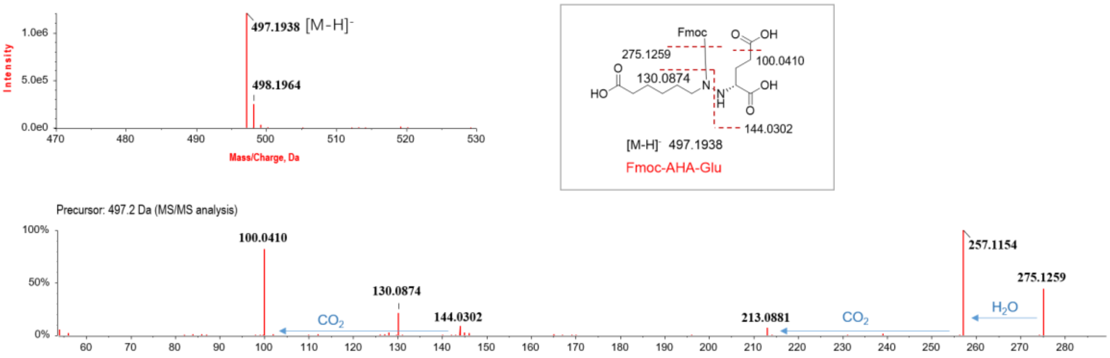

**(e)**

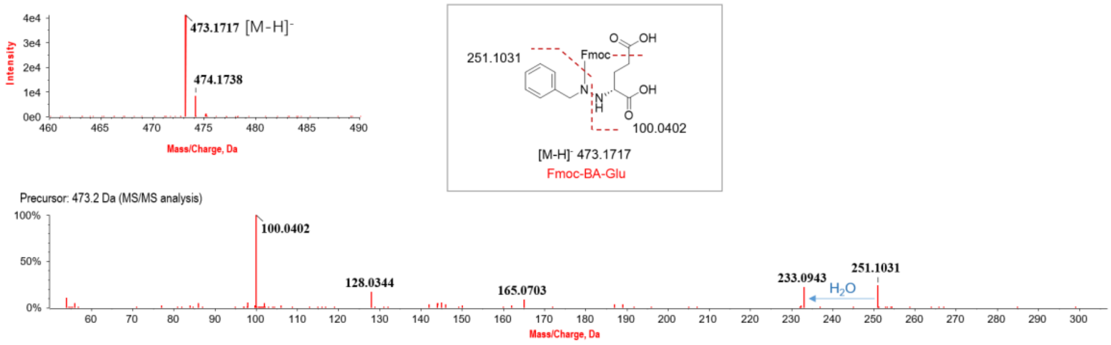

**(f)**

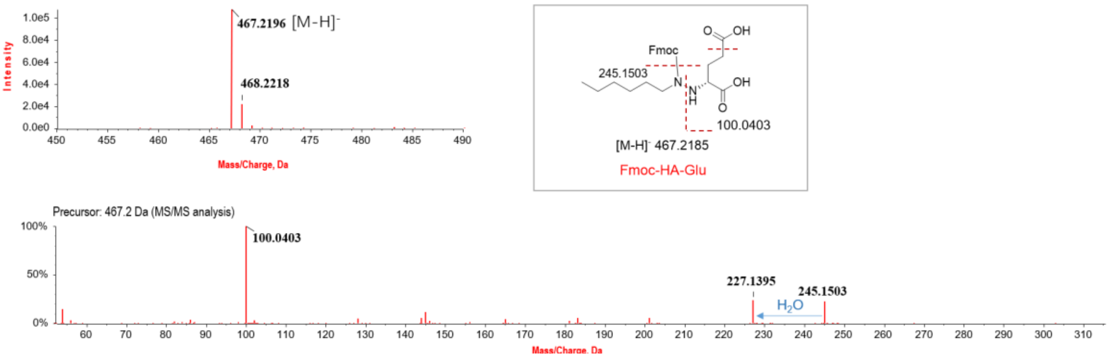

**Supplementary Fig. 39** LC-HR-MS/MS analysis (under negative mode) of the hydrazine products from the in vitro reaction mixtures of PyrN with D-Glu and various *N*-substituted hydroxylamines: (a) Native substrate $N^6$-hydroxy-L-lysine, (b) *N*-methylhydroxylamine (MHA), (c) *N*-ethylhydroxylamine (EHA), (d) *N*-aminohexylhydroxylamine (AHHA), (e) *N*-benzylhydroxylamine (BHA), (f) *N*-hexylhydroxylamine (HHA). Selected diagnostic fragment signals are indicated in the chemical structures of the corresponding products, and the fragmentation pattern is consistent with a hydrazine product, as revealed in our previous study[6].

**Supplementary Fig. 40**

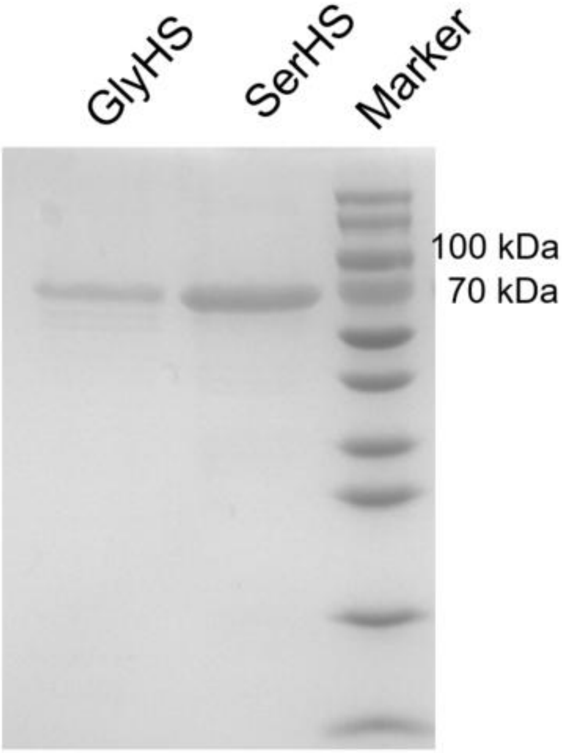

**Supplementary Fig. 40** The SDS-PAGE gel image of isolated GlyHS and SerHS. Note: GlyHS and SerHS are Gly- and Ser-utilizing didomain hydrazine synthetases from *Rhizobium loti* (Uniprot: Q8KGM6/NCBI: CAD31310) and *Microcystis aeruginosa* (Uniprot: A0A552E3D4/NCBI:WP_004159161), respectively.[6]

**Supplementary Fig. 41**

a

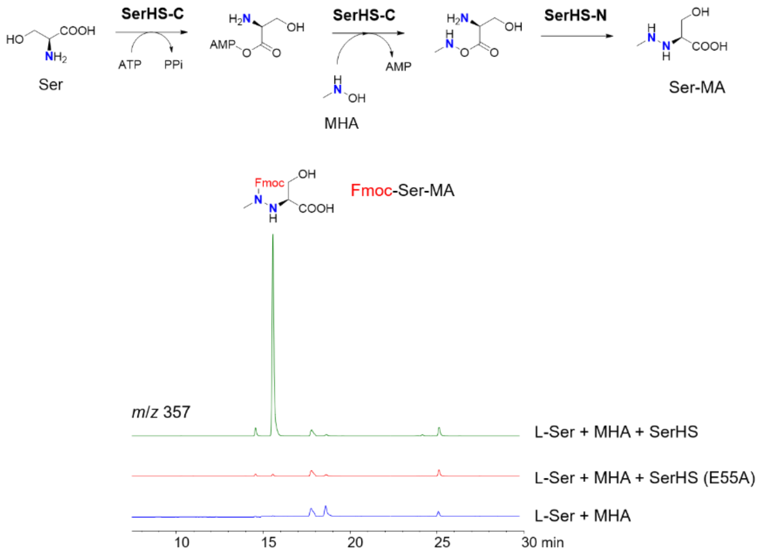

b

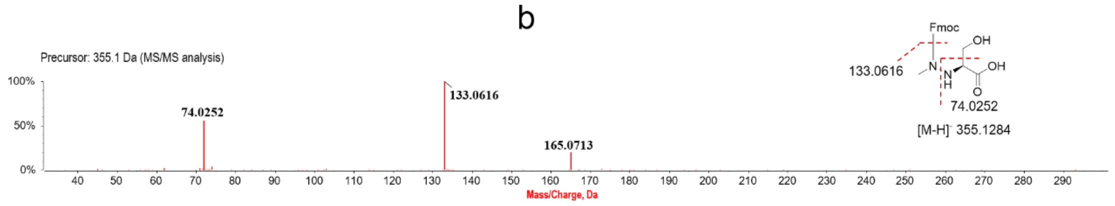

**Supplementary Fig. 41** In vitro assay of SerHS with L-Ser and MHA (a) and the LC-HR-MS/MS analysis (under negative mode) of the product from in vitro reaction mixture (b). Note: The SerHS variant E55A, which carries an inactive N-terminal cupin domain (SerHS-N), is used as a negative control.

#### Supplementary Fig. 42

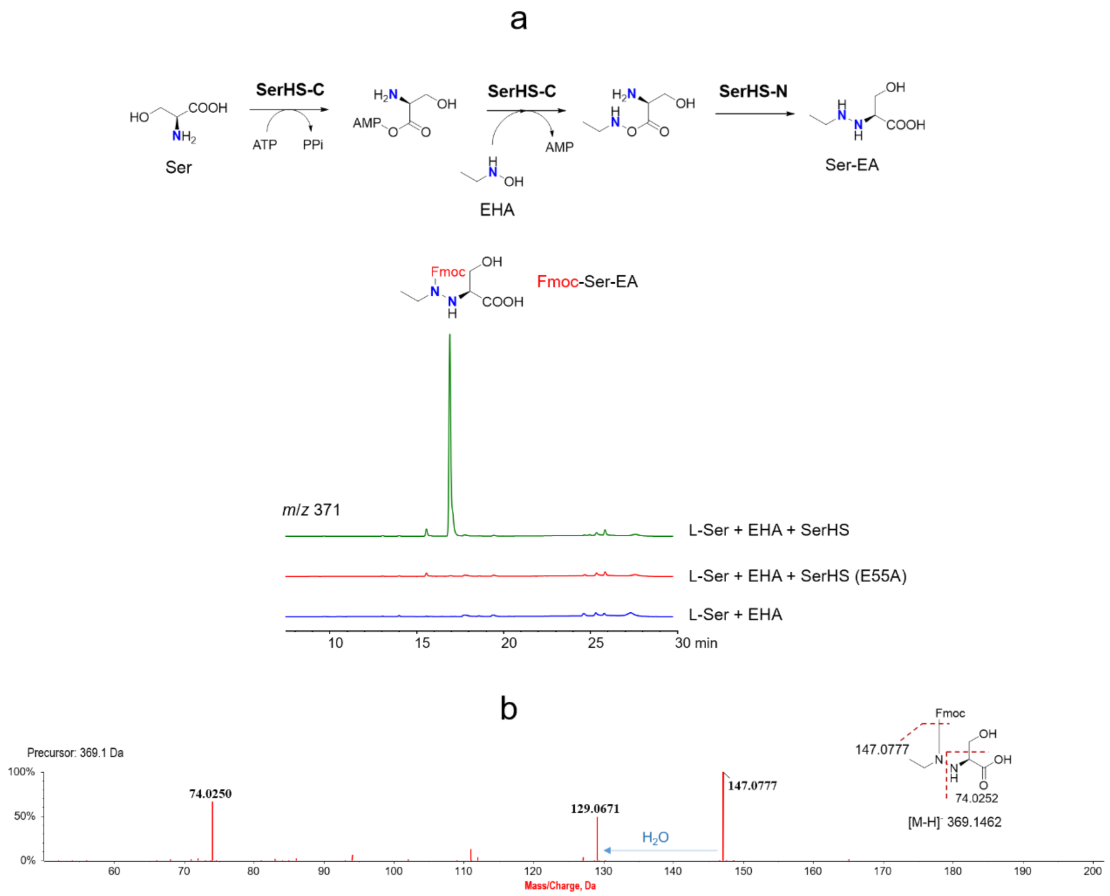

**Supplementary Fig. 42** In vitro assay of SerHS with L-Ser and EHA (a) and the LC-HR-MS/MS analysis (under negative mode) of the product from in vitro reaction mixture (b).

#### Supplementary Fig. 43

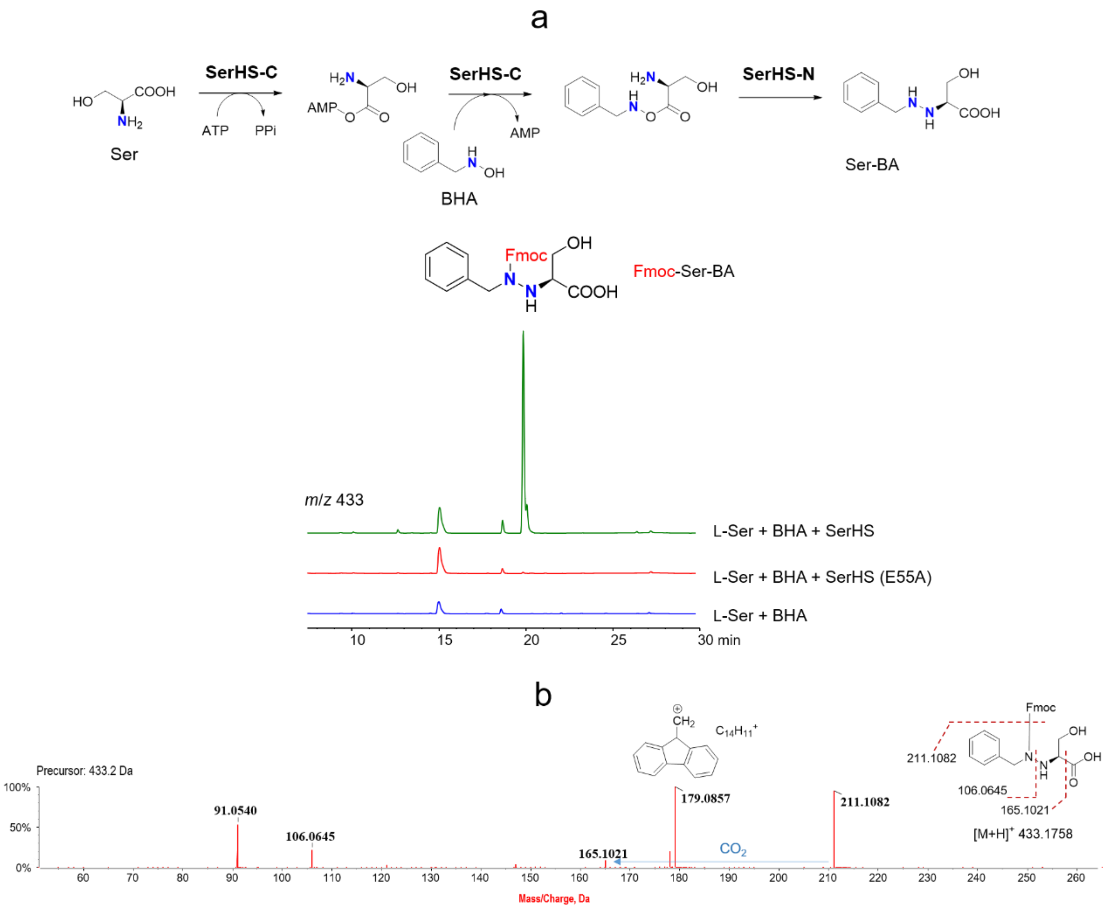

**Supplementary Fig. 44** In vitro assay of SerHS with L-Ser and BHA (a) and the LC-HR-MS/MS analysis (under positive mode) of the product from in vitro reaction mixture (b).

**Supplementary Fig. 45**

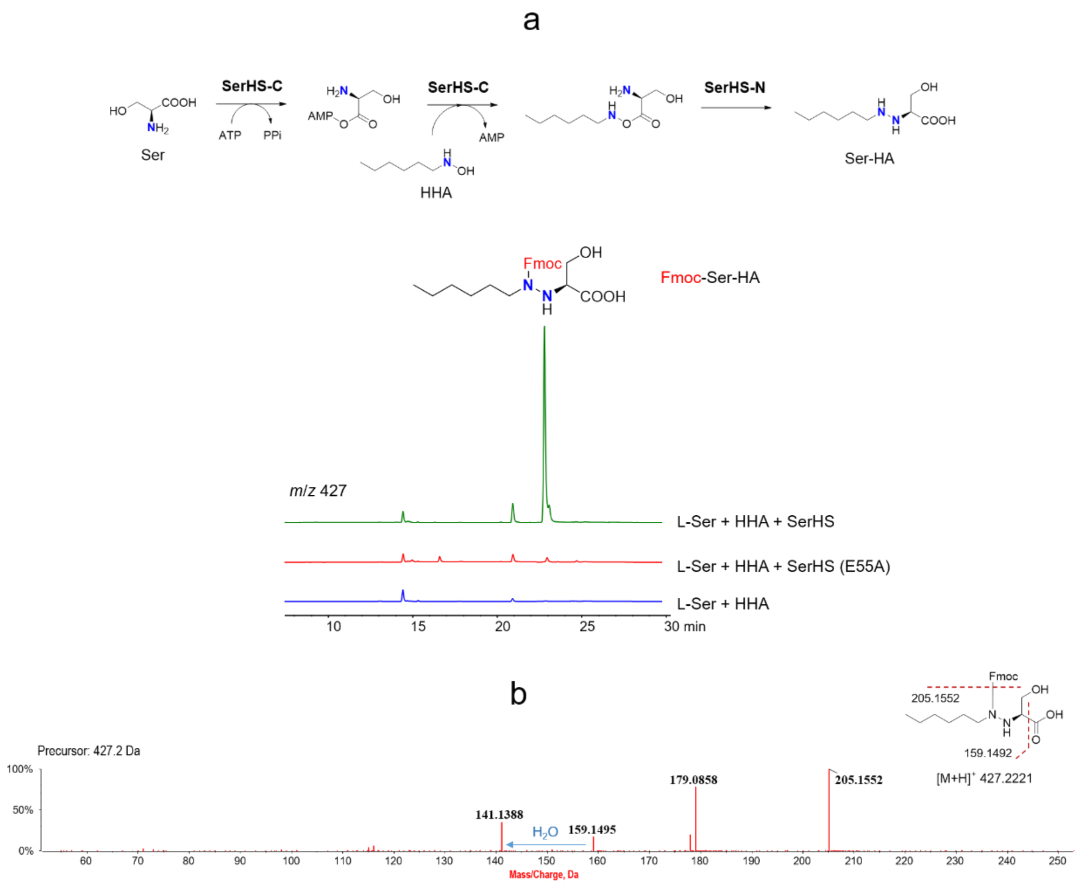

**Supplementary Fig. 45** In vitro assay of SerHS with L-Ser and HHA (a) and the LC-HR-MS/MS analysis (under positive mode) of the product from in vitro reaction mixture (b).

**Supplementary Fig. 46**

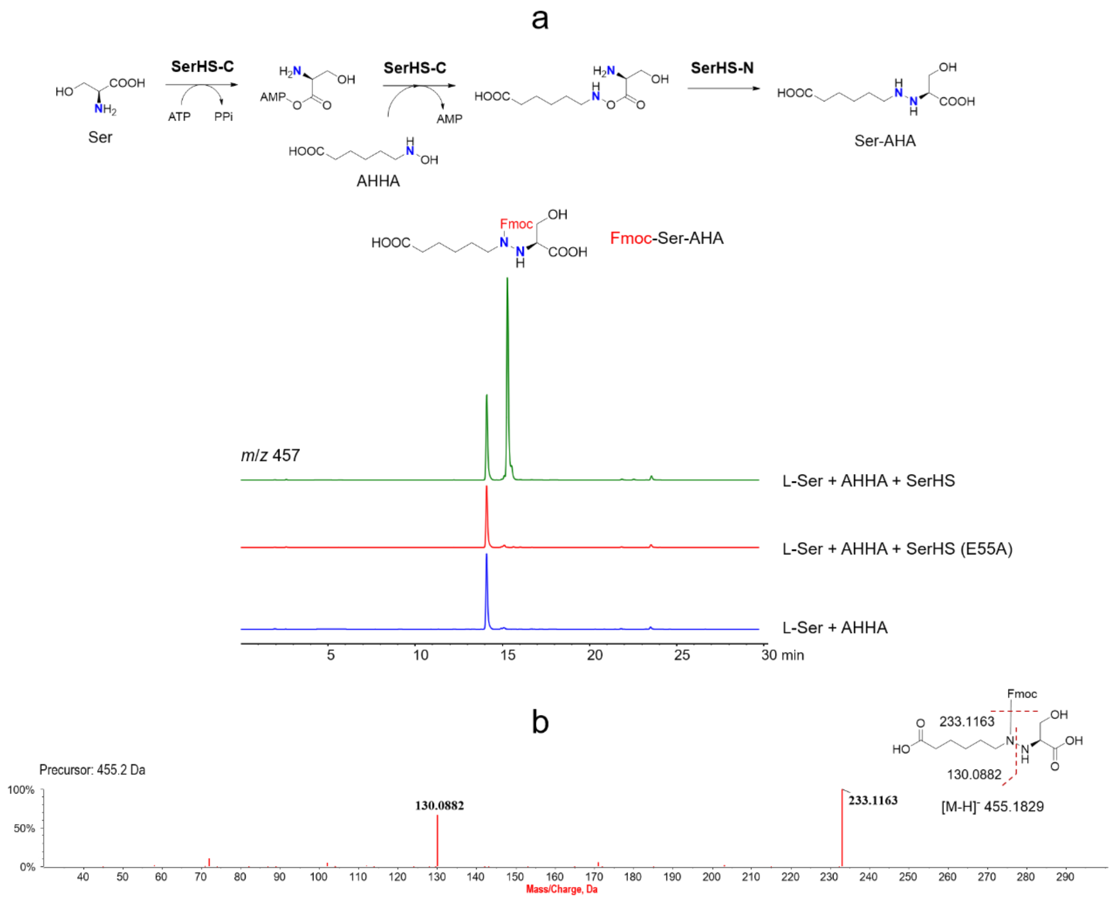

**Supplementary Fig. 46** In vitro assay of SerHS with L-Ser and AHHA (a) and the LC-HR-MS/MS analysis (under negative mode) of the product from in vitro reaction mixture (b).

**Supplementary Fig. 47**

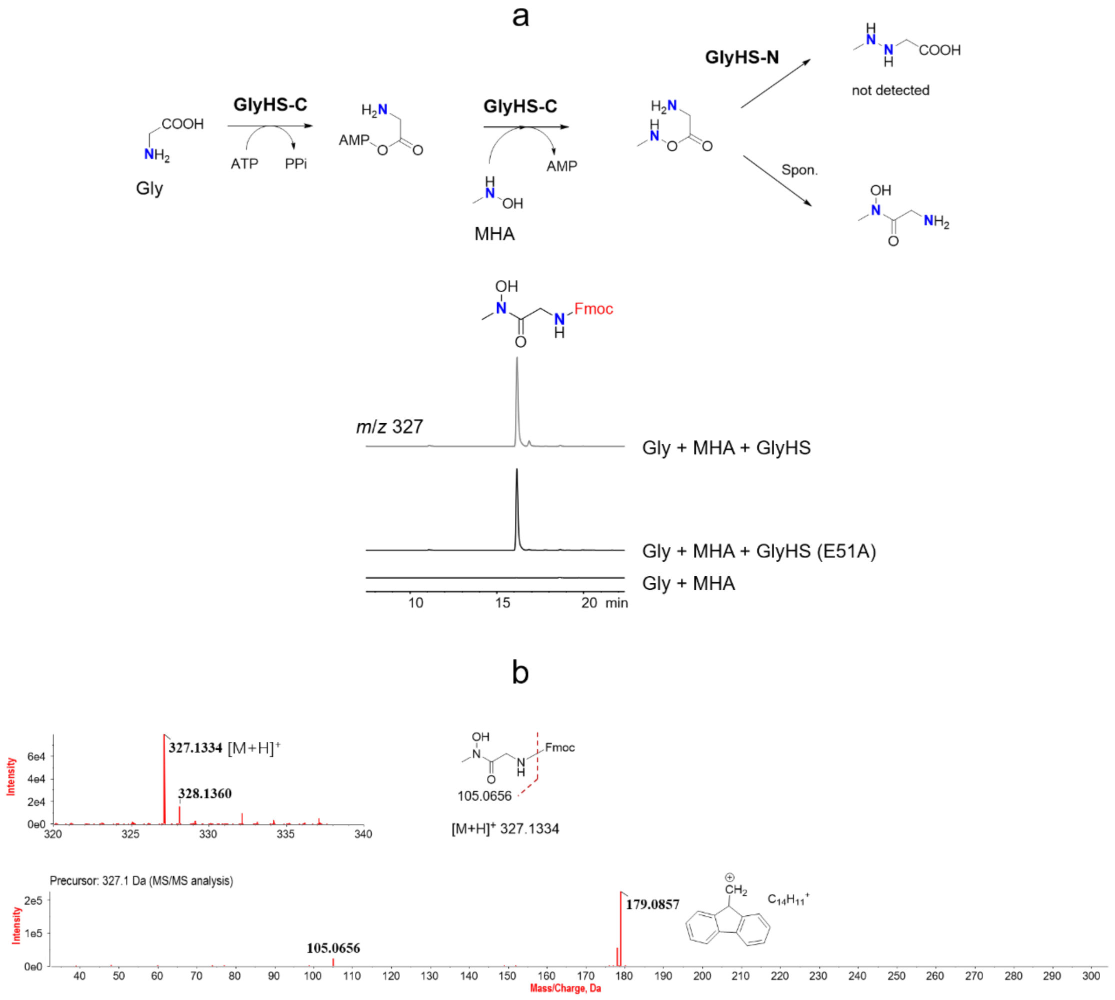

**Supplementary Fig. 47** In vitro assay of GlyHS with glyine and MHA (a) and the LC-HR-MS/MS analysis (under positive mode) of the product from in vitro reaction mixture (b). Note: Only amide shunt product was detected, indicating that the GlyHS-N cupin domain cannot process the non-native *O*-aminoacyl-hydroxylamine ester intermediate. The GlyHS variant E51A, which carries an inactive N-terminal cupin domain (GlyHS-N), was used as a negative control.

**Supplementary Fig. 48**

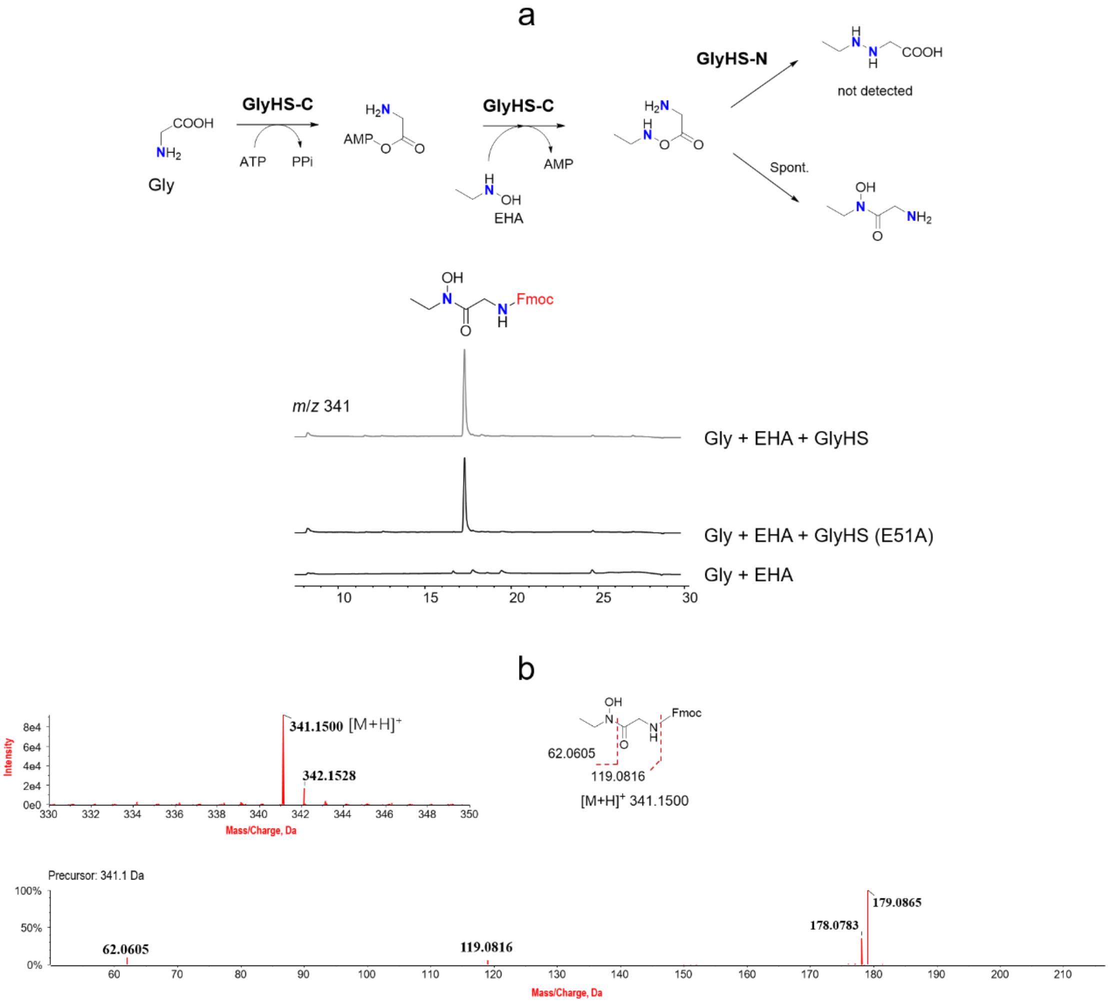

**Supplementary Fig. 48** In vitro assay of GlyHS with glyine and EHA (a) and the LC-HR-MS/MS analysis (under positive mode) of the product from in vitro reaction mixture (b). Note: signal *m*/*z* 179 is a fragment derived from the Fmoc moiety. Only amide shunt product was detected, indicating that the GlyHS-N cupin domain cannot process the non-native *O*-aminoacyl-hydroxylamine ester intermediate.

#### Supplementary Fig. 49

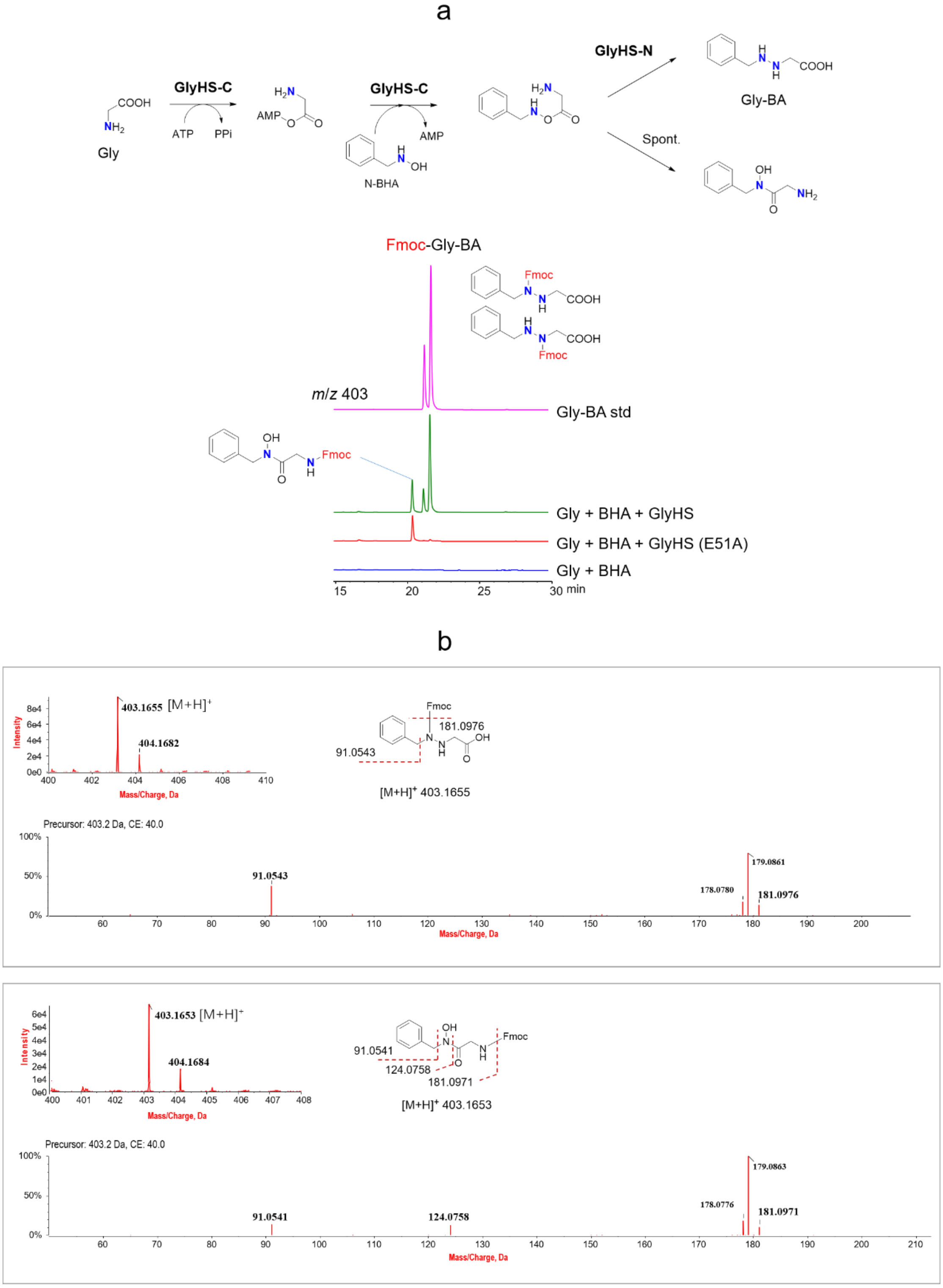

**Supplementary Fig. 49** In vitro assay of GlyHS with glyine and BHA (a) and the LC-HR-MS/MS analysis (under positive mode) of the hydrazine product and the amide shunt product from in vitro reaction mixture (b). Note: chemically prepared Gly-BA standard was used a product control in (a). Both hydrazine and amide shunt products were detected in this reaction mixture. The synthetic route for Gly-BA standard was provided in Supplementary Fig. 54.

**Supplementary Fig. 50**

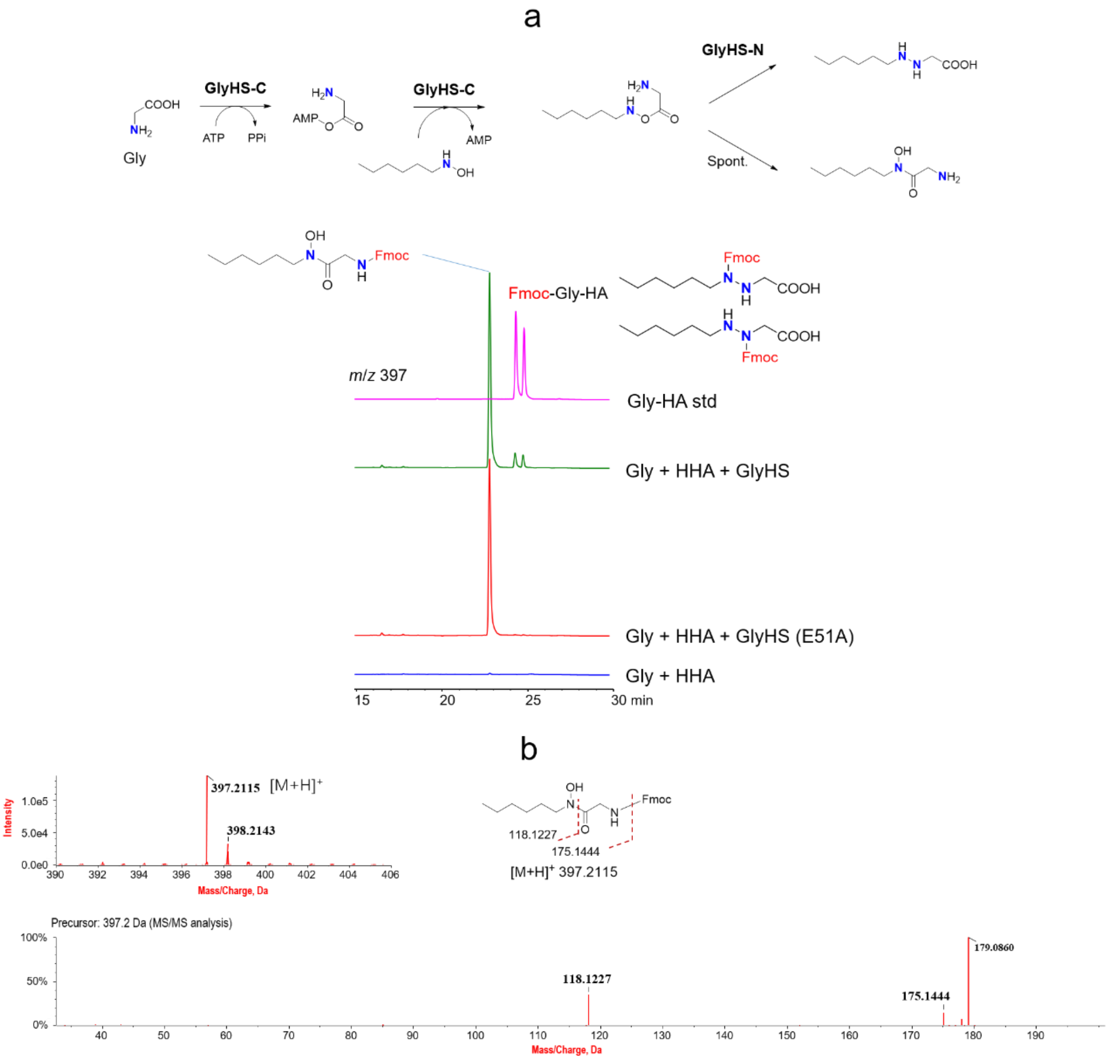

**Supplementary Fig. 50** In vitro assay of GlyHS with glyine and HHA (a) and the LC-HR-MS/MS analysis (under positive mode) of the amide shunt product from in vitro reaction mixture (b). Note: chemically prepared Gly-HA standard was used a product control in (a), and the major product was the amide, only a small amount of Gly-HA was detected. The synthetic route for Gly-HA standard was provided in Supplementary Fig. 55.

#### Supplementary Fig. 51

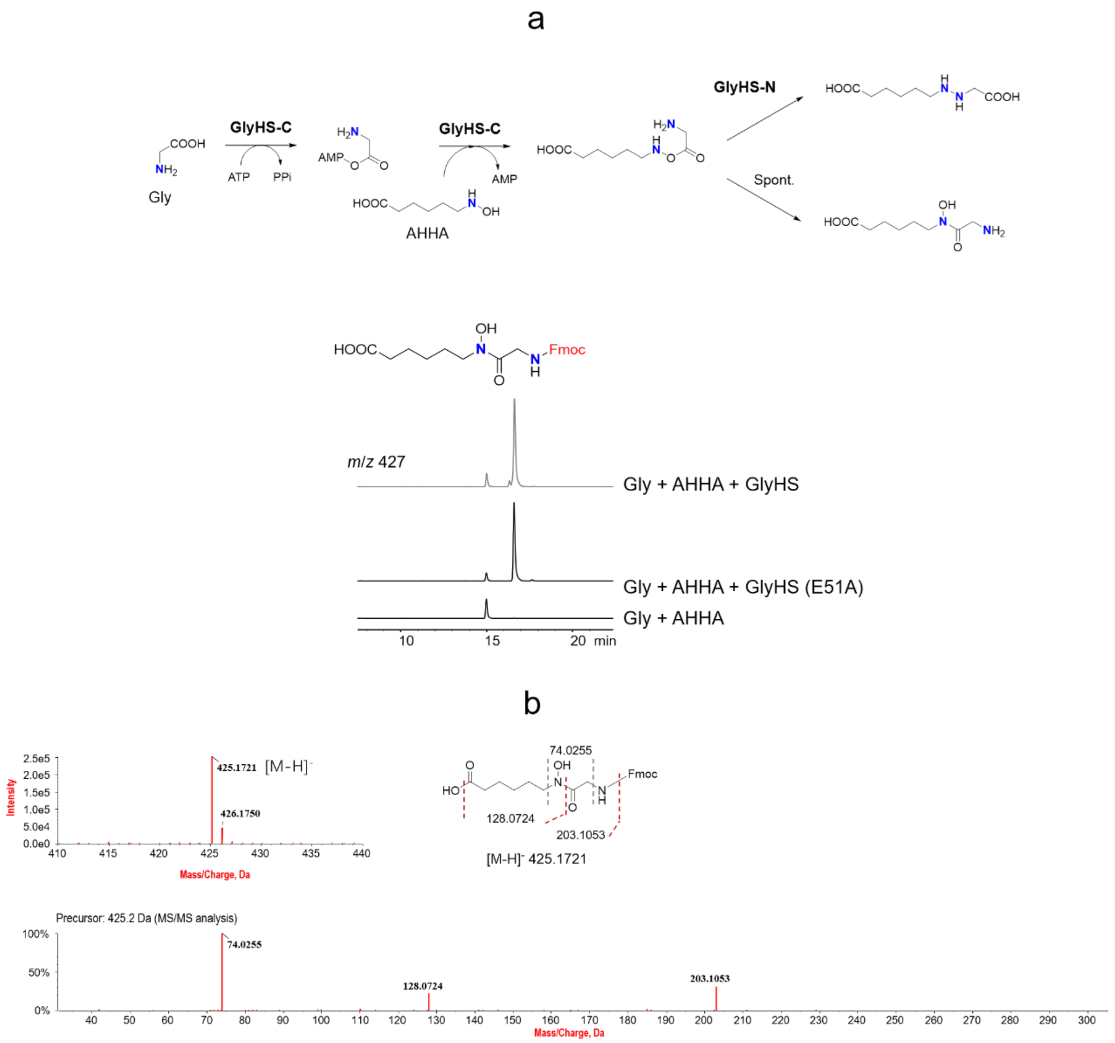

**Supplementary Fig. 51** In vitro assay of GlyHS with glyine and AHHA (a) and the LC-HR-MS/MS analysis (under negative mode) of the product from in vitro reaction mixture (b). Only amide shunt product was detected, indicating that the GlyHS-N cupin domain cannot process the corresponding non-native *O*-aminoacyl-hydroxylamine ester intermediate.

**Supplementary Fig. 52**

**a**

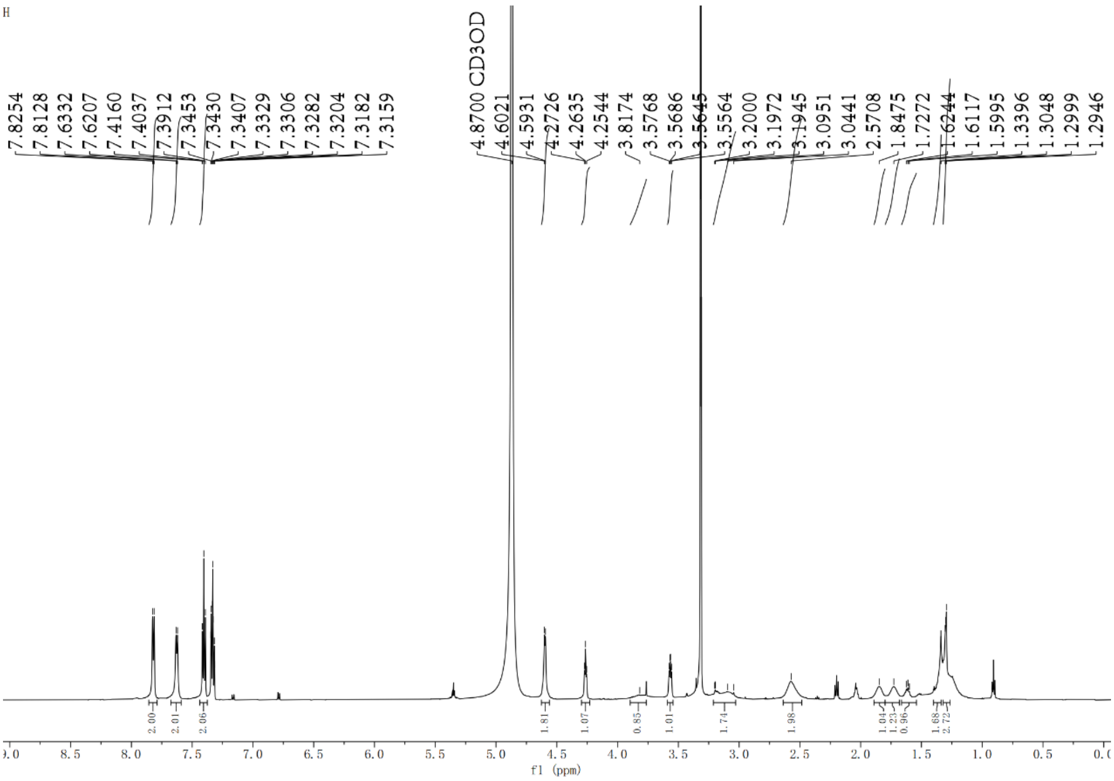

**b**

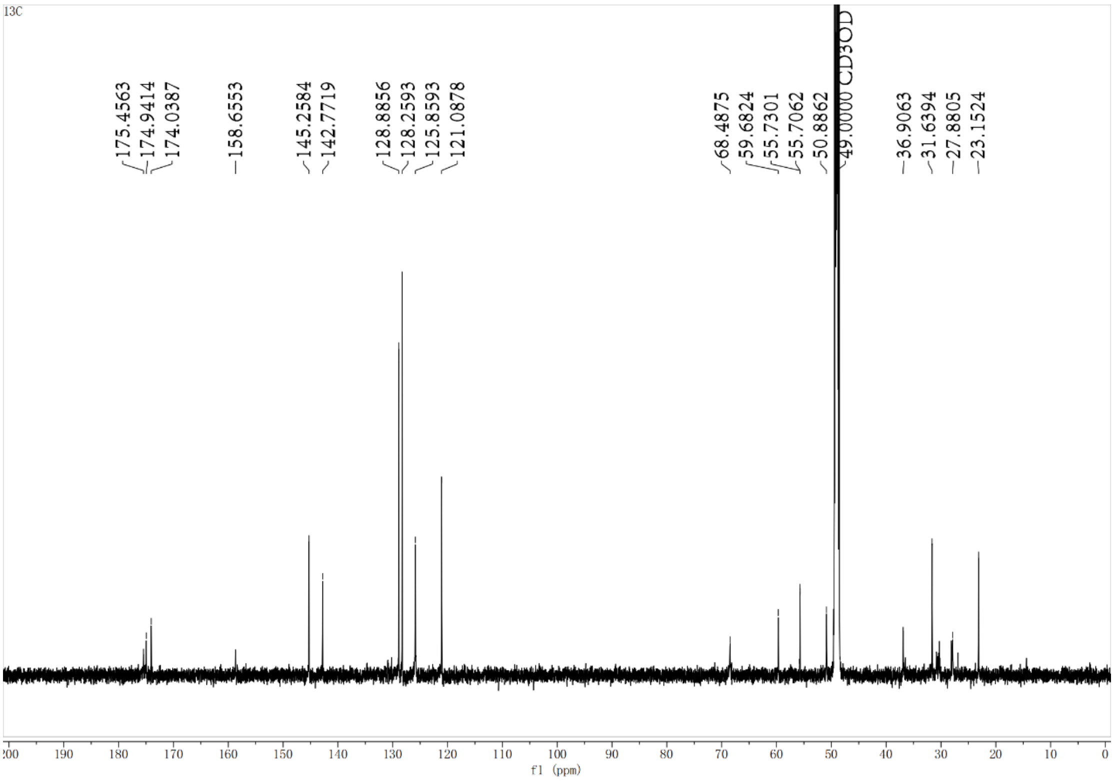

c

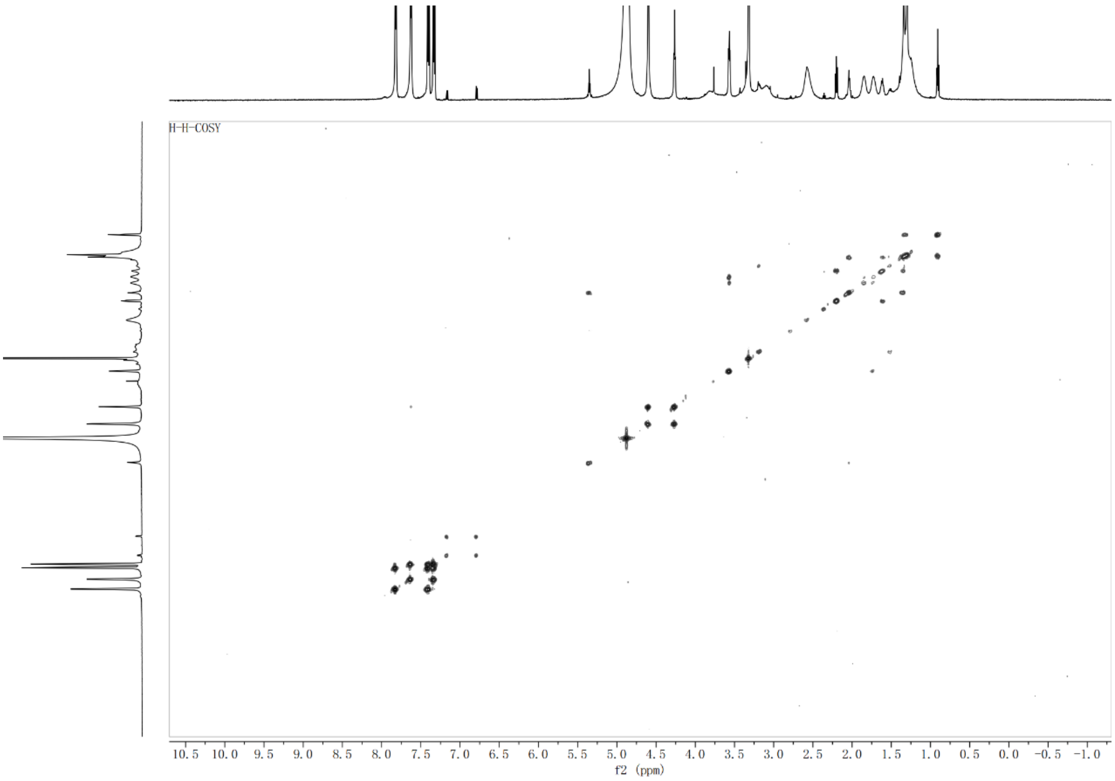
H-H-COSY
10.5 10.0 9.5 9.0 8.5 8.0 7.5 7.0 6.5 6.0 5.5 5.0 4.5 4.0 3.5 3.0 2.5 2.0 1.5 1.0 0.5 0.0 -0.5 -1.0
f2 (ppm)

d

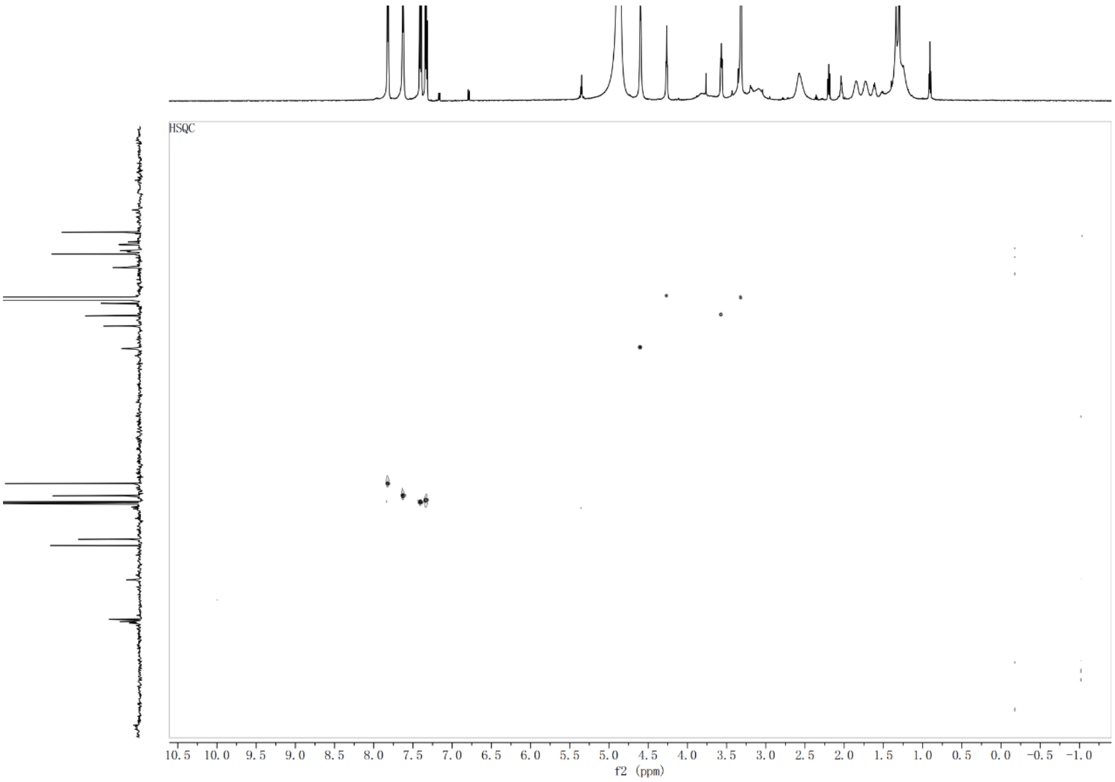
HSQC
10.5 10.0 9.5 9.0 8.5 8.0 7.5 7.0 6.5 6.0 5.5 5.0 4.5 4.0 3.5 3.0 2.5 2.0 1.5 1.0 0.5 0.0 -0.5 -1.0
f2 (ppm)

**e**

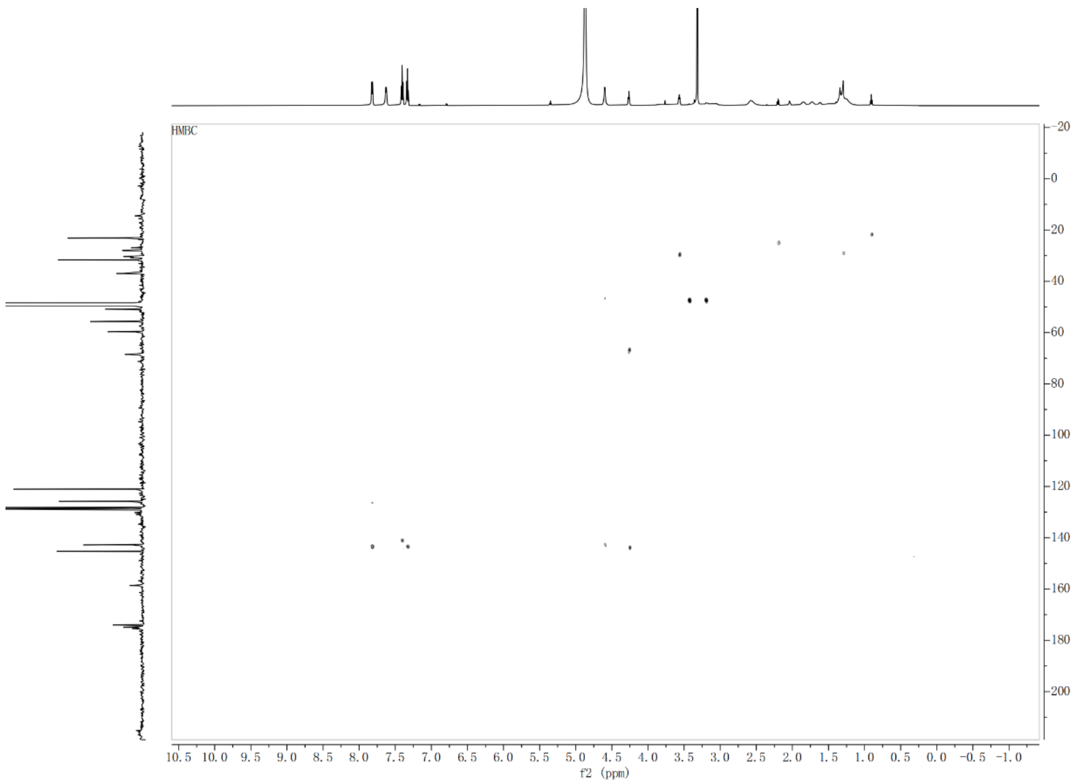

**Supplementary Fig. 52** NMR spectra of Lys-Asp-Fmoc. (a) $^{1}H$ NMR spectrum of Lys-Asp-Fmoc. (b) $^{13}C$ NMR spectrum of Lys-Asp-Fmoc. (c) $^{1}H$-$^{1}H$-COSY NMR spectrum of Lys-Asp-Fmoc. (d) HSQC NMRspectrum of Lys-Asp-Fmoc. (e) $^{1}H$-$^{13}C$-HMBC NMR spectrum of Lys-Asp-Fmoc. The isolation of Lys-Asp-Fmoc and Lys-Asn-Fmoc were performed similarly as described for Lys-Glu-Fmoc[6].

**Supplementary Fig. 53**

**a**

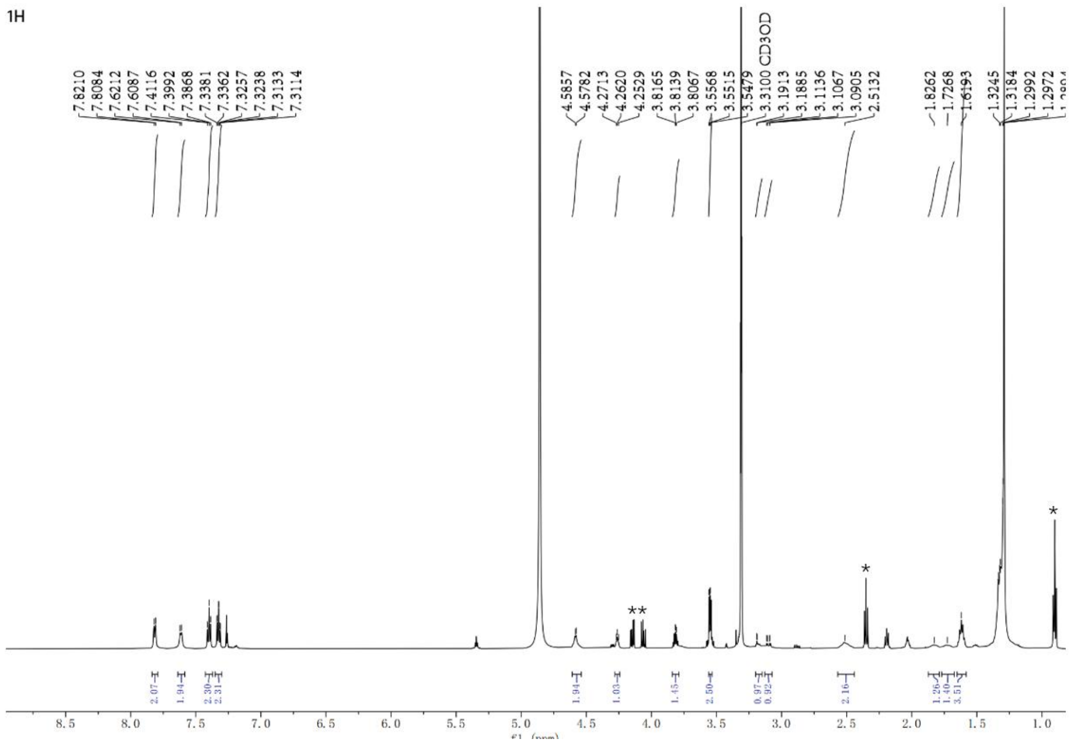

**b**

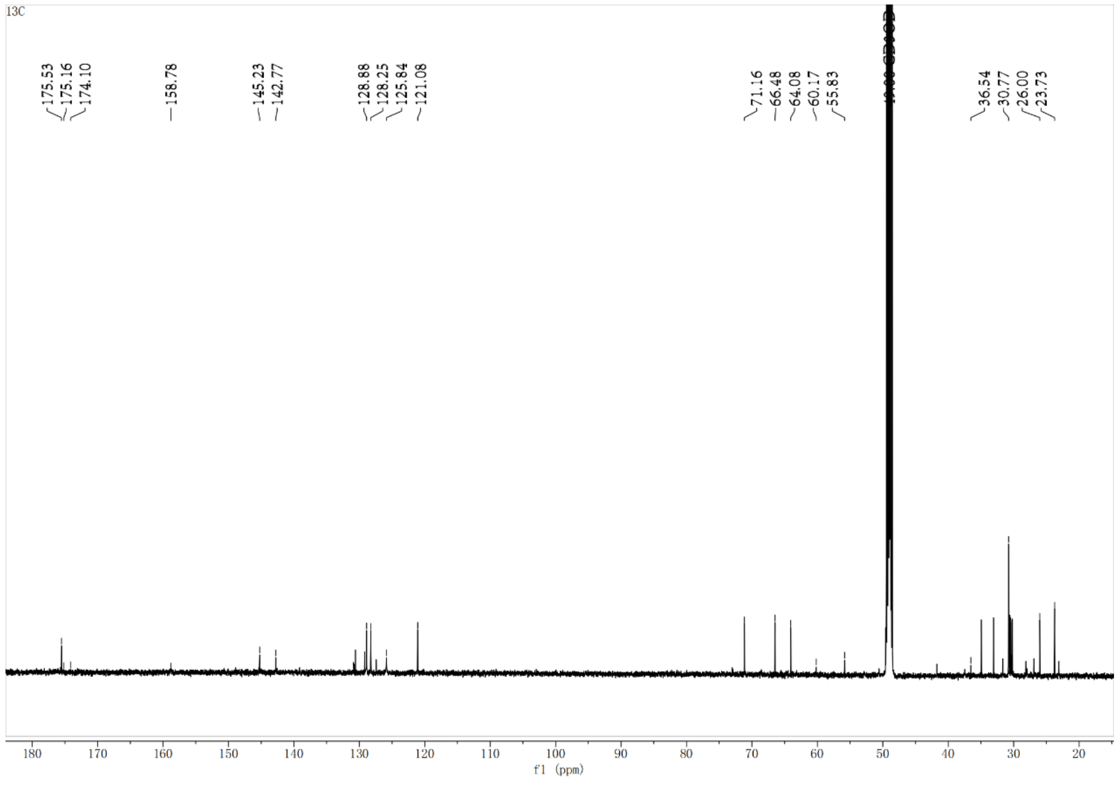

**c**

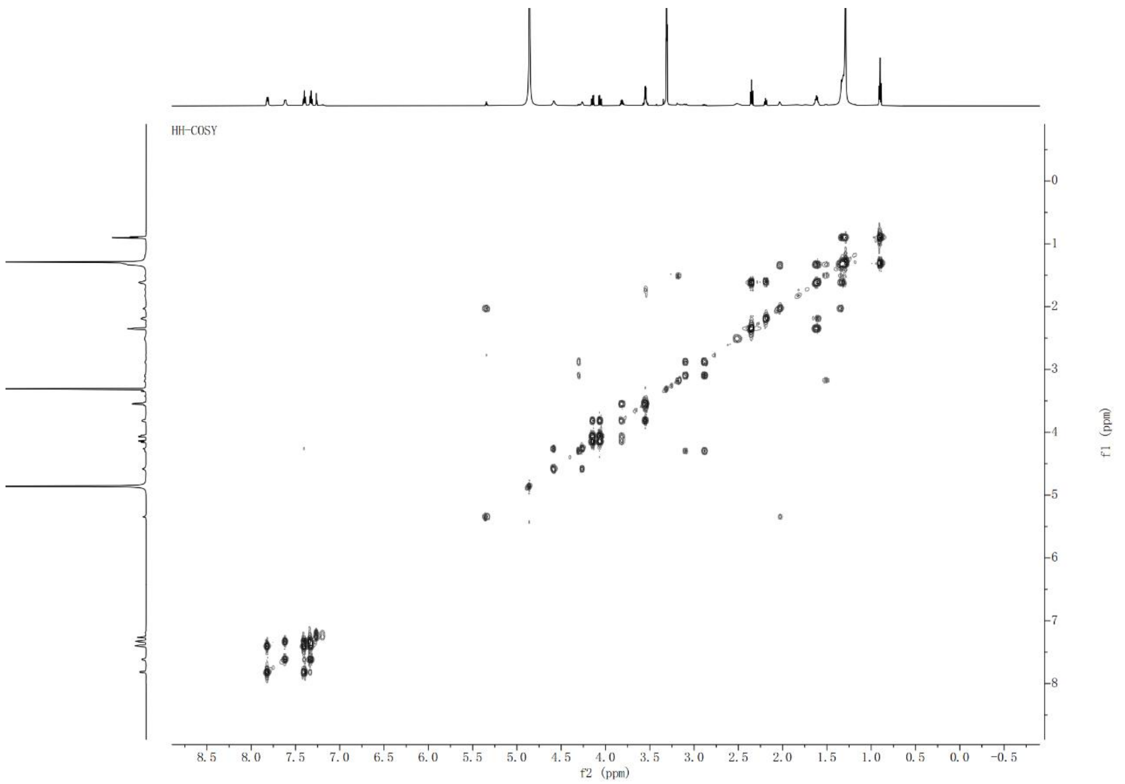

**d**

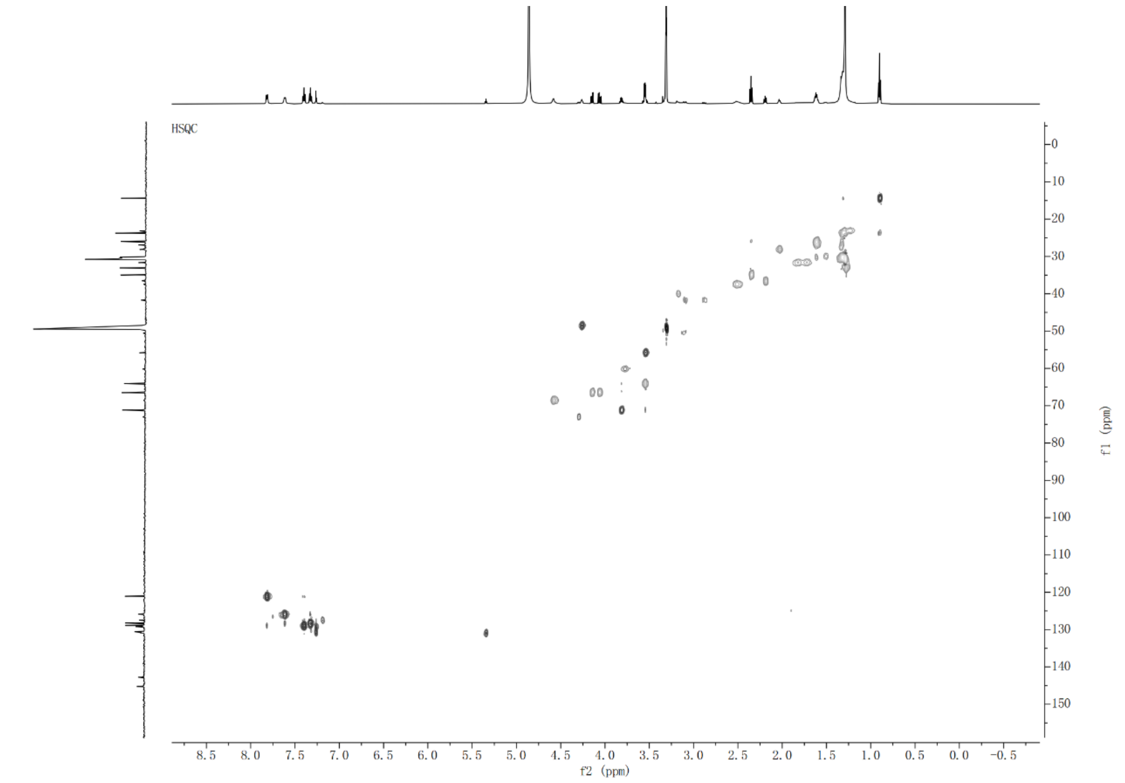

**e**

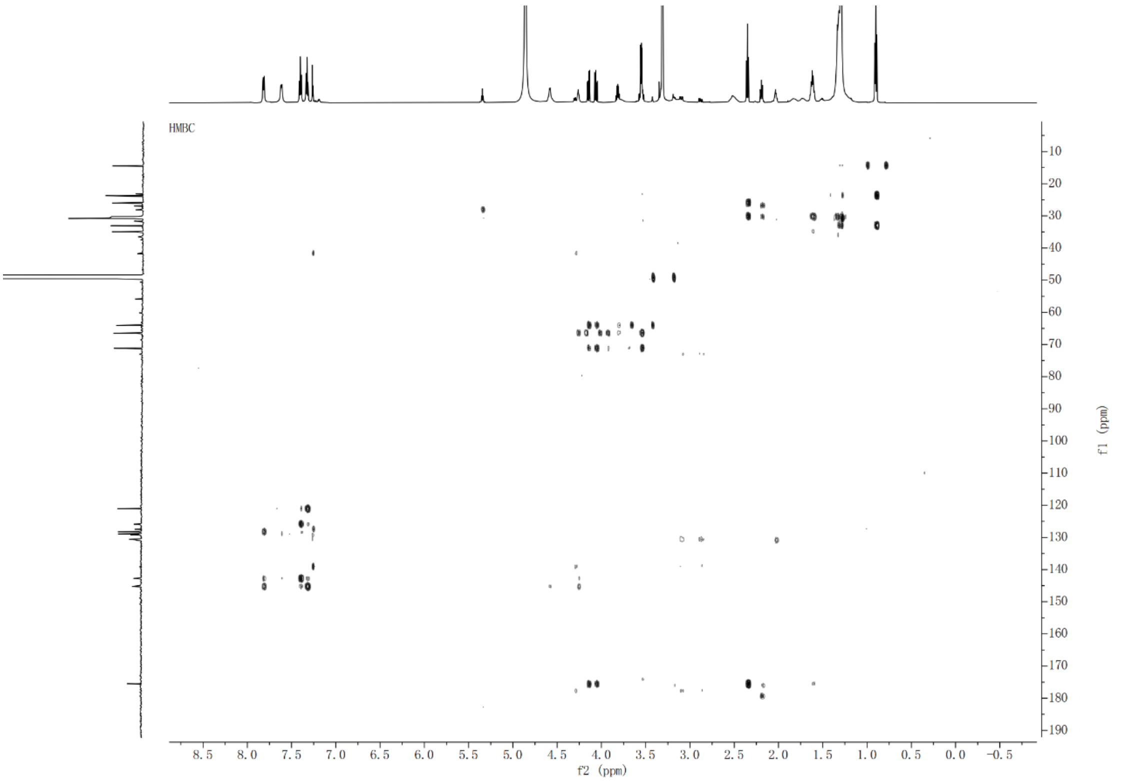

**Supplementary Fig. 53** NMR spectra of Lys-Asn-Fmoc. (a) $^{1}H$ NMR spectrum of Lys-Asn-Fmoc. Note: *These signals were attributed to impurities, as the compound is unstable, which prevented further purification. (b) $^{13}C$ NMR spectrum of Lys-Asn-Fmoc. (c) $^{1}H$-$^{1}H$-COSY NMR spectrum of Lys-Asn-Fmoc. (d) HSQC NMRspectrum of Lys-Asn-Fmoc. (e) $^{1}H$-$^{13}C$-HMBC NMR spectrum of Lys-Asn-Fmoc.

#### Supplementary Fig. 54

**a**

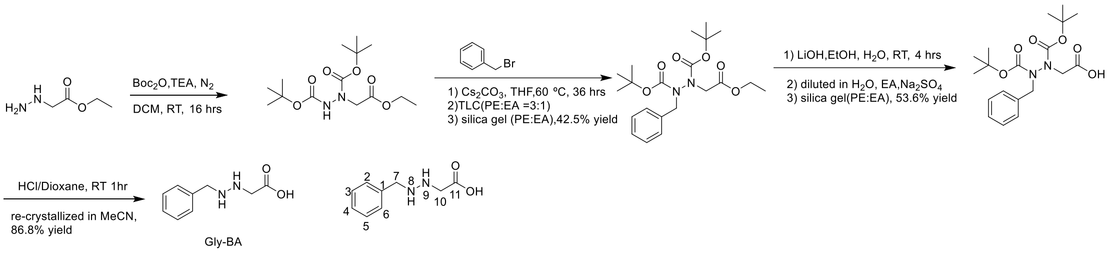

**b**

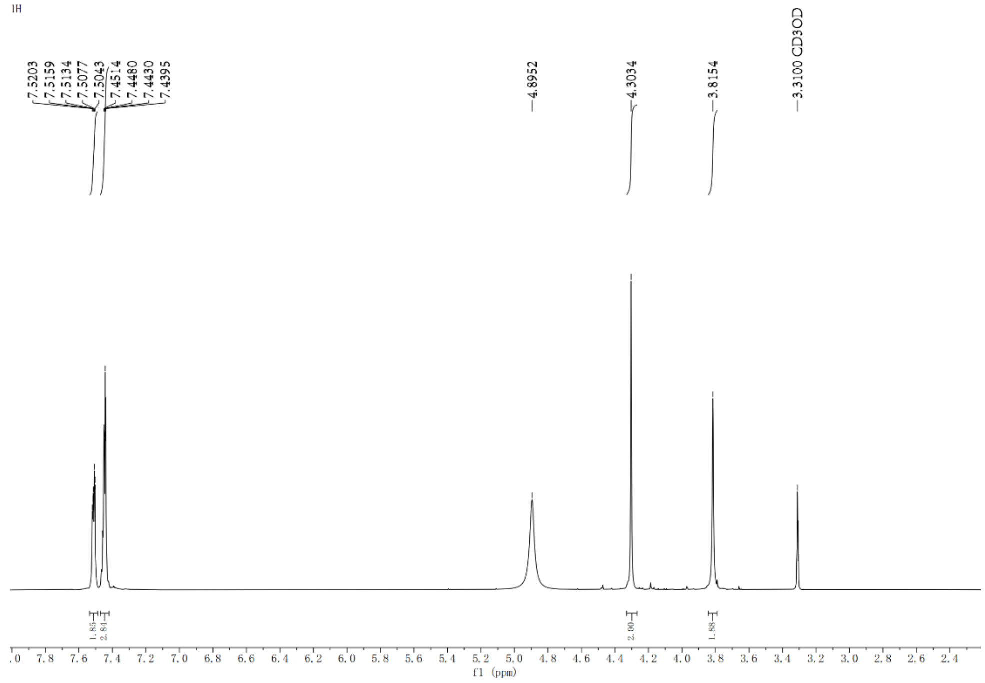

**c**

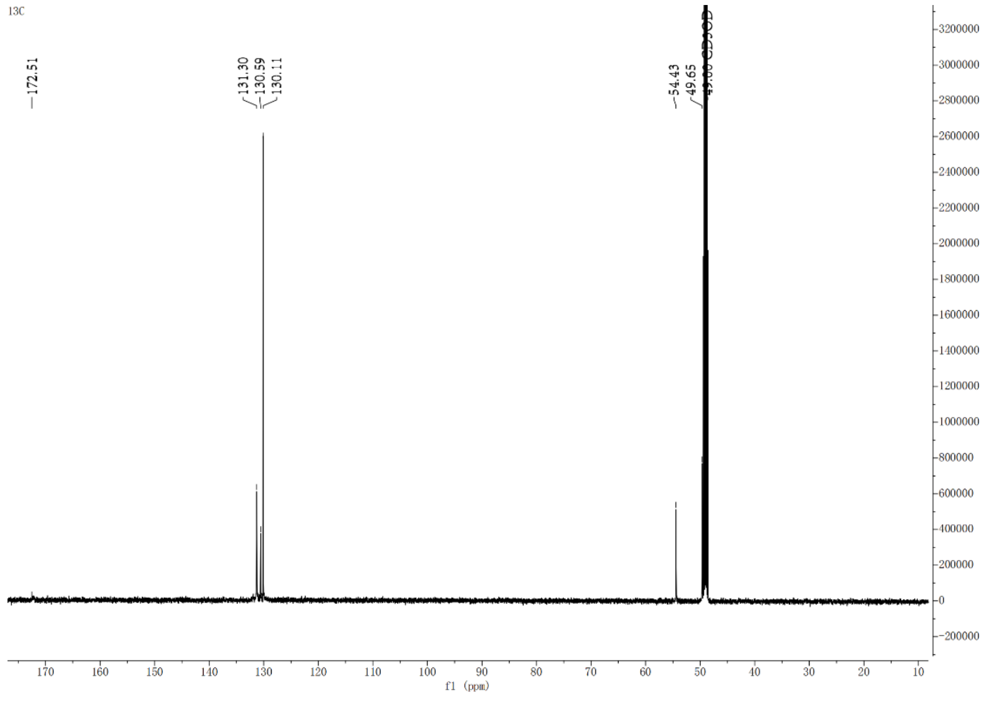

**Supplementary Fig. 54** Chemical synthesis of compound Gly-BA. (a) Chemical synthesis route of compound Gly-BA. (b) $^{1}$H NMR spectrum of compound Gly-BA ($CD_3OD$, 600 MHz). (c) $^{13}$C NMR spectrum of Gly-BA.

$^{1}$H NMR data for synthetic Gly-BA: $^{1}$H-NMR (600 MHz, DMSO-d$^{6}$) δ 7.52 (1H, m, -H-2); 7.52 (1H, m, H-6); 7.43 (1H, m, H-3); 7.43 (1H, m, H-4); 7.43 (1H, m, H-5); 4.43 (2H, m, H-7); 3.82 (2H, m, H-10). $^{13}$C-NMR (150 MHz, $CD_3OD$) δ 172.51 (C-11), 131.30 (C-1), 130.59 (C-3), 130.59 (C-5), 130.11 (C-2), 130.11 (C-4), 130.11 (C-6), 54.43 (C-10), 49.65 (C-7).

#### Supplementary Fig. 55

**a**

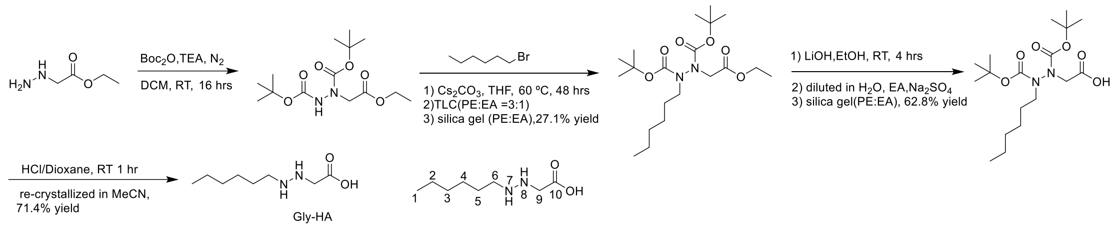

b

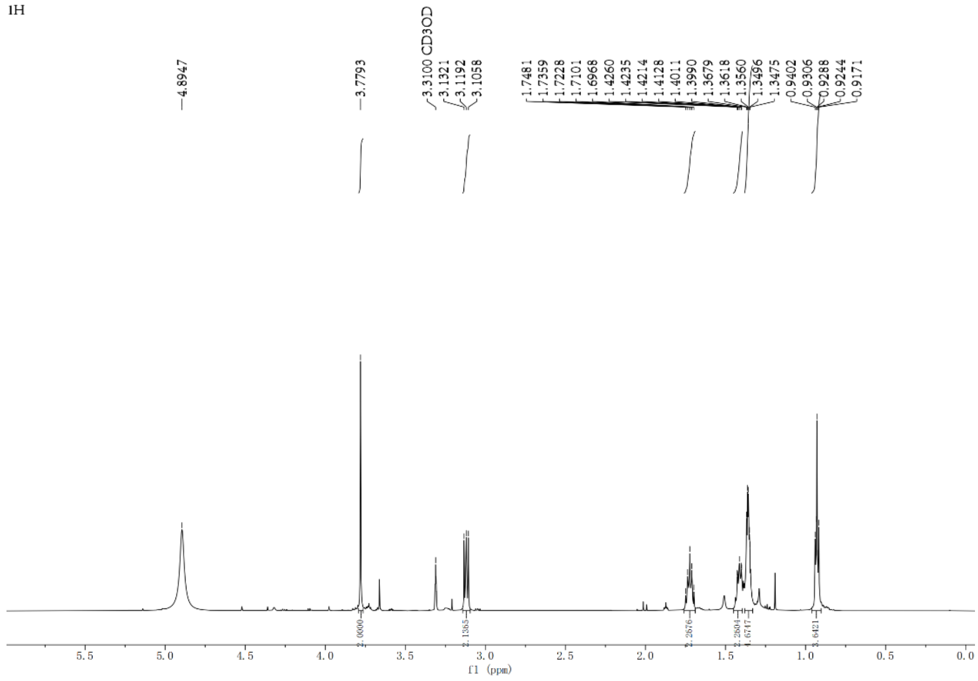

**c**

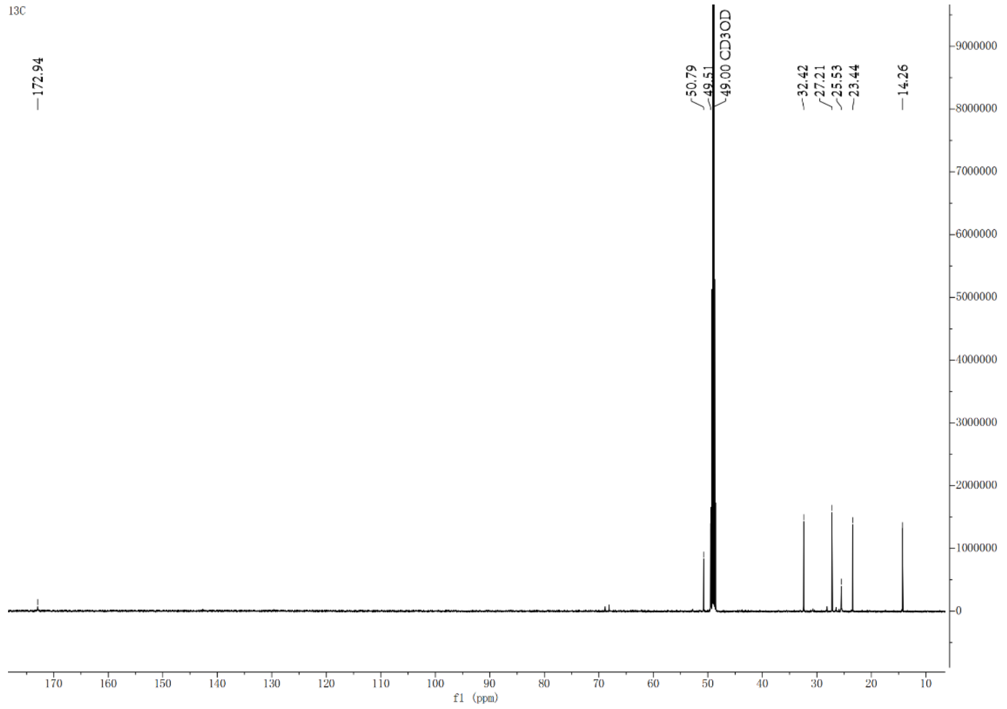

**Supplementary Fig. 55** Chemical synthesis of compound Gly-HA. (a) Synthetic route for Gly-HA. (b) $^{1}$H NMR spectrum compound Gly-HA ($CD_3OD$, 600 MHz). (c) $^{13}$C NMR spectrum of Gly- HA.

$^{1}$H NMR data for synthetic Gly-HA: $^{1}$H-NMR (600 MHz, $CD_3OD$) δ 3.78 (2H, m, H-9); 3.12 (2H, m, H-6); 1.72 (2H, m, H-5); 1.42 (2H, m, H-4); 1.36 (2H, m, H-3); 1.36 (2H, m, H-2); 0.92 (3H, m, H-1). $^{13}$C-NMR (150 MHz, $CD_3OD$) δ 172.94 (C-10), 50.79 (C-9), 49.51 (C-6), 27.21 (C-5), 25.53 (C-4), 32.42 (C-3), 23.44 (C-5), 14.26 (C-1).

#### RERFERENCE

(1) Kabsch, W. XDS. *Acta Crystallogr D Biol Crystallogr* **2010**, *66* (Pt 2), 125–132.

(2) Yu, F.; Wang, Q.; Li, M.; Zhou, H.; Liu, K.; Zhang, K.; Wang, Z.; Xu, Q.; Xu, C.; Pan, Q.; He, J. Aqua-rium: An Automatic Data-Processing and Experiment Information Management System for Biological Macromolecular Crystallography Beamlines. *J Appl Cryst* **2019**, *52* (2), 472–477.

(3) McCoy, A. J.; Grosse-Kunstleve, R. W.; Adams, P. D.; Winn, M. D.; Storoni, L. C.; Read, R. J. Phaser Crystallographic Software. *J Appl Crystallogr* **2007**, *40* (Pt 4), 658–674.

(4) Adams, P. D.; Grosse-Kunstleve, R. W.; Hung, L. W.; Ioerger, T. R.; McCoy, A. J.; Moriarty, N. W.; Read, R. J.; Sacchettini, J. C.; Sauter, N. K.; Terwilliger, T. C. PHENIX: Building New Software for Automated Crystallographic Structure Determination. *Acta Crystallogr D Biol Crystallogr* **2002**, *58* (Pt 11), 1948–1954.

(5) Emsley, P.; Cowtan, K. Coot: Model-Building Tools for Molecular Graphics. *Acta Crystallogr D Biol Crystallogr* **2004**, *60* (Pt 12 Pt 1), 2126–2132.

(6) Zhao, G.; Peng, W.; Song, K.; Shi, J.; Lu, X.; Wang, B.; Du, Y.-L. Molecular Basis of Enzymatic Nitrogen-Nitrogen Formation by a Family of Zinc-Binding Cupin Enzymes. *Nat Commun* **2021**, *12* (1), 7205.

(7) Richter, M.; Marquetand, P.; González-Vázquez, J.; Sola, I.; González, L. SHARC: Ab Initio Molecular Dynamics with Surface Hopping in the Adiabatic Representation Including Arbitrary Couplings. *J Chem Theory Comput* **2011**, *7* (5), 1253–1258.

(8) Li, P.; Merz, K. M. MCPB.Py: A Python Based Metal Center Parameter Builder. *J Chem Inf Model* **2016**, *56* (4), 599–604.

(9) Trott, O.; Olson, A. J. AutoDock Vina: Improving the Speed and Accuracy of Docking with a New Scoring Function, Efficient Optimization, and Multithreading. *J Comput Chem* **2010**, *31* (2), 455–461.

(10) Pettersen, E. F.; Goddard, T. D.; Huang, C. C.; Couch, G. S.; Greenblatt, D. M.; Meng, E. C.; Ferrin, T. E. UCSF Chimera--a Visualization System for Exploratory Research and Analysis. *J Comput Chem* **2004**, *25* (13), 1605–1612.

(11) Maier, J. A.; Martinez, C.; Kasavajhala, K.; Wickstrom, L.; Hauser, K. E.; Simmerling, C. ff14SB: Improving the Accuracy of Protein Side Chain and Backbone Parameters from ff99SB. *J Chem Theory Comput* **2015**, *11* (8), 3696–3713.

(12) Wang, J.; Wolf, R. M.; Caldwell, J. W.; Kollman, P. A.; Case, D. A. Development and Testing of a General Amber Force Field. *J Comput Chem* **2004**, *25* (9), 1157–1174.

(13) Bayly, C. I.; Cieplak, P.; Cornell, W.; Kollman, P. A. A Well-Behaved Electrostatic Potential Based Method Using Charge Restraints for Deriving Atomic Charges: The RESP Model. *J. Phys. Chem.* **1993**, *97* (40), 10269–10280.

(14) Becke, A. D. Density-functional Thermochemistry. I. The Effect of the Exchange-only Gradient Correction. *J. Chem. Phys.* **1992**, *96* (3), 2155–2160.

(15) Becke, A. D. Density-functional Thermochemistry. III. The Role of Exact Exchange. *J. Chem. Phys.* **1993**, *98* (7), 5648–5652.

(16) Lee, C.; Yang, W.; Parr, R. G. Development of the Colle-Salvetti Correlation-Energy Formula into a Functional of the Electron Density. *Phys Rev B Condens Matter* **1988**, *37* (2), 785–789.

(17) Becke, A. D. Density-functional Thermochemistry. II. The Effect of the Perdew–Wang Generalized-gradient Correlation Correction. *J. Chem. Phys.* **1992**, *97* (12), 9173–9177.

(18) Weigend, F.; Ahlrichs, R. Balanced Basis Sets of Split Valence, Triple Zeta Valence and Quadruple Zeta Valence Quality for H to Rn: Design and Assessment of Accuracy. *Phys Chem Chem Phys* **2005**, *7* (18), 3297–3305.
(19) Jorgensen, W. L.; Chandrasekhar, J.; Madura, J. D.; Impey, R. W.; Klein, M. L. Comparison of Simple Potential Functions for Simulating Liquid Water. *J. Chem. Phys.* **1983**, *79* (2), 926–935.
(20) Brooks, C. L. Computer Simulation of Liquids. *J Solution Chem* **1989**, *18* (1), 99–99.
(21) van Gunsteren, W. F.; Berendsen, H. J. C. Computer Simulation of Molecular Dynamics: Methodology, Applications, and Perspectives in Chemistry. *Angewandte Chemie International Edition in English* **1990**, *29* (9), 992–1023.
(22) Numerical Integration of the Cartesian Equations of Motion of a System with Constraints: Molecular Dynamics of n-Alkanes. *Journal of Computational Physics* **1977**, *23* (3), 327–341.
(23) Darden, T.; York, D.; Pedersen, L. Particle Mesh Ewald: An N·log(N) Method for Ewald Sums in Large Systems. *J. Chem. Phys.* **1993**, *98* (12), 10089–10092.
(24) Marenich, A. V.; Cramer, C. J.; Truhlar, D. G. Universal Solvation Model Based on Solute Electron Density and on a Continuum Model of the Solvent Defined by the Bulk Dielectric Constant and Atomic Surface Tensions. *J Phys Chem B* **2009**, *113* (18), 6378–6396.
(25) Metz, S.; Kästner, J.; Sokol, A. A.; Keal, T. W.; Sherwood, P. ChemShell—a Modular Software Package for QM/MM Simulations. *WIREs Computational Molecular Science* **2014**, *4* (2), 101–110.
(26) Furche, F.; Ahlrichs, R.; Hättig, C.; Klopper, W.; Sierka, M.; Weigend, F. Turbomole. *WIREs Computational Molecular Science* **2014**, *4* (2), 91–100.
(27) Smith, W.; Yong, C. W.; Rodger, P. M. DL_POLY: Application to Molecular Simulation. *Molecular Simulation* **2002**.
(28) Bakowies, D.; Thiel, W. Hybrid Models for Combined Quantum Mechanical and Molecular Mechanical Approaches. *J. Phys. Chem.* **1996**, *100* (25), 10580–10594.
(29) Kästner, J.; Carr, J. M.; Keal, T. W.; Thiel, W.; Wander, A.; Sherwood, P. DL-FIND: An Open-Source Geometry Optimizer for Atomistic Simulations. *J Phys Chem A* **2009**, *113* (43), 11856–11865.
(30) Grimme, S.; Antony, J.; Ehrlich, S.; Krieg, H. A Consistent and Accurate Ab Initio Parametrization of Density Functional Dispersion Correction (DFT-D) for the 94 Elements H-Pu. *J Chem Phys* **2010**, *132* (15), 154104.
(31) Grimme, S.; Ehrlich, S.; Goerigk, L. Effect of the Damping Function in Dispersion Corrected Density Functional Theory. *J Comput Chem* **2011**, *32* (7), 1456–1465.
(32) Zallot, R.; Oberg, N.; Gerlt, J. A. The EFI Web Resource for Genomic Enzymology Tools: Leveraging Protein, Genome, and Metagenome Databases to Discover Novel Enzymes and Metabolic Pathways. *Biochemistry* **2019**, *58* (41), 4169–4182.
(33) Eddy, S. R. Accelerated Profile HMM Searches. *PLoS Comput Biol* **2011**, *7* (10), e1002195.
(34) Fu, L.; Niu, B.; Zhu, Z.; Wu, S.; Li, W. CD-HIT: Accelerated for Clustering the next-Generation Sequencing Data. *Bioinformatics* **2012**, *28* (23), 3150–3152.
(35) Crepin, T.; Schmitt, E.; Mechulam, Y.; Sampson, P. B.; Vaughan, M. D.; Honek, J. F.; Blanquet, S. Use of Analogues of Methionine and Methionyl Adenylate to Sample Conformational Changes during Catalysis in Escherichia Coli Methionyl-tRNA Synthetase. *J Mol Biol* **2003**, *332* (1), 59–72.
(36) Abramson, J.; Adler, J.; Dunger, J.; Evans, R.; Green, T.; Pritzel, A.; Ronneberger, O.; Willmore,

L.; Ballard, A. J.; Bambrick, J.; Bodenstein, S. W.; Evans, D. A.; Hung, C.-C.; O'Neill, M.; Reiman, D.; Tunyasuvunakool, K.; Wu, Z.; Žemgulytė, A.; Arvaniti, E.; Beattie, C.; Bertolli, O.; Bridgland, A.; Cherepanov, A.; Congreve, M.; Cowen-Rivers, A. I.; Cowie, A.; Figurnov, M.; Fuchs, F. B.; Gladman, H.; Jain, R.; Khan, Y. A.; Low, C. M. R.; Perlin, K.; Potapenko, A.; Savy, P.; Singh, S.; Stecula, A.; Thillaisundaram, A.; Tong, C.; Yakneen, S.; Zhong, E. D.; Zielinski, M.; Žídek, A.; Bapst, V.; Kohli, P.; Jaderberg, M.; Hassabis, D.; Jumper, J. M. Accurate Structure Prediction of Biomolecular Interactions with AlphaFold 3. *Nature* **2024**, *630* (8016), 493–500.

(37) Maruyama, C.; Niikura, H.; Takakuwa, M.; Katano, H.; Hamano, Y. Colorimetric Detection of the Adenylation Activity in Nonribosomal Peptide Synthetases. *Methods Mol Biol* **2016**, *1401*, 77–84.

(38) Zhang, M.; Zhang, P.; Xu, G.; Zhou, W.; Gao, Y.; Gong, R.; Cai, Y.-S.; Cong, H.; Deng, Z.; Price, N. P. J.; Mao, X.; Chen, W. Comparative Investigation into Formycin A and Pyrazofurin A Biosynthesis Reveals Branch Pathways for the Construction of C-Nucleoside Scaffolds. *Appl Environ Microbiol* **2020**, *86* (2), e01971-19.

(39) Ren, D.; Wang, S.-A.; Ko, Y.; Geng, Y.; Ogasawara, Y.; Liu, H.-W. Identification of the C-Glycoside Synthases during Biosynthesis of the Pyrazole-C-Nucleosides Formycin and Pyrazofurin. *Angew Chem Int Ed Engl* **2019**, *58* (46), 16512–16516.

(40) Li, W.; Cheng, Z.; Zhao, Z.; Li, H.; Liu, Y.; Lu, X.; Zhao, G.; Du, Y.-L. Discovery of a Bacterial Hydrazine Transferase That Constructs the N-Aminolactam Pharmacophore in Albofungin Biosynthesis. *J Am Chem Soc* **2024**, *146* (19), 13399–13405.

(41) Shikai, Y.; Muramatsu, H.; Igarashi, M.; Katsuyama, Y.; Ohnishi, Y. Identification of a L-Threonine-Utilizing Hydrazine Synthetase for Thrazarine Biosynthesis in Streptomyces Coerulescens MH802-fF5. *Chembiochem* **2025**, e2500298.
